# Updated Variant Curation Expert Panel Criteria and Pathogenicity Classifications for 251 Variants for *RYR1*-related Malignant Hyperthermia Susceptibility

**DOI:** 10.1101/2022.06.23.497341

**Authors:** Jennifer J. Johnston, Robert T. Dirksen, Thierry Girard, Phil M. Hopkins, Natalia Kraeva, Mungunsukh Ognoon, K. Bailey Radenbaugh, Sheila Riazi, Rachel L. Robinson, Louis A. Saddic, Nyamkhishig Sambuughin, Richa Saxena, Sarah Shepherd, Kathryn Stowell, James Weber, Seeley Yoo, Henry Rosenberg, Leslie G. Biesecker

## Abstract

The ClinGen malignant hyperthermia susceptibility (MHS) variant curation expert panel specified the ACMG/AMP criteria for *RYR1*-related MHS and a pilot analysis of 84 variants was published. We have now classified an additional 251 variants for *RYR1-*related MHS according to current ClinGen standards and updated the criteria where necessary. Criterion PS4 was modified such that individuals with multiple *RYR1* variants classified as pathogenic (P), likely pathogenic (LP) or variant of uncertain significance (VUS) were not considered as providing evidence for pathogenicity. Critera PS1 and PM5 were revised to consider LP variants at the same amino acid residue as providing evidence for pathogenicity at reduced strength. Finally, PM1 was revised such that if PS1 or PM5 are used PM1, if applicable, should be downgraded to supporting. Of the 251 *RYR1* variants, 42 were classified as P/LP, 16 as B/LB, and 193 as VUS. The primary driver of 176 VUS classifications was insufficient evidence supporting pathogenicity, rather than evidence against pathogenicity. Functional data supporting PS3/BS3 was identified for only 13 variants. Based on the posterior probabilities of pathogenicity and variant frequencies in gnomAD, we estimated the prevalence of individuals with *RYR1*-related MHS pathogenic variants to be between 1/300 and 1/1,075, considerably higher than current estimates. We have updated ACMG/AMP criteria for *RYR1-*related MHS and classified 251 variants. We suggest that prioritization of functional studies is needed to resolve the large number of VUS classifications and allow for appropriate risk assessment. *RYR1*-related MHS pathogenic variants are likely to be more common than currently appreciated.

## INTRODUCTION

Malignant hyperthermia (MH) is a serious perioperative complication triggered by the use of inhaled anesthetics or succinylcholine (1). Susceptibility to MH (MHS) predominantly demonstrates autosomal dominant inheritance, although not all reported cases are consistent with simple autosomal dominant inheritance (2, 3). About three fourths of MH susceptible families have pathogenic variants in *RYR1* (4), and for individuals known to harbor pathogenic *RYR1* variants, the risk for an MH event can be eliminated by using non-triggering anesthetics. The molecular diagnosis of MHS can be challenging because of the large size of the *RYR1* gene and often limited data available on its variants. Robust and thorough classification of *RYR1* variants would allow pre-screening of surgical patients to enable appropriate clinical management based on genotype (5). As a ClinGen variant curation expert panel (VCEP), we previously specified the American College of Medical Genetics and Genomics/Association of Molecular Pathologists (ACMG/AMP) criteria for pathogenicity for the dyadic disease entity (6) of *RYR1*-related malignant hyperthermia susceptibility (MHS). The specified criteria were piloted on a set of 84 variants and here we expand that work classifying an additional 251 variants. We then used these variant classifications and a large population genomic database to estimate the prevalence of individuals with pathogenic MHS variants, which has been challenging to determine (1, 7).

## Results

### Classification of RYR1 variants

For this study, 251 *RYR1* variants were classified for pathogenicity for *RYR1*-related MHS, two were classified as pathogenic, 40 were classified as likely pathogenic, 193 were classified as VUS, 13 were classified as likely benign, and three were classified as benign (Figure 1A). Variant classifications and ACMG/AMP criteria are presented in Supplementary Material, Table S1. The VCEP previously classified 84 variants, and when those classifications are combined with those in this work, this provides 335 classified variants (Figure 1B, Supplementary Material, Table S2). For the complete set of variants, 29 are now classified as pathogenic, 57 are now classified as likely pathogenic, 219 are now classified as VUS, 17 are now classified as likely benign, and 13 are now classified as benign. Two variants classified as pathogenic in our prior work (7) were downgraded to likely pathogenic here, based on evidence discussed during the curation of the additional 251 variants. For NM_000540.3:c.11315G>A, p.(Arg3772Gln), BS2_Mod was applied for a genotype positive/phenotype negative individual newly identified in the literature (4). For NM_000540.3:c.7354C>T, p.(Arg2452Trp), we determined that identical segregation data were published multiple times in the literature and double counted in our prior variant biocuration, to correct for this, PP1 was downgraded from strong to moderate for this variant.

**Figure 1.**
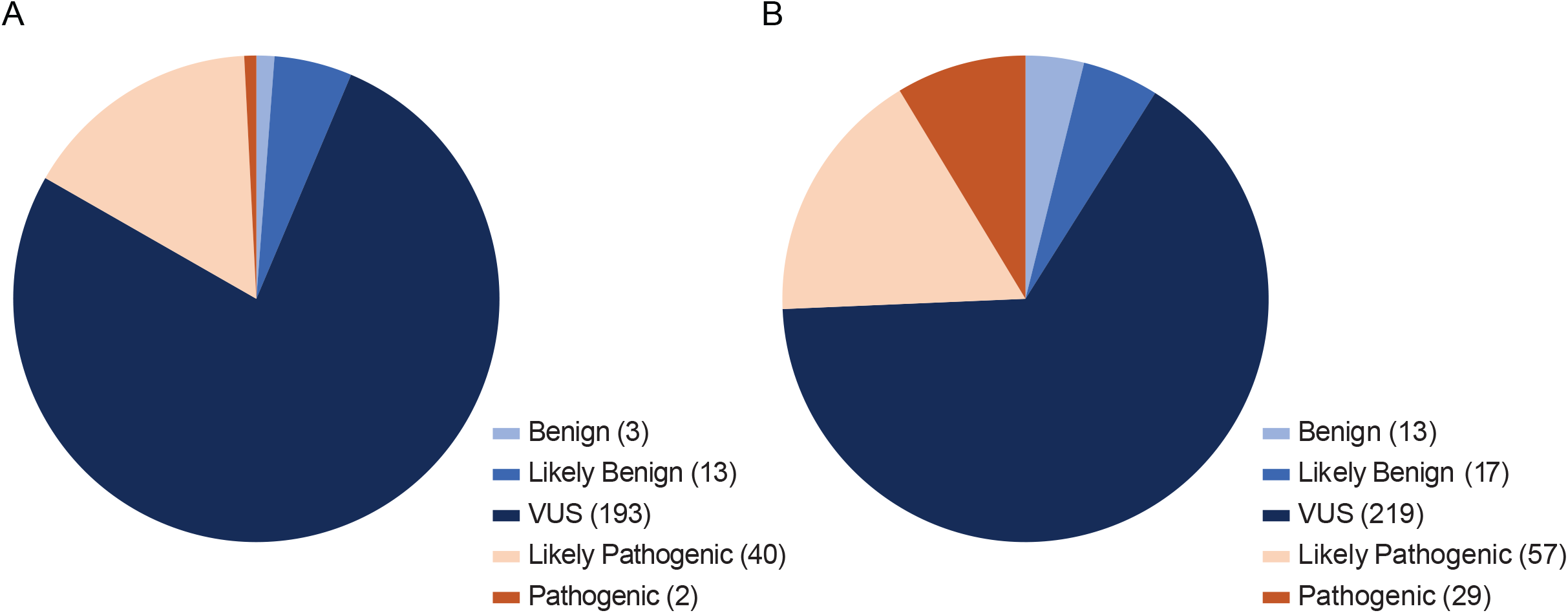
A) Variant pathogenicity classification distribution for the 251 variants classified in this study. B) Distribution for all 335 variants classified to date by the ClinGen *RYR1*-MHS VCEP.

While it is axiomatic that pathogenic criteria would be found more often in P/LP variants and benign criteria would be found more often in B/LB variants, it is useful to tabulate which criteria contributed the most frequently to these classifications. Variants classified as P/LP were more likely as compared to variants classified as B/LB/VUS to have been reported in individuals who met *RYR1*-specified criteria for PS4, providing case evidence (84/86 versus 113/239, p<0.0001) and segregation data (PP1, 59/86 versus 11/239, p<0.0001). (Eleven variants assigned BA1 were not fully assessed and are not included in these analyses.) Other criteria that were more often associated with P/LP variants as compared to B/LB/VUS variants included presence in a hotspot (PM1, 72/86 versus 117/239, p<0.0001) and a bioinformatic prediction of pathogenicity (PP3, REVEL ≥0.85, 73/86 versus 94/239, p<0.0001.). Criteria more often associated with B/LB variants as compared to P/LP variants included high minor allele frequency in gnomAD (BS1, popmax >0.0008, 10/19 versus 0/86, p<0.0001.) and presence in individuals who did not demonstrate sensitivity to RYR1 agonists on contracture testing (BS2, 13/19 versus 13/86, p<0.0001), Table 1.

**Table 1.** ACMG criteria assigned to 335 *RYR1*-related MHS variants. For each criterion, all strength levels are combined in a single column. Individual strength levels for each criterion are presented in Table S4.

|  | **PS1**<br>**(3)** | **PS2/PM6**<br>**(7)** | **PS3**<br>**(51)** | **PS4**<br>**(199)** | **PM1**<br>**(188)** | **PM5**<br>**(25)** | **PP1**<br>**(69)** | **PP3**<br>**(166)** | **BA1**<br>**(11)** | **BS1**<br>**(10)** | **BS2**<br>**(27)** | **BS3**<br>**(8)** | **BP2**<br>**(2)** | **BP4**<br>**(21)** |
| --- | --- | --- | --- | --- | --- | --- | --- | --- | --- | --- | --- | --- | --- | --- |
| Pathogenic<br>(29) | 2 | 1 | 27 | 29 | 26 | 6 | 29 | 27 | 0 | 0 | 9 | 0 | 0 | 0 |
| Likely<br>Pathogenic<br>(57) | 1 | 4 | 19 | 55 | 46 | 11 | 30 | 46 | 0 | 0 | 4 | 0 | 0 | 0 |
| VUS (219) | 0 | 2 | 3 | 110 | 112 | 9 | 10 | 91 | 0 | 0 | 2 | 4 | 1 | 18 |
| Likely<br>Benign (17) | 0 | 0 | 1 | 3 | 4 | 0 | 1 | 3 | 0 | 8 | 11 | 3 | 0 | 2 |
| Benign (13) | 0 | 0 | 0 | 0 | 1 | 0 | 0 | 0 | 11 | 2 | 2 | 1 | 1 | 1 |

During classification of the additional 251 *RYR1* variants, several changes were made to the ACMG/AMP rule specifications for *RYR1*-related MHS. For PS4 (prevalence of variant in affected individuals versus controls) we determined that only individuals with a single VUS/LP/P variant in *RYR1* would be considered. Our reasoning was that it was difficult to attribute contribution to MHS for a variant when it was present in a case with more than one possibly pathogenic variant. For PS1 (same amino acid change, different nucleotide change) and PM5 (different amino acid change, same codon) it was determined that a variant previously classified as likely pathogenic without the use of PS1 or PM5 could be used as supporting evidence so long as the Grantham score (8) difference of the new variant compared to reference was greater than that for the previously identified likely pathogenic variant compared to reference. Furthermore, it was determined that criteria PS1 and PM5 address evidence related to PM1 (location in a hotspot). To avoid overweighting, when either PS1 or PM5 is used, if PM1 is applicable it should be implemented at supporting strength, rather than the default of moderate. One variant classified as pathogenic in our prior work was downgraded to likely pathogenic based on these changes to the *RYR1*-related MHS criteria. Variant NM_000540.3:c.7372C>T p.(Arg2458Cys), was reclassified as LP using criteria PS3_Mod, PS4_Mod, PM1_Sup, PM5, PP3_Mod.

### Estimate of RYR1-related MHS Pathogenic Variant Prevalence

Altogether, the *RYR1* VCEP assessed 335 *RYR1* variants for pathogenicity with respect to *RYR1*-related MHS using Bayes (9) to combine the ACMG/AMP criteria and calculate posterior probabilities for pathogenicity. Using the posterior probabilities for classified variants and the gnomAD frequencies, one can estimate the prevalence of pathogenic *RYR1*-related MHS variants in the population. Using total gnomAD population frequencies for all LB/VUS/LP/P classified variants, we estimated a prevalence of pathogenic *RYR1*-related MHS variants of 1/300 individuals. In the five gnomAD continental populations the calculated prevalence ranged from 1/240 for the Non-Finnish European (NFE) population to 1/546 for the South Asian (SAS) population (Table 2). When only considering variants classified as P/LP the estimated prevalence of pathogenic *RYR1* MHS-related variants dropped to 1/1,075 for all of gnomAD. In the five gnomAD continental populations the calculated prevalence ranged from 1/802 for the NFE population to 1/5,990 for the AMR population.

**Table 2.**
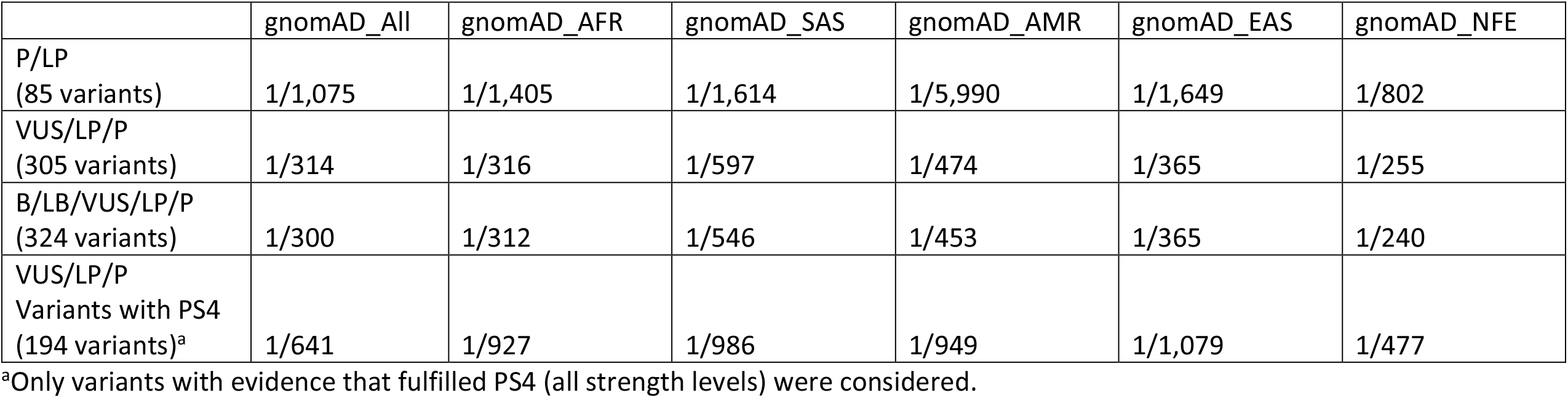
Predicted prevalence of *RYR1*-related MHS associated variants based on posterior probabilities of pathogenicity and frequencies in gnomAD for 324 classified variants.

## Discussion

The ClinGen *RYR1* VCEP has now classified a total of 335 *RYR1* variants from variant databases and the literature for pathogenicity for *RYR1*-related malignant hyperthermia susceptibility. These variant classifications were made available through ClinVar. The availability of these classifications should ease the interpretation burden for clinical laboratories for reporting diagnostic results. It should also facilitate the interpretation of variants for *RYR1*-related MHS as a secondary finding, as this entity has been included in the ACMG secondary findings recommendations since its inception (10-12).

This variant classification work across a large number of reported variants allowed our VCEP to refine and adjust our previously determined criteria. Based on this work, criterion PS4 (prevalence of variant in affected individuals versus controls) was modified to only consider individuals with a single VUS/LP/P variant in *RYR1*. In the *RYR1* specified criteria PS4 is implemented by counting MH cases with the *RYR1* variant under consideration. When an individual harbors multiple *RYR1* variants the probability that any one variant is causative of MH decreases, and thus, it was determined that these cases should not be considered for PS4. The PS1 (same amino acid change, different nucleotide change) and PM5 (different amino acid change, same codon) criteria were modified such that a variant previously classified as likely pathogenic could be used at a strength level of supporting, the previously classified variant must have been classified as likely pathogenic without the use of PS1 or PM5. Furthermore, it was determined that when PS1 or PM5 are used, if PM1 applies it should be reduced to a supporting strength level.

As expected, attributes that supported pathogenicity were used more often and at higher strength levels for P/LP variants as compared to VUS/LB/B variants (Table 1, Supplementary Material, Table S3). Specifically, PS4 (all strength levels) was used in 100% of variants that were classified as pathogenic and only 50% of variants that were classified as VUS. Also as expected, the assigned strength level of PS4 correlated with pathogenicity, with 1% of VUS having garnered PS4 at a strong level of evidence compared to 66% of pathogenic variants. Only 18% of likely benign variants garnered PS4, all at a strength of supporting. Similarly, segregation data that contributed to PP1 (all strength levels) were available for 100% of variants classified as pathogenic but only 5% of variants classified as VUS. Functional data that fulfilled PS3 (all strength levels) were available for 93% of pathogenic variants, 33% of likely pathogenic variants, but only 1% of VUS. Conversely BS2 was used for 65% of likely benign variants but only 1% of VUS. Interestingly, BS2 was invoked for 31% of pathogenic variants. As many pathogenic variants have been well studied, with multiple family members having undergone contracture testing, these variants were more likely to have been identified in at least one genotype positive and phenotype negative (contracture testing) individual. This is to be expected for a disorder with less than complete penetrance and it is reassuring to note that the criteria appear to be robust to these observations. The use of PS4 and PP1 in well studied variants balanced the use of BS2. In the Bayes system BS2_Moderate (for one genotype positive individual with a negative contracture test) would be balanced by a single moderate strength criterion. For well-studied pathogenic variants, more likely to have functional data and multiple reported cases, sufficient criteria were often met such that the use of criterion BS2 did not lead to a change in the five-level classification, even though the posterior probability was reduced.

Of the 335 classified *RYR1* variants, more than half, 219, were classified as VUS. This was primarily attributable to the absence of adequate data to support a classification of likely pathogenic or likely benign, rather than a mix of pathogenic and benign evidence. The posterior probability of pathogenicity for a VUS ranges from 0.10 to 0.89 which is challenging for clinicians using variant classifications to make management decisions. To resolve this issue, more data need to be collected to move variants from VUS classifications. ACMG/AMP criteria used to classify variants for *RYR1*-related MHS fit three major categories: 1) criteria intrinsic to the variant (PS1, same amino acid change; PM1, presence in hotspot; PM5, same residue different missense alteration; PP3/BP4, bioinformatic prediction; BA1, too common in gnomAD; BS1, more common than expected for disorder), 2) criteria based on case data (PS2/PM6, *de novo* status; PS4, case numbers; PP1, segregation; BS2, negative phenotype (contracture test) in variant positive individual; BP2, identified in *cis* to known variant), and 3) criteria based on functional data (PS3; BS3). It is axiomatic that criteria intrinsic to the variant do not change over time. Case-based criteria other than PP1 require new clinical observations of MH to be identified to trigger their use or increase their weight. Without the identification of additional cases, contracture testing of variant positive relatives (PP1; BS2) and/or functional analyses of variants (PS3; BS3) are the only actions that can resolve a classification of VUS. While segregation data based on genotype and contracture testing can support or refute pathogenicity, contracture testing is an invasive procedure requiring surgery and is often not feasible. For VUS that are at the higher posterior probability range of pathogenicity (>0.67, without functional data) the addition of pathogenic functional data at a moderate weight would allow reclassification from a VUS to likely pathogenic (67 variants). Similarly, for VUS with a low posterior probability of pathogenicity (<0.18, without functional data) the addition of functional data that supports a benign classification at the level of supporting will allow for reclassification from a VUS to likely benign (50 variants). We conclude that generating functional data should be a high priority for the field as it has the potential to better classify over half of the existing *RYR1-*related MHS VUS.

While the prevalence of MHS has been difficult to determine, the incidence of MH events has been estimated to be between one in 10,000 and one in 250,000 surgeries (1, 13).Furthermore, the combined prevalence of functionally characterized variants associated with MH in the ExAC cohort was reported to be one in 2,750 (14). With the classification of 335 *RYR1* variants reported here we predicted the combined variant frequency of *RYR1*-related MHS pathogenic variants to be one in 599 (MAF 0.0017) using individual variant frequencies in gnomAD and posterior probabilities of pathogenicity (Supplementary Material, Tables S4 and S5). This would predict the prevalence of MH-associated *RYR1* pathogenic variants to be about one in 300 individuals. This value is approximately ten times higher than prior estimates for MHS pathogenic variants in the population (13) and 33 to 833 times the recognized incidence of MH during surgery. *RYR1*-related MH is known to demonstrate reduced penetrance such that the frequency of pathogenic variants in the population is expected to be higher than the incidence of MH events, however, reduced penetrance may not entirely account for the discrepancy. One limitation of this approach is the accurate assessment of posterior probability, which requires both the prior probability to be set correctly, currently set at 0.1, and the accurate identification and weighting of all factors affecting pathogenicity. Toward that end, 34 variants had no information supporting or refuting pathogenicity and therefore their posterior probability remained at 0.1. These 34 variants accounted for 13% of the predicted population pathogenic variant frequency (0.00021). Considering all VUS variants, 67% of the pathogenic variant burden was accounted for. Using only the P/LP variants, we calculated an overall variant frequency of 0.00047 or a prevalence of *RYR1-*related MH associated pathogenic variants of one in 1,075 individuals. We predict that the calculated prevalence using only the P/LP variants is an underestimate of the true prevalence and the estimate that includes all variants is an overestimate. Therefore, the true prevalence of *RYR1*-related MHS pathogenic variants likely lies between one in 300 and one in 1,075. A further complication for these estimates is the well-recognized phenomenon of bias in genetics and genomics research towards Western European populations. The great majority of MHS research has been done in North America Australasia and Europe, with some work done in Japan. Relatively little has been done in the rest of East Asia, South Asia, Africa, or the Middle East. This dearth of research may explain the lower prevalence that we estimate for those populations (Table 2) as our method relies heavily on prior identification of pathogenic variants in populations. In spite of these limitations, these data suggest that the prevalence of *RYR1*-related MHS pathogenic variants is higher than the previous estimate of one in 2,000 to one in 3,000 (2). This, in turn, suggests that the penetrance of the trait is lower than is commonly presumed.

These data have broad implications for MH, MHS research, and anesthesia practice. First, it is clear that additional evidence needs to be generated to better classify the many VUS in this gene. We believe that the most practical approach is to generate high throughput functional data, which we estimate could shift nearly half of variants from VUS to either P/LP or B/LB. Another priority should be genomic assessment of individuals with MH from populations that are currently understudied, as has been emphasized for many other categories of genetic disease. The implications are that *RYR1*-mediated genetic susceptibility to MHS is not rare. If our estimates of prevalence are correct, more than 300,000 individuals in the US harbor a pathogenic *RYR1* variant for MHS, which does not meet the NIH Office of Rare Disease Research definition of rare disease. At the same time, recognized MH reactions are indeed rare – probably in the range of 1/100,000 intraoperative exposures. As well, it is likely that some MH reactions are subclinical. Together these data suggest that the penetrance of MHS is lower than currently thought and that the expressivity may be more variable than is currently recognized. Further research to understand the genetic architecture of MHS is essential to refine these risks.

## Materials and Methods

### Variant Classification

The ClinGen MHS VCEP set out to classify the pathogenicity of variants for *RYR1*-related MHS, also known by its OMIM descriptors of ‘susceptibility to malignant hyperthermia 1’ or ‘MHS1’ (MIM:145600). *RYR1* variants included in this study were either assessed by one or more submitters as pathogenic or likely pathogenic (P/LP) for *RYR1*-related MHS in ClinVar (15), noted as ‘disease mutations’ (DM) in relation to *RYR1*-related MHS in the Human Gene Mutation Database (16), or associated with *RYR1*-related MHS in the literature. Variants were classified using the ACMG/AMP criteria (10, 11) with specifications for *RYR1*-related MHS (7).Relevant criteria were combined using the Bayes model to determine the posterior probability of pathogenicity and the ACMG/AMP-specified five-level classification, pathogenic (P)/likely pathogenic (LP)/variant of uncertain significance (VUS)/likely benign (LB)/benign (B), Table S1 (9, 17).

All variants were biocurated and classified by a primary biocurator and reviewed by a second biocurator. Biocurations and classifications were distributed to the full ClinGen MHS VCEP and discussed at monthly variant review meetings. Classifications were finalized after open discussion with a minimum of three variant scientists required to be present to finalize variant classifications. Final classifications were entered into the ClinGen variant curation interface (18).

RYR1-related MHS variant specifications to the ACMG/AMP criteria were reviewed during this process to determine if revisions were warranted. Revisions were discussed on the VCEP calls and adopted with unanimous approval. (Current specifications can be found at https://www.clinicalgenome.org/affiliation/50038.)

### Population Prevalence

A total of 335 *RYR1* variants (251 in this work, 84 with the original VCEP rule specification) were classified by the MHS VCEP using the ACMG/AMP criteria specified for *RYR1*. An estimate of the burden of *RYR1*-related MHS pathogenic variants in a population was made using the sum of the products of the posterior probabilities of pathogenicity and population variant frequencies.

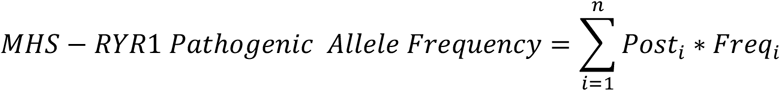

In this equation, “Post” is the posterior probability of pathogenicity of each variant and “Freq” is the population frequency of the variant from the genome aggregation database (gnomAD). Population frequencies from combined exome and genome data in gnomAD were downloaded for 324 classified *RYR1* variants and each variant’s contribution to the overall number of pathogenic variants was determined by multiplying the frequency of each variant with its posterior probability. For example, if a variant had a frequency of 0.0001 with a posterior probability of pathogenicity of 1.0 then this would contribute one pathogenic variant for every 5,000 individuals (pathogenic variant contribution 0.0001). However, if the probability of pathogenicity was reduced to 0.9 this would contribute one pathogenic variant for every 5,555 individuals (pathogenic variant contribution 0.00009). The overall number of pathogenic variants in a population was then calculated by summing all individual variant contributions. We excluded 11 variants classified as benign using BA1 (too common for disorder).

