## Supplemental Tables for "Updated Variant Curation Expert Panel Criteria and Pathogenicity Classifications for 251 Variants for *RYR1*-related Malignant Hyperthermia Susceptibility"

| | |
|---|---|
| Supplemental Table 1 | 251 variant classifications from this work. |
| Supplemental Table 2 | 335 variant classifications, inlcudes 84 variants from prior work PMID: 33767344. |
| Supplemental Table 3 | Usage of ACMG criteria by classification. |
| Supplemental Table 4 | Posterior probabilities and gnomAD frequencies for 335 *RYR1* variants. |
| Supplemental Table 5 | Predictions of RYR1-related MHS pathogenic allele frequency and prevalence in the population based on gnomAD frequencies and posterior probabilities. |

Supplemental Table 1

| Genomic Coordinate (GRCh37) | cDNA NM_000540.2 | Protein | REVEL | ACMG/AMP Criteria | Pathogenicity | Posterior Probability | Amino Acid Position | Publications |
|---|---|---|---|---|---|---|---|---|
| chr19:g.38924507T>G | c.38T>G | p.(Leu13Arg) | 0.828 | PS3_Mod, PS4_Mod, PM1, PP1 | Likely Pathogenic | 0.949 | 13 | PMID:34127251; PMID:19191329; PMID:30236257 ; PMID:16732084; PMID:23558838; PMID:16917943; PMID:23422674; PMID:25960145 |
| chr19:g.38931390_38931392del | c.51_53del | p.(Asp17del) | | PM1 | VUS | 0.325 | 17 | PMID:16732084 |
| chr19:g.38931436A>G | c.97A>G | p.(Lys33Glu) | 0.92 | PS2_PM6_Sup, PS4_Sup, PM1, PP3_Mod | Likely Pathogenic | 0.900 | 33 | PMID:18765655 |
| chr19:g.38931458G>C | c.119G>C | p.(Gly40Ala) | 0.897 | PS4_Sup, PM1, PP3_Mod | VUS | 0.812 | 40 | PMID:23558838 |
| chr19:g.38931470G>A | c.131G>A | p.(Arg44His) | 0.934 | PS4_Sup, PM1, PP3_Mod | VUS | 0.812 | 44 | PMID:30236257 ; PMID:16521288; PMID:16835904; PMID:16917943 |
| chr19:g.38933001G>A | c.178G>A | p.(Asp60Asn) | 0.736 | PS4_Sup, PM1 | VUS | 0.500 | 60 | PMID:30236257 ; PMID:16621918; PMID:16917943; PMID:23422674 |
| chr19:g.38933001G>T | c.178G>T | p.(Asp60Tyr) | 0.961 | PS4_Sup, PM1, PP3_Mod | VUS | 0.812 | 60 | PMID:24433488 |
| chr19:g.38933013T>C | c.190T>C | p.(Cys64Arg) | 0.809 | PM1 | VUS | 0.325 | 64 | PMID:21455645; PMID:23460944 |
| chr19:g.38933035C>A | c.212C>A | p.(Ser71Tyr) | 0.937 | PM1, PP3_Mod | VUS | 0.675 | 71 | PMID:16835904; PMID:16940308; PMID:17365175; PMID:17483490; PMID:19027160; PMID:23919265; PMID:30611313 |
| chr19:g.38933074C>T | c.251C>T | p.(Thr84Met) | 0.735 | PS3_Mod, PM1 | VUS | 0.675 | 84 | PMID:30236257 |
| chr19:g.38934430G>A | c.418G>A | p.(Ala140Thr) | 0.261 | PM1, BS2_Mod, BP4 | Likely Benign | 0.051 | 140 | PMID:21455645 |
| chr19:g.38934819C>A | c.455C>A | p.(Ala152Asp) | 0.9 | PS4_Sup, PM1, PP3_Mod | VUS | 0.812 | 152 | PMID:30236257 |
| chr19:g.38934827C>A | c.463C>A | p.(Gln155Lys) | 0.94 | PS3_Mod, PM1, PP3_Mod | Likely Pathogenic | 0.900 | 155 | PMID:16732084; PMID:16917943; PMID:31841587 |
| chr19:g.38934831G>A | c.467G>A | p.(Arg156Lys) | 0.886 | PM1, PP3_Mod | VUS | 0.675 | 156 | PMID:16835904; PMID:16917943; PMID:19541610 |
| chr19:g.38934843A>G | c.479A>G | p.(Glu160Gly) | 0.957 | PS4_Sup, PM1, PP3_Mod | VUS | 0.812 | 160 | PMID:30236257 ; PMID:14985404; PMID:16917943; PMID:21674524; PMID:19913485; PMID:20566647; PMID:23919265 |
| chr19:g.38934852G>A | c.488G>A | p.(Arg163His) | 0.864 | PS4_Sup, PM1, PP3_Mod | VUS | 0.812 | 163 | PMID:30236257 ; PMID:25658027 |
| chr19:g.38934857G>A | c.493G>A | p.(Gly165Arg) | 0.957 | PS4_Sup, PM1, PP3_Mod | VUS | 0.812 | 165 | PMID:16163667; PMID:16917943 |
| chr19:g.38934860G>A | c.496G>A | p.(Asp166Asn) | 0.829 | PS4_Mod, PM1 | VUS | 0.675 | 166 | PMID:12059893; PMID:16163667; PMID:12434264; PMID:16917943 |
| chr19:g.38934861A>G | c.497A>G | p.(Asp166Gly) | 0.967 | PM1, PP3_Mod | VUS | 0.675 | 166 | PMID:16732084; PMID:16917943 |
| chr19:g.38934890G>A | c.526G>A | p.(Glu176Lys) | 0.803 | PS4_Mod, PM1 | VUS | 0.675 | 176 | PMID:30236257 |
| chr19:g.38934892G>T | c.528G>T | p.(Glu176Asp) | 0.504 | PS4_Mod, PM1 | VUS | 0.675 | 176 | PMID:Kaulins 2008; PMID:29635721 |
| chr19:g.38934893C>T | c.529C>T | p.(Arg177Cys) | 0.931 | PS2_PM6_Mod, PS4, PM1, PP1_St, PP3_Mod, BS2_Mod | Pathogenic | 0.999 | 177 | PMID:30236257 ; PMID:16163667; PMID:18564801; PMID:16521288; PMID:16835904; PMID:16917943; PMID:19541610; PMID:19648156; PMID:20566647; PMID:25658027; PMID:28078069 |
| chr19:g.38934897A>C | c.533A>C | p.(Tyr178Ser) | 0.901 | PS4_Sup, PM1, PP3_Mod | VUS | 0.812 | 178 | PMID:30236257 |
| chr19:g.38934897A>G | c.533A>G | p.(Tyr178Cys) | 0.951 | PS2_PM6_Mod, PS4_Sup, PM1, PP1, PP3_Mod | Likely Pathogenic | 0.975 | 178 | PMID:16163667; PMID:16917943 |
| chr19:g.38935311G>A | c.625G>A | p.(Glu209Lys) | 0.711 | PM1, BP2 | VUS | 0.188 | 209 | PMID:21455645 |
| chr19:g.38937121C>T | c.641C>T | p.(Thr214Met) | 0.533 | PS4_Mod, PM1, PP1, BS2_Mod, BS3_Sup | VUS | 0.325 | 214 | PMID:30236257 ; PMID:27857962; PMID:25658027; PMID:33333461 |
| chr19:g.38937132G>A | c.652G>A | p.(Val218Ile) | 0.801 | PS4_Sup, PM1 | VUS | 0.500 | 218 | PMID:30236257 ; PMID:16732084; PMID:16917943; PMID:23422674 |
| chr19:g.38937157T>A | c.677T>A | p.(Met226Lys) | 0.841 | PS4_Mod, PM1 | VUS | 0.675 | 226 | PMID:19191329; PMID:24433488; PMID:30236257 ; PMID:16917943; PMID:18564801; PMID:21795085; PMID:30325262 |
| chr19:g.38937160A>T | c.680A>T | p.(Asp227Val) | 0.933 | PS4_Sup, PM1, PP3_Mod | VUS | 0.812 | 227 | PMID:16163667; PMID:16917943; PMID:30325262 |
| chr19:g.38939140C>T | c.946C>T | p.(Arg316Cys) | 0.793 | PM1 | VUS | 0.325 | 316 | PMID:31304636 |
| chr19:g.38939141G>T | c.947G>T | p.(Arg316Leu) | 0.928 | PM1, PP3_Mod | VUS | 0.675 | 316 | PMID:16732084; PMID:16917943 |
| chr19:g.38939320_38939321insGGA | c.992_994dup | p.(Glu331dup) | | PS4_Sup, PM1 | VUS | 0.500 | 331 | |
| chr19:g.38939355G>A | c.1024G>A | p.(Glu342Lys) | 0.927 | PS4_Sup, PM1, PP3_Mod | VUS | 0.812 | 342 | PMID:24433488 |
| chr19:g.38939431G>A | c.1100G>A | p.(Arg367Gln) | 0.651 | PS4_Mod, PM1 | VUS | 0.675 | 367 | PMID:21965348 |
| chr19:g.38939431G>T | c.1100G>T | p.(Arg367Leu) | 0.866 | PS4_Sup, PM1, PP3_Mod | VUS | 0.812 | 367 | PMID:24433488; PMID:16917943; PMID:19191329; PMID:21795085 |
| chr19:g.38942425C>A | c.1144C>A | p.(His382Asn) | 0.739 | PS4_Sup, PM1 | VUS | 0.500 | 382 | PMID:19346234; PMID:20142353 |
| chr19:g.38942482C>A | c.1201C>A | p.(Arg401Ser) | 0.839 | PS4_Sup, PM1_Sup, PM5 | VUS | 0.675 | 401 | PMID:16163667 |
| chr19:g.38942482C>G | c.1201C>G | p.(Arg401Gly) | 0.865 | PM1_Sup, PM5, PP3_Mod | VUS | 0.812 | 401 | PMID:16917943; PMID:23422674 |
| chr19:g.38942483G>A | c.1202G>A | p.(Arg401His) | 0.903 | PS3_Mod, PS4, PM1, PP1, PP3_Mod | Pathogenic | 0.997 | 401 | PMID:12059893; PMID:16163667; PMID:30236257 ; PMID:23460944; PMID:24433488; PMID:12434264; PMID:16917943; PMID:26115329; PMID:31841587 |

| | | | | | | | | |
|---|---|---|---|---|---|---|---|---|
| chr19:g.38942483G>T | c.1202G>T | p.(Arg401Leu) | 0.965 | PS4_Mod, PM1_Sup, PM5, PP1, PP3_Mod | Likely Pathogenic | 0.975 | 401 | PMID:30236257 |
| chr19:g.38943625C>T | c.1411C>T | p.(Arg471Cys) | 0.578 | PM1 | VUS | 0.325 | 471 | PMID:1354642 |
| chr19:g.38943636G>T | c.1422G>T | p.(Gln474His) | 0.835 | PM1 | VUS | 0.325 | 474 | PMID:16732084; PMID:16621918; PMID:16917943 |
| chr19:g.38945893C>G | c.1459C>G | p.(Leu487Val) | 0.668 | PS4_Sup, PM1 | VUS | 0.500 | 487 | PMID:30236257 |
| chr19:g.38945894T>C | c.1460T>C | p.(Leu487Pro) | 0.962 | PS4_Sup, PM1, PP3_Mod | VUS | 0.812 | 487 | PMID:23558838 |
| chr19:g.38945909G>A | c.1475G>A | p.(Arg492His) | 0.81 | PM1 | VUS | 0.325 | 492 | PMID:30236257 ; PMID:25658027; PMID:32054689 |
| chr19:g.38945987T>C | c.1553T>C | p.(Val518Ala) | 0.738 | PM1 | VUS | 0.325 | 518 | PMID:23558838 |
| chr19:g.38945999A>G | c.1565A>G | p.(Tyr522Cys) | 0.949 | PS4_Mod, PM1_Sup, PM5, PP3_Mod | Likely Pathogenic | 0.949 | 522 | PMID:30236257 ; PMID:32919876; PMID:16244001; PMID:16917943; PMID:19762757; PMID:21965348; PMID:26115329; PMID:31841587 |
| chr19:g.38946129T>C | c.1615T>C | p.(Phe539Leu) | 0.934 | PS4_Mod, PM1, PP1, PP3_Mod | Likely Pathogenic | 0.949 | 539 | PMID:18564801; PMID:30236257 |
| chr19:g.38946129T>G | c.1615T>G | p.(Phe539Val) | 0.937 | PS4_Mod, PM1_Sup, PM5_Sup, PP3_Mod | Likely Pathogenic | 0.900 | 539 | PMID:30236257 |
| chr19:g.38946144G>T | c.1630G>T | p.(Asp544Tyr) | 0.924 | PS4_Sup, PM1, PP1, PP3_Mod | Likely Pathogenic | 0.900 | 544 | PMID:19191329; PMID:21795085; PMID:22473935; PMID:28259615 |
| chr19:g.38948179G>C | c.1834G>C | p.(Ala612Pro) | 0.971 | PS4_Sup, PP1, PP3_Mod | VUS | 0.675 | 612 | PMID:22415532 |
| chr19:g.38948815G>C | c.2050G>C | p.(Gly684Arg) | 0.962 | PS4_Sup, PP3_Mod | VUS | 0.500 | 684 | PMID:30236257 |
| chr19:g.38951101C>T | c.2447C>T | p.(Pro816Leu) | 0.758 | PS4_Sup, BS2_Mod, BS3_Sup | Likely Benign | 0.025 | 816 | PMID:30236257 ; PMID:30864471; PMID:31903994 |
| chr19:g.38951191C>T | c.2537C>T | p.(Ser846Leu) | 0.558 | PS4_Sup | VUS | 0.188 | 846 | PMID:30236257 ; PMID:16917943 |
| chr19:g.38954139G>A | c.2654G>A | p.(Arg885His) | 0.609 | None | VUS | 0.100 | 885 | PMID:30236257 ; PMID:25960145; PMID:31395899 |
| chr19:g.38956784G>A | c.2924G>A | p.(Arg975Gln) | 0.329 | PS4_Sup, BP4 | VUS | 0.100 | 975 | PMID:30236257 |
| chr19:g.38956955G>A | c.3095G>A | p.(Arg1032His) | 0.547 | None | VUS | 0.100 | 1032 | PMID:30236257 ; PMID:24627108 |
| chr19:g.38956987C>T | c.3127C>T | p.(Arg1043Cys) | 0.936 | PP3_Mod | VUS | 0.325 | 1043 | PMID:16917943; PMID:23204524; PMID:23558838; PMID:26245150; PMID:29298851; PMID:30155738 ; PMID:33646171; PMID:19191329 |
| chr19:g.38957026G>A | c.3166G>A | p.(Asp1056Asn) | 0.685 | PS4_Sup | VUS | 0.188 | 1056 | PMID:20681998 |
| chr19:g.38957026G>C | c.3166G>C | p.(Asp1056His) | 0.824 | PS4_Mod, PP1_St | Likely Pathogenic | 0.900 | 1056 | PMID:30236257 ; PMID:24013571 |
| chr19:g.38957032G>A | c.3172G>A | p.(Glu1058Lys) | 0.78 | PS4_Mod | VUS | 0.325 | 1058 | PMID:19346234; PMID:30236257 ; PMID:20142353; PMID:25989378 |
| chr19:g.38958295G>A | c.3224G>A | p.(Arg1075Gln) | 0.846 | PS4_Sup, BS2_Mod | Likely Benign | 0.051 | 1075 | PMID:30236257 ; PMID:19454545; PMID:28818389 |
| chr19:g.38959642C>T | c.3418C>T | p.(Arg1140Cys) | 0.807 | None | VUS | 0.100 | 1140 | PMID:30236257 ; PMID:16917943; PMID:25370123 |
| chr19:g.38959751C>T | c.3527C>T | p.(Thr1176Ile) | 0.162 | BP4 | VUS | 0.100 | 1176 | PMID:30236257 |
| chr19:g.38960044A>C | c.3656A>C | p.(Gln1219Pro) | 0.891 | PP3_Mod | VUS | 0.325 | 1219 | PMID:21965348 |
| chr19:g.38960055G>A | c.3667G>A | p.(Glu1223Lys) | 0.831 | None | VUS | 0.100 | 1223 | PMID:30236257 |
| chr19:g.38965975A>G | c.4178A>G | p.(Lys1393Arg) | 0.555 | BA1 | Benign | <0.001 | 1393 | PMID:19346234; PMID:24433488; PMID:25658027; PMID:25735680; PMID:25960145; PMID:20142353; PMID:22473935; PMID:23329375; PMID:23628358; PMID:24195946; PMID:25637381; PMID:25985138; PMID:25989378; PMID:26332594; PMID:26565425; PMID:26972305; PMID:27153395; PMID:30788618; PMID:25614869 |
| chr19:g.38973969C>T | c.4747C>T | p.(Arg1583Cys) | 0.586 | None | VUS | 0.100 | 1583 | PMID:30864471; PMID:19346234; PMID:23035052; PMID:25989378 |
| chr19:g.38973985C>T | c.4763C>T | p.(Pro1588Leu) | 0.735 | None | VUS | 0.100 | 1588 | PMID:30236257 |
| chr19:g.38973997C>T | c.4775C>T | p.(Pro1592Leu) | 0.927 | PP3_Mod | VUS | 0.325 | 1592 | PMID:16732084; PMID:16917943 |
| chr19:g.38976319T>C | c.5024T>C | p.(Leu1675Pro) | 0.798 | PS4_Mod | VUS | 0.325 | 1675 | PMID:30236257 |
| chr19:g.38976328A>G | c.5033A>G | p.(Asn1678Ser) | 0.371 | BP4 | VUS | 0.100 | 1678 | PMID:30236257 ; PMID:22473935; PMID:27586648 |
| chr19:g.38976427A>G | c.5132A>G | p.(Tyr1711Cys) | 0.805 | PS4_Sup, PP1 | VUS | 0.325 | 1711 | PMID:25735680 |
| chr19:g.38976481T>G | c.5186T>G | p.(Met1729Arg) | 0.712 | PS4_Sup | VUS | 0.188 | 1729 | PMID:30236257 ; PMID:16917943 |
| chr19:g.38976636T>C | c.5341T>C | p.(Cys1781Arg) | 0.298 | PS4_Sup, BP4 | VUS | 0.100 | 1781 | PMID:30236257 |
| chr19:g.38976735A>G | c.5440A>G | p.(Met1814Val) | 0.486 | PS4_Sup, BP4 | VUS | 0.100 | 1814 | PMID:30236257 |
| chr19:g.38976736T>A | c.5441T>A | p.(Met1814Lys) | 0.81 | None | VUS | 0.100 | 1814 | PMID:16917943; PMID:30236257 |
| chr19:g.38980791C>T | c.5890C>T | p.(Arg1964Cys) | 0.232 | BP4 | VUS | 0.100 | 1964 | PMID:21965348; PMID:25989378 |
| chr19:g.38981282A>C | c.6037A>C | p.(Lys2013Gln) | 0.397 | PS4_Sup, BP4 | VUS | 0.100 | 2013 | PMID:20681998 |
| chr19:g.38985019T>A | c.6302T>A | p.(Met2101Lys) | 0.392 | PM1, BP4 | VUS | 0.188 | 2101 | PMID:30236257 ; PMID:12709367; PMID:20681998; PMID:25735680 |
| chr19:g.38985021G>C | c.6304G>C | p.(Val2102Leu) | 0.515 | PM1 | VUS | 0.325 | 2102 | PMID:23035052 |
| chr19:g.38985066G>C | c.6349G>C | p.(Val2117Leu) | 0.923 | PS4_Mod, PM1, PP3_Mod | Likely Pathogenic | 0.900 | 2117 | PMID:20681998; PMID:12709367; PMID:16917943 |
| chr19:g.38985094G>A | c.6377G>A | p.(Arg2126Gln) | 0.796 | PS4_Sup, PM1 | VUS | 0.500 | 2126 | PMID:24433488; PMID:21455645 |
| chr19:g.38985104C>G | c.6387C>G | p.(Asp2129Glu) | 0.683 | PS4_Mod, PM1, PP1_Mod | Likely Pathogenic | 0.900 | 2129 | PMID:12059893; PMID:24433488; PMID:11241852; PMID:12434264; PMID:16917943 |
| chr19:g.38985105G>A | c.6388G>A | p.(Gly2130Arg) | 0.894 | PM1, PP3_Mod | VUS | 0.675 | 2130 | PMID:18063506; PMID:18502356; PMID:26779337 |

| | | | | | | | | |
|---|---|---|---|---|---|---|---|---|
| chr19:g.38985205G>C | c.6488G>C | p.(Arg2163Pro) | 0.947 | PS4_Mod, PM1_Sup, PM5, PP1, PP3_Mod | Likely Pathogenic | 0.975 | 2163 | PMID:10757649; PMID:20681998; PMID:24433488; PMID:31559918; PMID:12709367; PMID:14500992; PMID:16917943; PMID:19191333; PMID:23447461; PMID:25895462; PMID:27646467 |
| chr19:g.38985205G>T | c.6488G>T | p.(Arg2163Leu) | 0.926 | PM1_Sup, PM5, PP3_Mod | VUS | 0.812 | 2163 | PMID:32919876 |
| chr19:g.38985261A>T | c.6544A>T | p.(Ile2182Phe) | 0.76 | PS4_Sup, PM1 | VUS | 0.500 | 2182 | PMID:12434264; PMID:14641996; PMID:24445638; PMID:15210166 |
| chr19:g.38985265G>A | c.6548G>A | p.(Gly2183Glu) | 0.786 | PS4_Sup, PM1 | VUS | 0.500 | 2183 | PMID:25735680 |
| chr19:g.38986905C>T | c.6599C>T | p.(Ala2200Val) | 0.564 | PS4_Sup, PM1, PP1_Mod | VUS | 0.812 | 2200 | PMID:15731587; PMID:21455645; PMID:30236257 ; PMID:16835904; PMID:16917943; PMID:19191333; PMID:23460944; PMID:24433488; PMID:26068069; PMID:30122538; PMID:31517061 |
| chr19:g.38986918C>G | c.6612C>G | p.(His2204Gln) | 0.739 | PS4_Mod, PM1, PP1_Mod | Likely Pathogenic | 0.900 | 2204 | PMID:30236257 |
| chr19:g.38986934G>T | c.6628G>T | p.(Val2210Phe) | 0.961 | PS4_Mod, PM1, PP3_Mod | Likely Pathogenic | 0.900 | 2210 | PMID:15731587; PMID:16917943; PMID:18362591 |
| chr19:g.38986941T>A | c.6635T>A | p.(Val2212Asp) | 0.973 | PM1, PP3_Mod | VUS | 0.675 | 2212 | PMID:16835904; PMID:19191333 |
| chr19:g.38986946G>A | c.6640G>A | p.(Val2214Ile) | 0.798 | PM1 | VUS | 0.325 | 2214 | PMID:11575529; PMID:11525881; PMID:14500992; PMID:15448513; PMID:16917943; PMID:25637381 |
| chr19:g.38987055C>T | c.6670C>T | p.(Arg2224Cys) | 0.728 | PM1, BS1 | Likely Benign | 0.025 | 2224 | PMID:30236257 |
| chr19:g.38987095G>A | c.6710G>A | p.(Cys2237Tyr) | 0.939 | PS4_Sup, PM1, PP3_Mod | VUS | 0.812 | 2237 | PMID:24433488; PMID:25960145; PMID:34462577 |
| chr19:g.38987127C>T | c.6742C>T | p.(Arg2248Cys) | 0.694 | PM1 | VUS | 0.325 | 2248 | PMID:20681998; PMID:30236257 ; PMID:25658027; PMID:31559918 |
| chr19:g.38987128G>A | c.6743G>A | p.(Arg2248His) | 0.654 | PM1 | VUS | 0.325 | 2248 | PMID:23558838 |
| chr19:g.38987142C>T | c.6757C>T | p.(His2253Tyr) | 0.973 | PS4_Sup, PM1, PP3_Mod | VUS | 0.812 | 2253 | PMID:21965348; PMID:29028638 |
| chr19:g.38987541G>A | c.6838G>A | p.(Val2280Ile) | 0.641 | PS4_Mod, PM1 | VUS | 0.675 | 2280 | PMID:30236257 ; PMID:30864471; PMID:12208234; PMID:16835904; PMID:16917943; PMID:27431030; PMID:28007021; PMID:30287922; PMID:30788618; PMID:31559918 |
| chr19:g.38987550A>C | c.6847A>C | p.(Asn2283His) | 0.943 | PS4_Sup, PM1, PP3_Mod | VUS | 0.812 | 2283 | PMID:20681998; PMID:16940308; PMID:17365175; PMID:17483490; PMID:23919265; PMID:30611313; PMID:19027160 |
| chr19:g.38989817A>G | c.6961A>G | p.(Ile2321Val) | 0.712 | PM1, BS1, BS2, BS3_Sup | Benign | <0.001 | 2321 | PMID:30236257 ; PMID:23558838; PMID:16917943; PMID:24055113; PMID:24195946; PMID:26972305; PMID:30788618 |
| chr19:g.38989874T>C | c.7018T>C | p.(Phe2340Leu) | 0.789 | PS4_Sup, PM1 | VUS | 0.500 | 2340 | PMID:25960145; PMID:28259615; PMID:32978841 |
| chr19:g.38989881A>G | c.7025A>G | p.(Asn2342Ser) | 0.639 | PS3_Mod, PM1, PP1, BS1, BS2 | Likely Benign | 0.012 | 2342 | PMID:30236257 ; PMID:25960145; PMID:23558838; PMID:15221887; PMID:16917943; PMID:19191333; PMID:20681998; PMID:22473935; PMID:23329375; PMID:23826317; PMID:24055113; PMID:24195946; PMID:24433488; PMID:25985138; PMID:30788618; PMID:28224104; PMID:16084090; PMID:19825159; PMID:28259615; PMID:31903994; PMID:31903994 |
| chr19:g.38990279G>C | c.7032G>C | p.(Glu2344Asp) | 0.657 | PM1 | VUS | 0.325 | 2344 | PMID:16163667; PMID:16917943 |
| chr19:g.38990282C>A | c.7035C>A | p.(Ser2345Arg) | 0.747 | PS3_Mod, PS4_Mod, PM1 | Likely Pathogenic | 0.900 | 2345 | PMID:25960145; PMID:Valaskova 2013; PMID:32535660; PMID:33490280 |
| chr19:g.38990283G>A | c.7036G>A | p.(Val2346Met) | 0.944 | PS4_Mod, PM1, PP3_Mod | Likely Pathogenic | 0.900 | 2346 | PMID:16163667; PMID:30236257 ; PMID:14985404; PMID:16917943; Valaskova 2013 |
| chr19:g.38990290A>G | c.7043A>G | p.(Glu2348Gly) | 0.964 | PS4_Mod, PM1, PP3_Mod | Likely Pathogenic | 0.900 | 2348 | PMID:30236257 ; PMID:14985404; PMID:16917943 |
| chr19:g.38990307G>A | c.7060G>A | p.(Val2354Met) | 0.867 | PS4_Mod, PM1, PP1_Mod, PP3_Mod | Likely Pathogenic | 0.975 | 2354 | PMID:23558838; PMID:24361844; PMID: 30864471; PMID:31559918 |
| chr19:g.38990322C>T | c.7075C>T | p.(Arg2359Trp) | 0.93 | PM1_Sup, PM5_Sup, PP3_Mod | VUS | 0.675 | 2359 | PMID:16521288 |
| chr19:g.38990323G>A | c.7076G>A | p.(Arg2359Gln) | 0.873 | PS4_Mod, PM1, PP3_Mod | Likely Pathogenic | 0.900 | 2359 | PMID:24433488; PMID:30236257 |
| chr19:g.38990331G>A | c.7084G>A | p.(Glu2362Lys) | 0.885 | PS4_Mod, PM1, PP3_Mod | Likely Pathogenic | 0.900 | 2362 | PMID:30236257 |
| chr19:g.38990332A>G | c.7085A>G | p.(Glu2362Gly) | 0.944 | PS4_Sup, PM1_Sup, PM5_Sup, PP3_Mod | VUS | 0.812 | 2362 | PMID:24433488; PMID:16835904; PMID:18813041; PMID:22473935 |
| chr19:g.38990336C>G | c.7089C>G | p.(Cys2363Trp) | 0.78 | PS4_Mod, PM1 | VUS | 0.675 | 2363 | PMID:30236257 |
| chr19:g.38990337T>G | c.7090T>G | p.(Phe2364Val) | 0.85 | PS4_Mod, PM1, PP3_Mod | Likely Pathogenic | 0.900 | 2364 | PMID:30236257 ; PMID:16917943; PMID:20566647 |
| chr19:g.38990344C>G | c.7097C>G | p.(Pro2366Arg) | 0.88 | PS4_Sup, PM1, PP3_Mod | VUS | 0.812 | 2366 | PMID:16732084; PMID:31559918; PMID:16917943 |

| | | | | | | | | |
|---|---|---|---|---|---|---|---|---|
| chr19:g.38990346G>A | c.7099G>A | p.(Ala2367Thr) | 0.899 | PS4_Sup, PM1, PP3_Mod | VUS | 0.812 | 2367 | PMID:11575529; PMID:15448513; PMID:15731587; PMID:16917943; PMID:25637381; PMID:11525881; PMID:12411786; PMID:23447461 |
| chr19:g.38990359A>G | c.7112A>G | p.(Glu2371Gly) | 0.891 | PS4_Sup, PM1, PP3_Mod | VUS | 0.812 | 2371 | PMID:24433488; PMID:19191333; PMID:25214167 |
| chr19:g.38990446A>G | c.7199A>G | p.(Asp2400Gly) | 0.548 | PM1 | VUS | 0.325 | 2400 | PMID:20681998 |
| chr19:g.38990457G>A | c.7210G>A | p.(Glu2404Lys) | 0.676 | PM1, PP1 | VUS | 0.500 | 2404 | PMID:19191329; PMID:25637381; PMID:19020143; PMID:23447461; PMID:30325262 |
| chr19:g.38990624G>A | c.7291G>A | p.(Asp2431Asn) | 0.888 | PS4_Mod, PM1, PP3_Mod | Likely Pathogenic | 0.900 | 2431 | PMID:11575529; PMID:30236257 ; PMID:15448513; PMID:16917943; PMID:25637381; PMID:11525881; PMID:12411786; PMID:14500992; PMID:23447461; PMID:31517061 |
| chr19:g.38990624G>T | c.7291G>T | p.(Asp2431Tyr) | 0.964 | PS3_Mod, PS4_Mod, PM1_Sup, PM5_Sup, PP1, PP3_Mod | Likely Pathogenic | 0.988 | 2431 | PMID:30236257 ; PMID:30115273; PMID:16917943; PMID:19648156; PMID:20566647 |
| chr19:g.38990625A>T | c.7292A>T | p.(Asp2431Val) | 0.952 | PS4_Sup, PM1_Sup, PM5_Sup, PP3_Mod | VUS | 0.812 | 2431 | PMID:30236257 |
| chr19:g.38990637G>T | c.7304G>T | p.(Arg2435Leu) | 0.948 | PS4_Mod, PM1_Sup, PM5, PP3_Mod | Likely Pathogenic | 0.949 | 2435 | PMID:10051009; PMID:23558838; PMID:30236257 ; PMID:10484775; PMID:12208234; PMID:12709367; PMID:14708096; PMID:15347586; PMID:16835904; PMID:16917943; PMID:17483490; PMID:19027160; PMID:20681998; PMID:23183335; PMID:25214167; PMID:28687594 |
| chr19:g.38990640G>A | c.7307G>A | p.(Cys2436Tyr) | 0.921 | PS4_Sup, PM1, PP3_Mod | VUS | 0.812 | 2436 | PMID:30236257 |
| chr19:g.38990643C>T | c.7310C>T | p.(Ala2437Val) | 0.917 | PS4_Mod, PM1, PP3_Mod | Likely Pathogenic | 0.900 | 2437 | PMID:15448513; PMID:16835904; PMID:16917943; PMID:23447461 |
| chr19:g.38990650G>C | c.7317G>C | p.(Glu2439Asp) | 0.734 | PM1 | VUS | 0.325 | 2439 | PMID:16835904 |
| chr19:g.38991277G>A | c.7355G>A | p.(Arg2452Gln) | 0.928 | PM1, PP3_Mod | VUS | 0.675 | 2452 | PMID:16732084; PMID:16917943 |
| chr19:g.38991277G>C | c.7355G>C | p.(Arg2452Pro) | 0.953 | PM1_Sup, PM5_Sup, PP3_Mod | VUS | 0.675 | 2452 | PMID:19191333 |
| chr19:g.38991280T>C | c.7358T>C | p.(Ile2453Thr) | 0.891 | PS2_PM6_Mod, PS4_Sup, PM1, PP3_Mod | Likely Pathogenic | 0.949 | 2453 | PMID:12434264; PMID:12810058; PMID:14708096; PMID:15108991; PMID:16917943; PMID:16958617; PMID:24433488; PMID:21514828 |
| chr19:g.38991295G>T | c.7373G>T | p.(Arg2458Leu) | 0.968 | PS4_Mod, PM1_Sup, PM5, PP3_Mod | Likely Pathogenic | 0.949 | 2458 | PMID:21965348; PMID:30236257 |
| chr19:g.38991503C>T | c.7487C>T | p.(Pro2496Leu) | 0.96 | PP3_Mod | VUS | 0.325 | 2496 | PMID:16732084; PMID:16917943; PMID:24195946; PMID:19685112 |
| chr19:g.38991544T>C | c.7528T>C | p.(Tyr2510His) | 0.922 | PP3_Mod | VUS | 0.325 | 2510 | PMID:16917943; PMID:30236257 ; PMID:19825159; PMID:19926015 |
| chr19:g.38993292A>G | c.7760A>G | p.(Tyr2587Cys) | 0.911 | PP3_Mod | VUS | 0.325 | 2587 | PMID:25960145 |
| chr19:g.38993303C>G | c.7771C>G | p.(Arg2591Gly) | 0.642 | None | VUS | 0.100 | 2591 | PMID:16835904 |
| chr19:g.38993303C>T | c.7771C>T | p.(Arg2591Trp) | 0.697 | None | VUS | 0.100 | 2591 | PMID:16917943; PMID:16835904 |
| chr19:g.38993309C>G | c.7777C>G | p.(Arg2593Gly) | 0.769 | None | VUS | 0.100 | 2593 | PMID:20681998 |
| chr19:g.38993310G>A | c.7778G>A | p.(Arg2593His) | 0.825 | PS4_Sup | VUS | 0.188 | 2593 | PMID:30236257 ; PMID:17968765 |
| chr19:g.38993319C>T | c.7787C>T | p.(Thr2596Ile) | 0.822 | None | VUS | 0.100 | 2596 | PMID:16917943 |
| chr19:g.38993348T>A | c.7816T>A | p.(Cys2606Ser) | 0.818 | BS2_Mod | Likely Benign | 0.025 | 2606 | PMID:30236257 ; PMID:21603587; PMID:23417605 |
| chr19:g.38993563G>A | c.7879G>A | p.(Val2627Met) | 0.808 | PS4_Mod | VUS | 0.325 | 2627 | PMID:30236257 ; PMID:24013571 |
| chr19:g.38993563G>C | c.7879G>C | p.(Val2627Leu) | 0.894 | PS4_Mod, PP1_St, PP3_Mod | Likely Pathogenic | 0.975 | 2627 | PMID:30236257 ; PMID:16835904; PMID:30916033 |
| chr19:g.38994938G>A | c.8005G>A | p.(Glu2669Lys) | 0.82 | None | VUS | 0.100 | 2669 | PMID:21157159; PMID:29635721 |
| chr19:g.38994959C>T | c.8026C>T | p.(Arg2676Trp) | 0.601 | PS4_Mod, PP1_St | Likely Pathogenic | 0.900 | 2676 | PMID:30236257 ; PMID:16163667; PMID:25960145; PMID:16917943; PMID:19191329; PMID:19919814; PMID:21157159; PMID:30788618; PMID:14732627 |
| chr19:g.38994987C>T | c.8054C>T | p.(Ser2685Phe) | 0.777 | PS4_Sup | VUS | 0.188 | 2685 | PMID:30236257 ; PMID:30788618; PMID:27431030 |
| chr19:g.38995508G>C | c.8188G>C | p.(Asp2730His) | 0.881 | PP3_Mod | VUS | 0.325 | 2730 | PMID:16732084; PMID:16917943; PMID:31559918; PMID:19020143; PMID:21795085; PMID:23447461; PMID:28259615 |
| chr19:g.38995509A>G | c.8189A>G | p.(Asp2730Gly) | 0.722 | PS4_Sup, PP1_St | VUS | 0.812 | 2730 | PMID:19191329; PMID:16917943; PMID:21088110; PMID:31559918; PMID:19191329; PMID:28259615 |
| chr19:g.38995518G>A | c.8198G>A | p.(Gly2733Asp) | 0.923 | PS4_Mod, PP3_Mod | VUS | 0.675 | 2733 | PMID:15731587; PMID:30236257 ; PMID:16917943 |
| chr19:g.38995701G>A | c.8290G>A | p.(Glu2764Lys) | 0.842 | None | VUS | 0.100 | 2764 | PMID:16835904; PMID:22705209; PMID:22913516; PMID:23413940; PMID:23204524; PMID:31068157 |

| | | | | | | | | |
|---|---|---|---|---|---|---|---|---|
| chr19:g.38995965C>T | c.8327C>T | p.(Ser2776Phe) | 0.693 | BS1 | Likely Benign | 0.006 | 2776 | PMID:21965348; PMID:30236257 ; PMID:21514828; PMID:22705209; PMID:24195946; PMID:25658027; PMID:25960145; PMID:30788618; PMID:32054689 ; PMID:23204524 |
| chr19:g.38995998C>G | c.8360C>G | p.(Thr2787Ser) | 0.466 | BA1 | Benign | <0.001 | 2787 | PMID:16163667; PMID:19191329; PMID:25735680; PMID:30236257 ; PMID:14732627; PMID:16917943; PMID:20839240; PMID:20981092; PMID:22473935; PMID:22705209; PMID:22913516; PMID:15315621; PMID:23204524; PMID:24195946 |
| chr19:g.38996563C>T | c.8518C>T | p.(Arg2840Trp) | 0.659 | None | VUS | 0.100 | 2840 | PMID:16732084; PMID:16917943; PMID:22705209; PMID:22913516; PMID:23204524 |
| chr19:g.38996572T>C | c.8527T>C | p.(Ser2843Pro) | 0.761 | PS4_Sup | VUS | 0.188 | 2843 | PMID:21455645; PMID:22705209; PMID:23460944; PMID:23204524 |
| chr19:g.38997001T>A | c.8600T>A | p.(Leu2867Gln) | 0.923 | PP3_Mod | VUS | 0.325 | 2867 | PMID:16835904 |
| chr19:g.38997132G>A | c.8638G>A | p.(Glu2880Lys) | 0.901 | PS4_Sup, PP3_Mod | VUS | 0.500 | 2880 | PMID:19191329; PMID:16917943; PMID:22705209; PMID:22913516; PMID:23447461 |
| chr19:g.38997148C>G | c.8654C>G | p.(Thr2885Arg) | 0.656 | None | VUS | 0.100 | 2885 | PMID:25735680 |
| chr19:g.38997505C>T | c.8729C>T | p.(Thr2910Met) | 0.779 | PS4_Sup | VUS | 0.188 | 2910 | PMID:30236257 ; PMID:30916033 |
| chr19:g.38998461C>T | c.8926C>T | p.(His2976Tyr) | 0.201 | PS4_Sup, BP4 | VUS | 0.100 | 2976 | PMID:20681998 |
| chr19:g.39002230G>A | c.9152G>A | p.(Arg3051His) | 0.581 | None | VUS | 0.100 | 3051 | PMID:30236257 |
| chr19:g.39002919G>A | c.9268G>A | p.(Ala3090Thr) | 0.454 | PS4_Sup, BP4 | VUS | 0.100 | 3090 | PMID:30236257 |
| chr19:g.39003007G>A | c.9356G>A | p.(Arg3119His) | 0.711 | None | VUS | 0.100 | 3119 | PMID:16732084; PMID:16917943; PMID:23919265; PMID:30611313 |
| chr19:g.39005692C>T | c.9499C>T | p.(Arg3167Term) | | None | VUS | 0.100 | 3167 | PMID:26951757; PMID:23558838; PMID:27562486 |
| chr19:g.39006807A>G | c.9635A>G | p.(Glu3212Gly) | 0.624 | None | VUS | 0.100 | 3212 | PMID:30236257 ; PMID:29590070 |
| chr19:g.39006821T>C | c.9649T>C | p.(Ser3217Pro) | 0.918 | PP3_Mod | VUS | 0.325 | 3217 | PMID:19191329; PMID:24433488; PMID:23447461; PMID:28259615 |
| chr19:g.39006824G>A | c.9652G>A | p.(Val3218Met) | 0.608 | PS4_Sup | VUS | 0.188 | 3218 | PMID:30236257 |
| chr19:g.39006848G>C | c.9676G>C | p.(Glu3226Gln) | 0.62 | PS4_Sup | VUS | 0.188 | 3226 | PMID:30236257 ; PMID:25658027 |
| chr19:g.39008110T>C | c.9797T>C | p.(Met3266Thr) | 0.742 | PS4_Sup | VUS | 0.188 | 3266 | PMID:30236257 |
| chr19:g.39008161G>A | c.9848G>A | p.(Arg3283Gln) | 0.527 | None | VUS | 0.100 | 3283 | PMID:23558838; PMID:25268394 |
| chr19:g.39008163T>A | c.9850T>A | p.(Trp3284Arg) | 0.873 | PP3_Mod | VUS | 0.325 | 3284 | PMID:21455645; PMID:26068069 |
| chr19:g.39008181G>A | c.9868G>A | p.(Glu3290Lys) | 0.848 | None | VUS | 0.100 | 3290 | PMID:19191329; PMID:23447461 |
| chr19:g.39009878G>A | c.10043G>A | p.(Arg3348His) | 0.705 | PS4_Sup | VUS | 0.188 | 3348 | PMID:15731587; PMID:25735680; PMID:16835904; PMID:16917943 |
| chr19:g.39009935A>G | c.10100A>G | p.(Lys3367Arg) | 0.632 | BS3_Sup | Likely Benign | 0.051 | 3367 | PMID:16732084; PMID:16621918; PMID:16917943; PMID:21674524; PMID:32535660 |
| chr19:g.39010064C>A | c.10229C>A | p.(Pro3410Gln) | 0.934 | PP3_Mod | VUS | 0.325 | 3410 | PMID:20681998 |
| chr19:g.39010072A>T | c.10237A>T | p.(Ile3413Phe) | 0.914 | PS4_Sup, PP3_Mod | VUS | 0.500 | 3413 | PMID:25735680 |
| chr19:g.39010087A>G | c.10252A>G | p.(Asn3418Asp) | 0.535 | PS4_Mod | VUS | 0.325 | 3418 | PMID:30236257 |
| chr19:g.39016132G>A | c.10616G>A | p.(Arg3539His) | 0.877 | PP3_Mod, BS1, BS2 | Likely Benign | 0.001 | 3539 | PMID:30236257 ; PMID:23460944; PMID:23558838; PMID:24433488; PMID:25960145; PMID:16521288; PMID:18253926; PMID:22473935; PMID:24055113; PMID:24195946; PMID:30788618; PMID:31559918; PMID:32054689 |
| chr19:g.39018350G>C | c.10750G>C | p.(Glu3584Gln) | 0.798 | None | VUS | 0.100 | 3584 | PMID:16917943 |
| chr19:g.39019012G>T | c.10891G>T | p.(Ala3631Ser) | 0.85 | PS4_Sup, PP3_Mod | VUS | 0.500 | 3631 | PMID:30236257 |
| chr19:g.39019642G>C | c.11086G>C | p.(Asp3696His) | 0.809 | PS4_Sup | VUS | 0.188 | 3696 | PMID:30236257 |
| chr19:g.39019676G>T | c.11120G>T | p.(Arg3707Leu) | 0.906 | PP3_Mod | VUS | 0.325 | 3707 | PMID:16917943 |
| chr19:g.39019682C>T | c.11126C>T | p.(Ala3709Val) | 0.793 | PS4_Sup | VUS | 0.188 | 3709 | PMID:21965348 |
| chr19:g.39019688C>T | c.11132C>T | p.(Thr3711Met) | 0.637 | PS4_Sup, PP1 | VUS | 0.325 | 3711 | PMID:30236257 ; PMID:32054689 ; PMID: 30916033 |
| chr19:g.39025837G>A | c.11416G>A | p.(Gly3806Arg) | 0.933 | PS4_Sup, PP1, PP3_Mod | VUS | 0.675 | 3806 | PMID:19191329 |
| chr19:g.39026638G>A | c.11518G>A | p.(Val3840Ile) | 0.648 | None | VUS | 0.100 | 3840 | PMID:16732084; PMID:16917943; PMID:25637381 |
| chr19:g.39034005G>A | c.11708G>A | p.(Arg3903Gln) | 0.97 | PS4, PP3_Mod | Likely Pathogenic | 0.900 | 3903 | PMID:20681998; PMID:24433488; PMID:30236257 ; PMID:16835904; PMID:16917943; PMID:19191333; PMID:19252784; PMID:25958340 |
| chr19:g.39034020A>T | c.11723A>T | p.(Asn3908Ile) | 0.949 | PS4_Sup, PP3_Mod | VUS | 0.500 | 3908 | PMID:24433488 |
| chr19:g.39034045T>G | c.11748T>G | p.(Ile3916Met) | 0.765 | None | VUS | 0.100 | 3916 | PMID:12411788; PMID:16163667; PMID:16917943 |
| chr19:g.39034206G>A | c.11813G>A | p.(Gly3938Asp) | 0.799 | None | VUS | 0.100 | 3938 | PMID:21455645; PMID:21911697 |
| chr19:g.39034456T>C | c.11953T>C | p.(Trp3985Arg) | 0.941 | PS4_Mod, PP3_Mod | VUS | 0.675 | 3985 | PMID:18719443; PMID:23558838 |
| chr19:g.39037100G>A | c.12028G>A | p.(Glu4010Lys) | 0.178 | PS4_Sup, BP4 | VUS | 0.100 | 4010 | PMID:25658027; PMID:30236257 |
| chr19:g.39037136A>G | c.12064A>G | p.(Met4022Val) | 0.491 | BP4 | VUS | 0.100 | 4022 | PMID:21157159 |
| chr19:g.39038893A>T | c.12115A>T | p.(Ile4039Phe) | 0.621 | PS4_Sup | VUS | 0.188 | 4039 | PMID:30236257 |
| chr19:g.39038899C>T | c.12121C>T | p.(Arg4041Trp) | 0.435 | BP4 | VUS | 0.100 | 4041 | PMID:16835904; PMID:24055113; PMID:25637381 |
| chr19:g.39038927C>A | c.12149C>A | p.(Ser4050Tyr) | 0.942 | PS4_Mod, PP1_St, PP3_Mod | Likely Pathogenic | 0.975 | 4050 | PMID:30236257 ; PMID:16917943 |

| | | | | | | | | |
|---|---|---|---|---|---|---|---|---|
| chr19:g.39039020C>T | c.12242C>T | p.(Thr4081Met) | 0.649 | None | VUS | 0.100 | 4081 | PMID:16732084; PMID:16917943; PMID:27558158 |
| chr19:g.39051780G>C | c.12310G>C | p.(Gly4104Arg) | 0.307 | BS2_Mod, BP4 | Likely Benign | 0.012 | 4104 | PMID:21455645 |
| chr19:g.39051825A>T | c.12355A>T | p.(Asn4119Tyr) | 0.813 | PS4_Sup | VUS | 0.188 | 4119 | PMID:15731587; PMID:16917943; PMID:27558158 |
| chr19:g.39051853C>T | c.12383C>T | p.(Ala4128Val) | 0.248 | PS4_Sup, BP4 | VUS | 0.100 | 4128 | PMID:30236257 |
| chr19:g.39051868A>G | c.12398A>G | p.(Glu4133Gly) | 0.881 | PS4_Sup, PP3_Mod | VUS | 0.500 | 4133 | PMID:24433488 |
| chr19:g.39051876C>A | c.12406C>A | p.(Arg4136Ser) | 0.783 | BS2_Mod | Likely Benign | 0.025 | 4136 | PMID:12208234; PMID:16835904; PMID:16917943 |
| chr19:g.39051883T>C | c.12413T>C | p.(Ile4138Thr) | 0.96 | PS4_Sup, PP3_Mod | VUS | 0.500 | 4138 | PMID:24433488; PMID:16306761; PMID:16917943 |
| chr19:g.39052003G>T | c.12533G>T | p.(Gly4178Val) | 0.918 | PS4_Mod, PP3_Mod | VUS | 0.675 | 4178 | PMID:23558838; PMID:25735680; PMID:30236257 |
| chr19:g.39052023G>A | c.12553G>A | p.(Ala4185Thr) | 0.402 | BP4 | VUS | 0.100 | 4185 | PMID:21455645; PMID:30236257 ; PMID:31559918; PMID:27555149; PMID:23460944; PMID:24195946 |
| chr19:g.39055663T>G | c.12689T>G | p.(Met4230Arg) | 0.928 | PS4_Mod, PP3_Mod | VUS | 0.675 | 4230 | PMID:23558838; PMID:30236257 ; PMID:25658027 |
| chr19:g.39055674G>C | c.12700G>C | p.(Val4234Leu) | 0.89 | PS4_Mod, PP1_St, PP3_Mod | Likely Pathogenic | 0.975 | 4234 | PMID:30236257 ; PMID:12208234; PMID:21965348; PMID:28290972; PMID:16835904; PMID:16917943; PMID:19191333; PMID:20681998; PMID:22696611; PMID:24053352 |
| chr19:g.39055674G>T | c.12700G>T | p.(Val4234Leu) | 0.89 | PS1_Mod, PS4_Mod, PP1, PP3_Mod | Likely Pathogenic | 0.949 | 4234 | PMID:24013571 |
| chr19:g.39055822A>T | c.12848A>T | p.(Glu4283Val) | 0.561 | None | VUS | 0.100 | 4283 | PMID:16732084; PMID:16917943 |
| chr19:g.39061259C>T | c.13672C>T | p.(Arg4558Trp) | 0.79 | PS4_Sup | VUS | 0.188 | 4558 | PMID:30236257 |
| chr19:g.39062672C>T | c.13760C>T | p.(Pro4587Leu) | 0.785 | None | VUS | 0.100 | 4587 | PMID:16521288; PMID:25637381 |
| chr19:g.39062825G>A | c.13913G>A | p.(Gly4638Asp) | 0.917 | PM1_Sup, PP3_Mod | VUS | 0.500 | 4638 | PMID:17483490; PMID:12565913; PMID:14985404; PMID:16917943; PMID:23553787 |
| chr19:g.39062846G>A | c.13934G>A | p.(Arg4645Gln) | 0.567 | PM1_Sup | VUS | 0.188 | 4645 | PMID:16732084; PMID:16917943; PMID:19223216; PMID:19931341 |
| chr19:g.39062902T>C | c.13990T>C | p.(Cys4664Arg) | 0.936 | PS4_Sup, PM1_Sup, PP3_Mod | VUS | 0.675 | 4664 | PMID:19191333; PMID:24433488; PMID:27646467 |
| chr19:g.39062906_39062907delinsCT | c.13994_13995delTCinsCT | p.(Leu4665Pro) | | PS2_PM6_Sup, PM1_Sup | VUS | 0.325 | 4665 | PMID:17483490 |
| chr19:g.39063820C>T | c.14002C>T | p.(Pro4668Ser) | 0.888 | PM1_Sup, PP3_Mod | VUS | 0.500 | 4668 | PMID:11928716; PMID:16917943 |
| chr19:g.39063869T>C | c.14051T>C | p.(Phe4684Ser) | 0.982 | PS4_Sup, PM1_Sup, PP3_Mod | VUS | 0.675 | 4684 | PMID:16163667; PMID:16835904; PMID:16917943; PMID:23447461 |
| chr19:g.39066597G>A | c.14168G>A | p.(Arg4723His) | 0.79 | PM1_Sup, BS2_Mod | Likely Benign | 0.051 | 4723 | PMID:30236257 ; PMID:25658027 |
| chr19:g.39068571A>C | c.14186A>C | p.(His4729Pro) | 0.758 | PM1_Sup | VUS | 0.188 | 4729 | PMID:26951757; PMID:27562486 |
| chr19:g.39068582T>G | c.14197T>G | p.(Tyr4733Asp) | 0.842 | PS4_Sup, PM1_Sup | VUS | 0.325 | 4733 | PMID:15731587; PMID:16917943; PMID:27558158 |
| chr19:g.39068586G>A | c.14201G>A | p.(Gly4734Glu) | 0.871 | PS4_Sup, PM1_Sup, PP3_Mod | VUS | 0.675 | 4734 | PMID:30236257 ; PMID:16917943; PMID:27558158 |
| chr19:g.39068594C>T | c.14209C>T | p.(Arg4737Trp) | 0.882 | PS4_Mod, PM1_Sup, PM5_Sup, PP1, PP3_Mod | Likely Pathogenic | 0.949 | 4737 | PMID:16163667; PMID:26631338; PMID:30236257 ; PMID:29137581; PMID:12208234; PMID:16835904; PMID:16917943; PMID:23460944; PMID:27558158; PMID:20566647; PMID:23447461 |
| chr19:g.39068595G>A | c.14210G>A | p.(Arg4737Gln) | 0.89 | PS4, PM1_Sup, PP1_St, PP3_Mod, BS2_Mod | Likely Pathogenic | 0.988 | 4737 | PMID:30236257 ; PMID:16163667; PMID:18564801; PMID:24433488; PMID:29137581; PMID:16917943; PMID:19454545; PMID:19648156; PMID:23460944; PMID:23628358; PMID:25960145; PMID:26972305; PMID:27558158; PMID:27663056; PMID:28326467; PMID:30155738 ; PMID:30788618; PMID:30916033; PMID:19825159; PMID:20566647; PMID:23447461 |
| chr19:g.39068655G>A | c.14270G>A | p.(Arg4757His) | 0.452 | PM1_Sup, BP4 | VUS | 0.100 | 4757 | PMID:30236257 |
| chr19:g.39068845G>T | c.14364+1G>T | | | PS4_Sup, PM1_Sup | VUS | 0.325 | | PMID:25960145; PMID:30788618 |
| chr19:g.39070679_39070680delinsAA | c.14422_14423delinsAA | p.(Phe4808Asn) | | PM1_Sup | VUS | 0.188 | 4808 | PMID:21455645; PMID:12565913; PMID:27447704; PMID:28357410; PMID:30155738; PMID:32272370; PMID:23460944; PMID:23919265 |
| chr19:g.39070681C>A | c.14424C>A | p.(Phe4808Leu) | 0.923 | PS4_Sup, PM1_Sup, PP3_Mod | VUS | 0.675 | 4808 | PMID:23460944; PMID:23183335 |
| chr19:g.39070706A>T | c.14449A>T | p.(Ile4817Phe) | 0.911 | PS4_Sup, PM1_Sup, PP3_Mod | VUS | 0.675 | 4817 | PMID:30236257 ; PMID:16917943 |
| chr19:g.39070715G>A | c.14458G>A | p.(Gly4820Arg) | 0.835 | PS4_Sup, PM1_Sup | VUS | 0.325 | 4820 | PMID:23267001; PMID:28326467; PMID:30155738 |
| chr19:g.39070715G>T | c.14458G>T | p.(Gly4820Trp) | 0.918 | PS4_Sup, PM1_Sup, PP3_Mod | VUS | 0.675 | 4820 | PMID:30236257 ; PMID:16917943; PMID:28326467 |

| | | | | | | | | |
|---|---|---|---|---|---|---|---|---|
| chr19:g.39070728T>C | c.14471T>C | p.(Leu4824Pro) | 0.984 | PS4_Mod, PM1_Sup, PP3_Mod | VUS | 0.812 | 4824 | PMID:30236257 ; PMID:15448513; PMID:14985404; PMID:16917943; PMID:31559918; PMID:20566647; PMID:23447461; PMID:23919265; PMID:29674523 |
| chr19:g.39070766C>G | c.14509C>G | p.(Gln4837Glu) | 0.914 | PS4_Sup, PM1_Sup, PP3_Mod | VUS | 0.675 | 4837 | PMID:23558838; PMID:31559918 |
| chr19:g.39070767del | c.14510delA | p.(Gln4837Argfs*3) | | PS4_Sup, BS2_Mod, BS3_Sup | Likely Benign | 0.025 | 4837 | PMID:17293538; PMID:16917943 |
| chr19:g.39071022G>A | c.14524G>A | p.(Val4842Met) | 0.933 | PM1_Sup, PP3_Mod | VUS | 0.500 | 4842 | PMID:21455645; PMID:18253926; PMID:20839240; PMID:22473935 |
| chr19:g.39071037G>C | c.14539G>C | p.(Val4847Leu) | 0.762 | PS4_Sup, PM1_Sup, PP1_St | Likely Pathogenic | 0.900 | 4847 | PMID:21455645; PMID:23558838; PMID:29635721 |
| chr19:g.39071056C>T | c.14558C>T | p.(Thr4853Ile) | 0.982 | PM1_Sup, PP3_Mod | VUS | 0.500 | 4853 | PMID:16244682; PMID:22473935; PMID:23394784 |
| chr19:g.39071065C>G | c.14567C>G | p.(Ala4856Gly) | 0.937 | PM1_Sup, PP3_Mod | VUS | 0.500 | 4856 | PMID:16917943; PMID:28357410 |
| chr19:g.39071079C>T | c.14581C>T | p.(Arg4861Cys) | 0.916 | PS4_Sup, PM1_Sup, PM5_Sup, PP3_Mod | VUS | 0.812 | 4861 | PMID:30236257 ; PMID:12565913; PMID:16621918; PMID:16917943; PMID:17226826; PMID:17483490; PMID:21088110; PMID:21911697; PMID:23553484; PMID:23919265; PMID:25960145; PMID:26684984; PMID:26799446; PMID:27066551; PMID:27447704; PMID:15564033; PMID:16084090; PMID:29449963; PMID:29669168; PMID:33333461; PMID:33458582 |
| chr19:g.39071125A>G | c.14627A>G | p.(Lys4876Arg) | 0.912 | PS4_Mod, PM1_Sup, PP1_Mod, PP3_Mod | Likely Pathogenic | 0.949 | 4876 | PMID:15731587; PMID:16163667; PMID:19191329; PMID:24433488; PMID:16917943; PMID:27854207 |
| chr19:g.39071137T>C | c.14639T>C | p.(Met4880Thr) | 0.956 | PS4_Sup, PM1_Sup, PP3_Mod | VUS | 0.675 | 4880 | PMID:15731587; PMID:16917943 |
| chr19:g.39075614G>A | c.14678G>A | p.(Arg4893Gln) | 0.963 | PM1_Sup, PP3_Mod | VUS | 0.500 | 4893 | PMID:12565913; PMID:16621918; PMID:16917943; PMID:17483490; PMID:22473935; PMID:22550088; PMID:23183335; PMID:25214167; PMID:25331388; PMID:25628744; PMID:30236257 ; PMID:15564033 |
| chr19:g.39075616G>A | c.14680G>A | p.(Ala4894Thr) | 0.922 | PS3_Mod, PM1_Sup, PP3_Mod | VUS | 0.812 | 4894 | PMID:16732084; PMID:16917943; PMID:21926372; PMID:33490280 |
| chr19:g.39075718A>G | c.14782A>G | p.(Ile4928Val) | 0.837 | PM1_Sup | VUS | 0.188 | 4928 | PMID:21455645 |
| chr19:g.39075739G>A | c.14803G>A | p.(Gly4935Ser) | 0.962 | PS4_Mod, PM1_Sup, PP1_St, PP3_Mod | Likely Pathogenic | 0.988 | 4935 | PMID:19918671; PMID:21455645 |
| chr19:g.39076587T>C | c.14813T>C | p.(Ile4938Thr) | 0.956 | PS4_Sup, PM1_Sup, PP1_St, PP3_Mod, BS2_Mod | Likely Pathogenic | 0.900 | 4938 | PMID:19191329; PMID:21795085 |
| chr19:g.39076588C>G | c.14814C>G | p.(Ile4938Met) | 0.855 | PM1_Sup, PP3_Mod | VUS | 0.500 | 4938 | PMID:14985404; PMID:16917943; PMID:30236257 |
| chr19:g.39076591C>A | c.14817C>A | p.(Asp4939Glu) | 0.808 | PS4_Sup, PM1_Sup | VUS | 0.325 | 4939 | PMID:23558838; PMID:14985404; PMID:16163667; PMID:16917943; PMID:30236257 |
| chr19:g.39076592G>A | c.14818G>A | p.(Ala4940Thr) | 0.882 | PS4_Mod, PM1_Sup, PP3_Mod | VUS | 0.812 | 4940 | PMID:15731587; PMID:12467748; PMID:12565913; PMID:14670767; PMID:16917943; PMID:23183335; PMID:23558838; PMID:28687594; PMID:30155738 ; PMID:31559918 |
| chr19:g.39076599G>T | c.14825G>T | p.(Gly4942Val) | 0.922 | PM1_Sup, PP3_Mod | VUS | 0.500 | 4942 | PMID:12208234; PMID:16917943 |
| chr19:g.39076741T>A | c.14879T>A | p.(Phe4960Tyr) | 0.965 | PM1_Sup, PP3_Mod | VUS | 0.500 | 4960 | PMID:16732084; PMID:16917943 |
| chr19:g.39076830A>G | c.14968A>G | p.(Met4990Val) | 0.646 | PS4_Sup, PM1_Sup | VUS | 0.325 | 4990 | PMID:21455645; PMID:26068069 |
| chr19:g.39078002G>C | c.15059G>C | p.(Trp5020Ser) | 0.918 | PP3_Mod | VUS | 0.325 | 5020 | PMID:24433488 |
| chr19:g.39078003G>C | c.15060G>C | p.(Trp5020Cys) | 0.892 | PS4_Sup, PP3_Mod | VUS | 0.500 | 5020 | PMID:25960145 |

Supplemental Table 2

| Genomic Coordinate (GRCh37) | cDNA NM_000540.2 | Protein | REVEL | ACMG / AMP Criteria | Pathogenicity | Posterior Probability | Amino Acid Position | Publication |
|---|---|---|---|---|---|---|---|---|
| chr19:g.38924507T>G | c.38T>G | p.(Leu13Arg) | 0.828 | PS3_Mod, PS4_Mod, PM1, PP1 | Likely Pathogenic | 0.949 | 13 | This work |
| chr19:g.38931390_38931392del | c.51_53del | p.(Asp17del) | | PM1 | VUS | 0.325 | 17 | This work |
| chr19:g.38931436A>G | c.97A>G | p.(Lys33Glu) | 0.92 | PS2_PM6_Sup, PS4_Sup, PM1, PP3_Mod | Likely Pathogenic | 0.900 | 33 | This work |
| chr19:g.38931442T>C | c.103T>C | p.(Cys35Arg) | 0.947 | PS4_Mod, PM1, PP1_St, PP3_Mod | Pathogenic | 0.994 | 35 | PMID: 33767344 |
| chr19:g.38931458G>C | c.119G>C | p.(Gly40Ala) | 0.897 | PS4_Sup, PM1, PP3_Mod | VUS | 0.812 | 40 | This work |
| chr19:g.38931469C>T | c.130C>T | p.(Arg44Cys) | 0.951 | PS3_Mod, PS4_Mod, PM1, PP3_Mod | Likely Pathogenic | 0.975 | 44 | PMID: 33767344 |
| chr19:g.38931470G>A | c.131G>A | p.(Arg44His) | 0.934 | PS4_Sup, PM1, PP3_Mod | VUS | 0.812 | 44 | This work |
| chr19:g.38931491C>A | c.152C>A | p.(Thr51Asn) | 0.769 | PM1 | VUS | 0.325 | 51 | PMID: 33767344 |
| chr19:g.38933001G>A | c.178G>A | p.(Asp60Asn) | 0.736 | PS4_Sup, PM1 | VUS | 0.500 | 60 | This work |
| chr19:g.38933001G>T | c.178G>T | p.(Asp60Tyr) | 0.961 | PS4_Sup, PM1, PP3_Mod | VUS | 0.812 | 60 | This work |
| chr19:g.38933013T>C | c.190T>C | p.(Cys64Arg) | 0.809 | PM1 | VUS | 0.325 | 64 | This work |
| chr19:g.38933035C>A | c.212C>A | p.(Ser71Tyr) | 0.937 | PM1, PP3_Mod | VUS | 0.675 | 71 | This work |
| chr19:g.38933074C>T | c.251C>T | p.(Thr84Met) | 0.735 | PS3_Mod, PM1 | VUS | 0.675 | 84 | This work |
| chr19:g.38934430G>A | c.418G>A | p.(Ala140Thr) | 0.261 | PM1, BS2_Mod, BP4 | Likely Benign | 0.051 | 140 | This work |
| chr19:g.38934819C>A | c.455C>A | p.(Ala152Asp) | 0.9 | PS4_Sup, PM1, PP3_Mod | VUS | 0.812 | 152 | This work |
| chr19:g.38934827C>A | c.463C>A | p.(Gln155Lys) | 0.94 | PS3_Mod, PM1, PP3_Mod | Likely Pathogenic | 0.900 | 155 | This work |
| chr19:g.38934831G>A | c.467G>A | p.(Arg156Lys) | 0.886 | PM1, PP3_Mod | VUS | 0.675 | 156 | This work |
| chr19:g.38934843A>G | c.479A>G | p.(Glu160Gly) | 0.957 | PS4_Sup, PM1, PP3_Mod | VUS | 0.812 | 160 | This work |
| chr19:g.38934851C>T | c.487C>T | p.(Arg163Cys) | 0.959 | PS3, PS4, PM1_Sup, PM5_Sup, PP1_St, PP3_Mod, BS2_Mod | Pathogenic | >0.999 | 163 | PMID: 33767344 |
| chr19:g.38934852G>A | c.488G>A | p.(Arg163His) | 0.864 | PS4_Sup, PM1, PP3_Mod | VUS | 0.812 | 163 | This work |
| chr19:g.38934852G>T | c.488G>T | p.(Arg163Leu) | 0.882 | PS3_Mod, PS4_Mod, PM1, PP3_Mod | Likely Pathogenic | 0.975 | 163 | PMID: 33767344 |
| chr19:g.38934857G>A | c.493G>A | p.(Gly165Arg) | 0.957 | PS4_Sup, PM1, PP3_Mod | VUS | 0.812 | 165 | This work |
| chr19:g.38934860G>A | c.496G>A | p.(Asp166Asn) | 0.829 | PS4_Mod, PM1 | VUS | 0.675 | 166 | This work |
| chr19:g.38934861A>G | c.497A>G | p.(Asp166Gly) | 0.967 | PM1, PP3_Mod | VUS | 0.675 | 166 | This work |
| chr19:g.38934890G>A | c.526G>A | p.(Glu176Lys) | 0.803 | PS4_Mod, PM1 | VUS | 0.675 | 176 | This work |
| chr19:g.38934892G>T | c.528G>T | p.(Glu176Asp) | 0.504 | PS4_Mod, PM1 | VUS | 0.675 | 176 | This work |
| chr19:g.38934893C>T | c.529C>T | p.(Arg177Cys) | 0.931 | PS2_PM6_Mod, PS4, PM1, PP1_St, PP3_Mod, BS2_Mod | Pathogenic | 0.999 | 177 | This work |
| chr19:g.38934897A>C | c.533A>C | p.(Tyr178Ser) | 0.901 | PS4_Sup, PM1, PP3_Mod | VUS | 0.812 | 178 | This work |
| chr19:g.38934897A>G | c.533A>G | p.(Tyr178Cys) | 0.951 | PS2_PM6_Mod, PS4_Sup, PM1, PP1, PP3_Mod | Likely Pathogenic | 0.975 | 178 | This work |
| chr19:g.38935311G>A | c.625G>A | p.(Glu209Lys) | 0.711 | PM1, BP2 | VUS | 0.188 | 209 | This work |
| chr19:g.38937121C>T | c.641C>T | p.(Thr214Met) | 0.533 | PS4_Mod, PM1, PP1, BS2_Mod, BS3_Sup | VUS | 0.325 | 214 | This work |
| chr19:g.38937132G>A | c.652G>A | p.(Val218Ile) | 0.801 | PS4_Sup, PM1 | VUS | 0.500 | 218 | This work |
| chr19:g.38937157T>A | c.677T>A | p.(Met226Lys) | 0.841 | PS4_Mod, PM1 | VUS | 0.675 | 226 | This work |
| chr19:g.38937160A>T | c.680A>T | p.(Asp227Val) | 0.933 | PS4_Sup, PM1, PP3_Mod | VUS | 0.812 | 227 | This work |
| chr19:g.38937350G>A | c.742G>A | p.(Gly248Arg) | 0.889 | PS3_Mod, PS4, PM1, PP1, PP3_Mod, BS2_Mod | Likely Pathogenic | 0.988 | 248 | PMID: 33767344 |
| chr19:g.38937350G>C | c.742G>C | p.(Gly248Arg) | 0.883 | PS1, PS3_Mod, PS4_Mod, PM1_Sup, PP1_Mod, PP3_Mod | Pathogenic | 0.999 | 248 | PMID: 33767344 |
| chr19:g.38939140C>T | c.946C>T | p.(Arg316Cys) | 0.793 | PM1 | VUS | 0.325 | 316 | This work |
| chr19:g.38939141G>T | c.947G>T | p.(Arg316Leu) | 0.928 | PM1, PP3_Mod | VUS | 0.675 | 316 | This work |
| chr19:g.38939313C>T | c.982C>T | p.(Arg328Trp) | 0.76 | PS3_Mod, PS4_Sup, PM1, PP1 | Likely Pathogenic | 0.900 | 328 | PMID: 33767344 |
| chr19:g.38939320_38939321insGGA | c.992_994dup | p.(Glu331dup) | | PS4_Sup, PM1 | VUS | 0.500 | 331 | This work |
| chr19:g.38939352G>A | c.1021G>A | p.(Gly341Arg) | 0.864 | PS3_Mod, PS4, PM1, PP1_St, PP3_Mod, BS2_Mod | Pathogenic | 0.999 | 341 | PMID: 33767344 |
| chr19:g.38939352G>C | c.1021G>C | p.(Gly341Arg) | 0.876 | PS1, PS3_Mod, PS4_Mod, PM1_Sup, PP1, PP3_Mod | Pathogenic | 0.999 | 341 | PMID: 33767344 |
| chr19:g.38939355G>A | c.1024G>A | p.(Glu342Lys) | 0.927 | PS4_Sup, PM1, PP3_Mod | VUS | 0.812 | 342 | This work |
| chr19:g.38939431G>A | c.1100G>A | p.(Arg367Gln) | 0.651 | PS4_Mod, PM1 | VUS | 0.675 | 367 | This work |
| chr19:g.38939431G>T | c.1100G>T | p.(Arg367Leu) | 0.866 | PS4_Sup, PM1, PP3_Mod | VUS | 0.812 | 367 | This work |
| chr19:g.38942425C>A | c.1144C>A | p.(His382Asn) | 0.739 | PS4_Sup, PM1 | VUS | 0.500 | 382 | This work |
| chr19:g.38942482C>A | c.1201C>A | p.(Arg401Ser) | 0.839 | PS4_Sup, PM1_Sup, PM5 | VUS | 0.675 | 401 | This work |
| chr19:g.38942482C>G | c.1201C>G | p.(Arg401Gly) | 0.865 | PM1_Sup, PM5, PP3_Mod | VUS | 0.812 | 401 | This work |
| chr19:g.38942482C>T | c.1201C>T | p.(Arg401Cys) | 0.886 | PS3_Mod, PS4_Mod, PM1_Sup, PM5, PP1, PP3_Mod | Pathogenic | 0.994 | 401 | PMID: 33767344 |
| chr19:g.38942483G>A | c.1202G>A | p.(Arg401His) | 0.903 | PS3_Mod, PS4, PM1, PP1, PP3_Mod | Pathogenic | 0.997 | 401 | This work |
| chr19:g.38942483G>T | c.1202G>T | p.(Arg401Leu) | 0.965 | PS4_Mod, PM1_Sup, PM5, PP1, PP3_Mod | Likely Pathogenic | 0.975 | 401 | This work |
| chr19:g.38943625C>T | c.1411C>T | p.(Arg471Cys) | 0.578 | PM1 | VUS | 0.325 | 471 | This work |
| chr19:g.38943636G>T | c.1422G>T | p.(Gln474His) | 0.835 | PM1 | VUS | 0.325 | 474 | This work |
| chr19:g.38945887A>G | c.1453A>G | p.(Met485Val) | 0.551 | PM1 | VUS | 0.325 | 485 | PMID: 33767344 |
| chr19:g.38945893C>G | c.1459C>G | p.(Leu487Val) | 0.668 | PS4_Sup, PM1 | VUS | 0.500 | 487 | This work |
| chr19:g.38945894T>C | c.1460T>C | p.(Leu487Pro) | 0.962 | PS4_Sup, PM1, PP3_Mod | VUS | 0.812 | 487 | This work |
| chr19:g.38945909G>A | c.1475G>A | p.(Arg492His) | 0.81 | PM1 | VUS | 0.325 | 492 | This work |
| chr19:g.38945987T>C | c.1553T>C | p.(Val518Ala) | 0.738 | PM1 | VUS | 0.325 | 518 | This work |
| chr19:g.38945999A>C | c.1565A>C | p.(Tyr522Ser) | 0.942 | PS3, PS4_Sup, PM1, PP1, PP3_Mod | Pathogenic | 0.994 | 522 | PMID: 33767344 |
| chr19:g.38945999A>G | c.1565A>G | p.(Tyr522Cys) | 0.949 | PS4_Mod, PM1_Sup, PM5, PP3_Mod | Likely Pathogenic | 0.949 | 522 | This work |
| chr19:g.38946103G>A | c.1589G>A | p.(Arg530His) | 0.93 | PS4_Sup, PM1, PP3_Mod | VUS | 0.812 | 530 | PMID: 33767344 |
| chr19:g.38946111C>A | c.1597C>A | p.(Arg533Ser) | 0.86 | PM1, PP3_Mod | VUS | 0.675 | 533 | PMID: 33767344 |
| chr19:g.38946111C>T | c.1597C>T | p.(Arg533Cys) | 0.918 | PS3_Mod, PS4_Sup, PM1, PP1_St, PP3_Mod | Pathogenic | 0.997 | 533 | PMID: 33767344 |
| chr19:g.38946112G>A | c.1598G>A | p.(Arg533His) | 0.824 | PS3_Mod, PM1 | VUS | 0.675 | 533 | PMID: 33767344 |
| chr19:g.38946129T>C | c.1615T>C | p.(Phe539Leu) | 0.934 | PS4_Mod, PM1, PP1, PP3_Mod | Likely Pathogenic | 0.949 | 539 | This work |
| chr19:g.38946129T>G | c.1615T>G | p.(Phe539Val) | 0.937 | PS4_Mod, PM1_Sup, PM5_Sup, PP3_Mod | Likely Pathogenic | 0.900 | 539 | This work |
| chr19:g.38946144G>T | c.1630G>T | p.(Asp544Tyr) | 0.924 | PS4_Sup, PM1, PP1, PP3_Mod | Likely Pathogenic | 0.900 | 544 | This work |
| chr19:g.38946168C>T | c.1654C>T | p.(Arg552Trp) | 0.831 | PS3_Mod, PS4_Mod, PM1, PP1 | Likely Pathogenic | 0.949 | 552 | PMID: 33767344 |
| chr19:g.38948179G>C | c.1834G>C | p.(Ala612Pro) | 0.971 | PS4_Sup, PP1, PP3_Mod | VUS | 0.675 | 612 | This work |
| chr19:g.38948185C>T | c.1840C>T | p.(Arg614Cys) | 0.927 | PS3_Mod, PS4, PM5, PP1_St, PP3_Mod, BS2_Mod | Pathogenic | 0.999 | 614 | PMID: 33767344 |
| chr19:g.38948186G>T | c.1841G>T | p.(Arg614Leu) | 0.931 | PS3_Mod, PS4, PP1_St, PP3_Mod | Pathogenic | 0.999 | 614 | PMID: 33767344 |
| chr19:g.38948815G>C | c.2050G>C | p.(Gly684Arg) | 0.962 | PS4_Sup, PP3_Mod | VUS | 0.500 | 684 | This work |
| chr19:g.38948887G>A | c.2122G>A | p.(Asp708Asn) | 0.726 | None | VUS | 0.100 | 708 | PMID: 33767344 |
| chr19:g.38951101C>T | c.2447C>T | p.(Pro816Leu) | 0.758 | PS4_Sup, BS2_Mod, BS3_Sup | Likely Benign | 0.025 | 816 | This work |
| chr19:g.38951191C>T | c.2537C>T | p.(Ser846Leu) | 0.558 | PS4_Sup | VUS | 0.188 | 846 | This work |
| chr19:g.38954139G>A | c.2654G>A | p.(Arg885His) | 0.609 | None | VUS | 0.100 | 885 | This work |
| chr19:g.38955289G>A | c.2797G>A | p.(Ala933Thr) | 0.926 | BA1 | Benign | <0.001 | 933 | PMID: 33767344 |
| chr19:g.38956784G>A | c.2924G>A | p.(Arg975Gln) | 0.329 | PS4_Sup, BP4 | VUS | 0.100 | 975 | This work |

| | | | | | | | | |
|---|---|---|---|---|---|---|---|---|
| chr19:g.38956856G>A | c.2996G>A | p.(Arg999His) | 0.817 | None | VUS | 0.100 | 999 | PMID: 33767344 |
| chr19:g.38956955G>A | c.3095G>A | p.(Arg1032His) | 0.547 | None | VUS | 0.100 | 1032 | This work |
| chr19:g.38956987C>T | c.3127C>T | p.(Arg1043Cys) | 0.936 | PP3_Mod | VUS | 0.325 | 1043 | This work |
| chr19:g.38957026G>A | c.3166G>A | p.(Asp1056Asn) | 0.685 | PS4_Sup | VUS | 0.188 | 1056 | This work |
| chr19:g.38957026G>C | c.3166G>C | p.(Asp1056His) | 0.824 | PS4_Mod, PP1_St | Likely Pathogenic | 0.900 | 1056 | This work |
| chr19:g.38957032G>A | c.3172G>A | p.(Glu1058Lys) | 0.78 | PS4_Mod | VUS | 0.325 | 1058 | This work |
| chr19:g.38958295G>A | c.3224G>A | p.(Arg1075Gln) | 0.846 | PS4_Sup, BS2_Mod | Likely Benign | 0.051 | 1075 | This work |
| chr19:g.38959642C>T | c.3418C>T | p.(Arg1140Cys) | 0.807 | None | VUS | 0.100 | 1140 | This work |
| chr19:g.38959751C>T | c.3527C>T | p.(Thr1176Ile) | 0.162 | BP4 | VUS | 0.100 | 1176 | This work |
| chr19:g.38960044A>C | c.3656A>C | p.(Gln1219Pro) | 0.891 | PP3_Mod | VUS | 0.325 | 1219 | This work |
| chr19:g.38960055G>A | c.3667G>A | p.(Glu1223Lys) | 0.831 | None | VUS | 0.100 | 1223 | This work |
| chr19:g.38964275A>G | c.4024A>G | p.(Ser1342Gly) | 0.293 | BA1 | Benign | <0.001 | 1342 | PMID: 33767344 |
| chr19:g.38965975A>G | c.4178A>G | p.(Lys1393Arg) | 0.555 | BA1 | Benign | <0.001 | 1393 | This work |
| chr19:g.38968456A>G | c.4400A>G | p.(Lys1467Arg) | 0.371 | BP4 | VUS | 0.100 | 1467 | PMID: 33767344 |
| chr19:g.38973933A>G | c.4711A>G | p.(Ile1571Val) | 0.56 | BS1 | Likely Benign | 0.006 | 1571 | PMID: 33767344 |
| chr19:g.38973969C>T | c.4747C>T | p.(Arg1583Cys) | 0.586 | None | VUS | 0.100 | 1583 | This work |
| chr19:g.38973985C>T | c.4763C>T | p.(Pro1588Leu) | 0.735 | None | VUS | 0.100 | 1588 | This work |
| chr19:g.38973997C>T | c.4775C>T | p.(Pro1592Leu) | 0.927 | PP3_Mod | VUS | 0.325 | 1592 | This work |
| chr19:g.38976319T>C | c.5024T>C | p.(Leu1675Pro) | 0.798 | PS4_Mod | VUS | 0.325 | 1675 | This work |
| chr19:g.38976328A>G | c.5033A>G | p.(Asn1678Ser) | 0.371 | BP4 | VUS | 0.100 | 1678 | This work |
| chr19:g.38976331G>A | c.5036G>A | p.(Arg1679His) | 0.918 | PP3_Mod, BS1, BS2_Mod | Likely Benign | 0.006 | 1679 | PMID: 33767344 |
| chr19:g.38976427A>G | c.5132A>G | p.(Tyr1711Cys) | 0.805 | PS4_Sup, PP1 | VUS | 0.325 | 1711 | This work |
| chr19:g.38976478C>T | c.5183C>T | p.(Ser1728Phe) | 0.477 | PS4, PP1_Mod, BS2_Mod, BP4 | VUS | 0.500 | 1728 | PMID: 33767344 |
| chr19:g.38976481T>G | c.5186T>G | p.(Met1729Arg) | 0.712 | PS4_Sup | VUS | 0.188 | 1729 | This work |
| chr19:g.38976612C>T | c.5317C>T | p.(Pro1773Ser) | 0.46 | BA1 | Benign | <0.001 | 1773 | PMID: 33767344 |
| chr19:g.38976636T>C | c.5341T>C | p.(Cys1781Arg) | 0.298 | PS4_Sup, BP4 | VUS | 0.100 | 1781 | This work |
| chr19:g.38976655C>T | c.5360C>T | p.(Pro1787Leu) | 0.055 | BA1 | Benign | <0.001 | 1787 | PMID: 33767344 |
| chr19:g.38976735A>G | c.5440A>G | p.(Met1814Val) | 0.486 | PS4_Sup, BP4 | VUS | 0.100 | 1814 | This work |
| chr19:g.38976736T>A | c.5441T>A | p.(Met1814Lys) | 0.81 | None | VUS | 0.100 | 1814 | This work |
| chr19:g.38980791C>T | c.5890C>T | p.(Arg1964Cys) | 0.232 | BP4 | VUS | 0.100 | 1964 | This work |
| chr19:g.38981282A>C | c.6037A>C | p.(Lys2013Gln) | 0.397 | PS4_Sup, BP4 | VUS | 0.100 | 2013 | This work |
| chr19:g.38983180G>T | c.6178G>T | p.(Gly2060Cys) | 0.172 | BA1 | Benign | <0.001 | 2060 | PMID: 33767344 |
| chr19:g.38985019T>A | c.6302T>A | p.(Met2101Lys) | 0.392 | PM1, BP4 | VUS | 0.188 | 2101 | This work |
| chr19:g.38985021G>C | c.6304G>C | p.(Val2102Leu) | 0.515 | PM1 | VUS | 0.325 | 2102 | This work |
| chr19:g.38985066G>C | c.6349G>C | p.(Val2117Leu) | 0.923 | PS4_Mod, PM1, PP3_Mod | Likely Pathogenic | 0.900 | 2117 | This work |
| chr19:g.38985094G>A | c.6377G>A | p.(Arg2126Gln) | 0.796 | PS4_Sup, PM1 | VUS | 0.500 | 2126 | This work |
| chr19:g.38985104C>G | c.6387C>G | p.(Asp2129Glu) | 0.683 | PS4_Mod, PM1, PP1_Mod | Likely Pathogenic | 0.900 | 2129 | This work |
| chr19:g.38985105G>A | c.6388G>A | p.(Gly2130Arg) | 0.894 | PM1, PP3_Mod | VUS | 0.675 | 2130 | This work |
| chr19:g.38985195G>A | c.6478G>A | p.(Gly2160Ser) | 0.635 | PM1 | VUS | 0.325 | 2160 | PMID: 33767344 |
| chr19:g.38985204C>T | c.6487C>T | p.(Arg2163Cys) | 0.956 | PS3_Mod, PS4_Mod, PM1_Sup, PM5, PP1_St, PP3_Mod, BS2_Mod | Pathogenic | 0.997 | 2163 | PMID: 33767344 |
| chr19:g.38985205G>A | c.6488G>A | p.(Arg2163His) | 0.933 | PS3_Mod, PS4, PM1, PP1_St, PP3_Mod, BS2 | Pathogenic | 0.994 | 2163 | PMID: 33767344 |
| chr19:g.38985205G>C | c.6488G>C | p.(Arg2163Pro) | 0.947 | PS4_Mod, PM1_Sup, PM5, PP1, PP3_Mod | Likely Pathogenic | 0.975 | 2163 | This work |
| chr19:g.38985205G>T | c.6488G>T | p.(Arg2163Leu) | 0.926 | PM1_Sup, PM5, PP3_Mod | VUS | 0.812 | 2163 | This work |
| chr19:g.38985219G>A | c.6502G>A | p.(Val2168Met) | 0.896 | PS3_Mod, PS4, PM1, PP1_St, PP3_Mod | Pathogenic | >0.999 | 2168 | PMID: 33767344 |
| chr19:g.38985261A>T | c.6544A>T | p.(Ile2182Phe) | 0.76 | PS4_Sup, PM1 | VUS | 0.500 | 2182 | This work |
| chr19:g.38985265G>A | c.6548G>A | p.(Gly2183Glu) | 0.786 | PS4_Sup, PM1 | VUS | 0.500 | 2183 | This work |
| chr19:g.38986905C>T | c.6599C>T | p.(Ala2200Val) | 0.564 | PS4_Sup, PM1, PP1_Mod | VUS | 0.812 | 2200 | This work |
| chr19:g.38986918C>G | c.6612C>G | p.(His2204Gln) | 0.739 | PS4_Mod, PM1, PP1_Mod | Likely Pathogenic | 0.900 | 2204 | This work |
| chr19:g.38986923C>G | c.6617C>G | p.(Thr2206Arg) | 0.968 | PS3_Mod, PS4_Mod, PM1, PP1_Mod, PP3_Mod | Pathogenic | 0.994 | 2206 | PMID: 33767344 |
| chr19:g.38986923C>T | c.6617C>T | p.(Thr2206Met) | 0.997 | PS3_Mod, PS4, PM1_Sup, PM5, PP1_St, PP3_Mod, BS2 | Pathogenic | 0.997 | 2206 | PMID: 33767344 |
| chr19:g.38986934G>T | c.6628G>T | p.(Val2210Phe) | 0.961 | PS4_Mod, PM1, PP3_Mod | Likely Pathogenic | 0.900 | 2210 | This work |
| chr19:g.38986941T>A | c.6635T>A | p.(Val2212Asp) | 0.973 | PM1, PP3_Mod | VUS | 0.675 | 2212 | This work |
| chr19:g.38986946G>A | c.6640G>A | p.(Val2214Ile) | 0.798 | PM1 | VUS | 0.325 | 2214 | This work |
| chr19:g.38987055C>T | c.6670C>T | p.(Arg2224Cys) | 0.728 | PM1, BS1 | Likely Benign | 0.025 | 2224 | This work |
| chr19:g.38987056G>A | c.6671G>A | p.(Arg2224His) | 0.605 | PM1 | VUS | 0.325 | 2224 | PMID: 33767344 |
| chr19:g.38987095G>A | c.6710G>A | p.(Cys2237Tyr) | 0.939 | PS4_Sup, PM1, PP3_Mod | VUS | 0.812 | 2237 | This work |
| chr19:g.38987127C>T | c.6742C>T | p.(Arg2248Cys) | 0.694 | PM1 | VUS | 0.325 | 2248 | This work |
| chr19:g.38987128G>A | c.6743G>A | p.(Arg2248His) | 0.654 | PM1 | VUS | 0.325 | 2248 | This work |
| chr19:g.38987142C>T | c.6757C>T | p.(His2253Tyr) | 0.973 | PS4_Sup, PM1, PP3_Mod | VUS | 0.812 | 2253 | This work |
| chr19:g.38987541G>A | c.6838G>A | p.(Val2280Ile) | 0.641 | PS4_Mod, PM1 | VUS | 0.675 | 2280 | This work |
| chr19:g.38987550A>C | c.6847A>C | p.(Asn2283His) | 0.943 | PS4_Sup, PM1, PP3_Mod | VUS | 0.812 | 2283 | This work |
| chr19:g.38989817A>G | c.6961A>G | p.(Ile2321Val) | 0.712 | PM1, BS1, BS2, BS3_Sup | Benign | <0.001 | 2321 | This work |
| chr19:g.38989863G>A | c.7007G>A | p.(Arg2336His) | 0.903 | PS3_Mod, PS4, PM1, PP1_St, PP3_Mod | Pathogenic | >0.999 | 2336 | PMID: 33767344 |
| chr19:g.38989874T>C | c.7018T>C | p.(Phe2340Leu) | 0.789 | PS4_Sup, PM1 | VUS | 0.500 | 2340 | This work |

| | | | | | | | | |
|---|---|---|---|---|---|---|---|---|
| chr19:g.38989881A>G | c.7025A>G | p.(Asn2342Ser) | 0.639 | PS3_Mod, PM1, PP1, BS1, BS2 | Likely Benign | 0.012 | 2342 | This work |
| chr19:g.38990279G>C | c.7032G>C | p.(Glu2344Asp) | 0.657 | PM1 | VUS | 0.325 | 2344 | This work |
| chr19:g.38990282C>A | c.7035C>A | p.(Ser2345Arg) | 0.747 | PS3_Mod, PS4_Mod, PM1 | Likely Pathogenic | 0.900 | 2345 | This work |
| chr19:g.38990283G>A | c.7036G>A | p.(Val2346Met) | 0.944 | PS4_Mod, PM1, PP3_Mod | Likely Pathogenic | 0.900 | 2346 | This work |
| chr19:g.38990289_ 38990291del | c.7042_7044del | p.(Glu2348del) | NA | PS3_Mod, PS4_Mod, PM1, PP1 | Likely Pathogenic | 0.949 | 2348 | PMID: 33767344 |
| chr19:g.38990290A>G | c.7043A>G | p.(Glu2348Gly) | 0.964 | PS4_Mod, PM1, PP3_Mod | Likely Pathogenic | 0.900 | 2348 | This work |
| chr19:g.38990295G>A | c.7048G>A | p.(Ala2350Thr) | 0.952 | PS3_Mod, PS4, PM1, PP1_St, PP3_Mod | Pathogenic | >0.999 | 2350 | PMID: 33767344 |
| chr19:g.38990307G>A | c.7060G>A | p.(Val2354Met) | 0.867 | PS4_Mod, PM1, PP1_Mod, PP3_Mod | Likely Pathogenic | 0.975 | 2354 | This work |
| chr19:g.38990310C>T | c.7063C>T | p.(Arg2355Trp) | 0.861 | PS3_Mod, PS4, PM1, PP1_St, PP3_Mod | Pathogenic | >0.999 | 2355 | PMID: 33767344 |
| chr19:g.38990320T>A | c.7073T>A | p.(Ile2358Asn) | 0.867 | PM1, PP3_Mod | VUS | 0.675 | 2358 | PMID: 33767344 |
| chr19:g.38990322C>T | c.7075C>T | p.(Arg2359Trp) | 0.93 | PM1_Sup, PM5_Sup, PP3_Mod | VUS | 0.675 | 2359 | This work |
| chr19:g.38990323G>A | c.7076G>A | p.(Arg2359Gln) | 0.873 | PS4_Mod, PM1, PP3_Mod | Likely Pathogenic | 0.900 | 2359 | This work |
| chr19:g.38990331G>A | c.7084G>A | p.(Glu2362Lys) | 0.885 | PS4_Mod, PM1, PP3_Mod | Likely Pathogenic | 0.900 | 2362 | This work |
| chr19:g.38990332A>G | c.7085A>G | p.(Glu2362Gly) | 0.944 | PS4_Sup, PM1_Sup, PM5_Sup, PP3_Mod | VUS | 0.812 | 2362 | This work |
| chr19:g.38990336C>G | c.7089C>G | p.(Cys2363Trp) | 0.78 | PS4_Mod, PM1 | VUS | 0.675 | 2363 | This work |
| chr19:g.38990337T>G | c.7090T>G | p.(Phe2364Val) | 0.85 | PS4_Mod, PM1, PP3_Mod | Likely Pathogenic | 0.900 | 2364 | This work |
| chr19:g.38990344C>G | c.7097C>G | p.(Pro2366Arg) | 0.88 | PS4_Sup, PM1, PP3_Mod | VUS | 0.812 | 2366 | This work |
| chr19:g.38990346G>A | c.7099G>A | p.(Ala2367Thr) | 0.899 | PS4_Sup, PM1, PP3_Mod | VUS | 0.812 | 2367 | This work |
| chr19:g.38990359A>G | c.7112A>G | p.(Glu2371Gly) | 0.891 | PS4_Sup, PM1, PP3_Mod | VUS | 0.812 | 2371 | This work |
| chr19:g.38990370G>A | c.7123G>A | p.(Gly2375Arg) | 0.9 | PS4_Sup, PM1_Sup, PM5, PP3_Mod | Likely Pathogenic | 0.900 | 2375 | PMID: 33767344 |
| chr19:g.38990371G>C | c.7124G>C | p.(Gly2375Ala) | 0.905 | PS3_Mod, PS4_Mod, PM1, PP1_Mod, PP3_Mod | Pathogenic | 0.994 | 2375 | PMID: 33767344 |
| chr19:g.38990446A>G | c.7199A>G | p.(Asp2400Gly) | 0.548 | PM1 | VUS | 0.325 | 2400 | This work |
| chr19:g.38990457G>A | c.7210G>A | p.(Glu2404Lys) | 0.676 | PM1, PP1 | VUS | 0.500 | 2404 | This work |
| chr19:g.38990615G>A | c.7282G>A | p.(Ala2428Thr) | 0.85 | PS3_Mod, PS4_Mod, PM1, PP3_Mod | Likely Pathogenic | 0.975 | 2428 | PMID: 33767344 |
| chr19:g.38990624G>A | c.7291G>A | p.(Asp2431Asn) | 0.888 | PS4_Mod, PM1, PP3_Mod | Likely Pathogenic | 0.900 | 2431 | This work |
| chr19:g.38990624G>T | c.7291G>T | p.(Asp2431Tyr) | 0.964 | PS3_Mod, PS4_Mod, PM1_Sup, PM5_Sup, PP1, PP3_Mod | Likely Pathogenic | 0.988 | 2431 | This work |
| chr19:g.38990625A>T | c.7292A>T | p.(Asp2431Val) | 0.952 | PS4_Sup, PM1_Sup, PM5_Sup, PP3_Mod | VUS | 0.812 | 2431 | This work |
| chr19:g.38990633G>A | c.7300G>A | p.(Gly2434Arg) | 0.965 | PS3, PS4, PM1, PP1_St, PP3_Mod, BS2 | Pathogenic | 0.999 | 2434 | PMID: 33767344 |
| chr19:g.38990637G>A | c.7304G>A | p.(Arg2435His) | 0.944 | PS3_Mod, PS4, PM1, PP1_St, PP3_Mod | Pathogenic | >0.999 | 2435 | PMID: 33767344 |
| chr19:g.38990637G>T | c.7304G>T | p.(Arg2435Leu) | 0.948 | PS4_Mod, PM1_Sup, PM5, PP3_Mod | Likely Pathogenic | 0.949 | 2435 | This work |
| chr19:g.38990640G>A | c.7307G>A | p.(Cys2436Tyr) | 0.921 | PS4_Sup, PM1, PP3_Mod | VUS | 0.812 | 2436 | This work |
| chr19:g.38990643C>T | c.7310C>T | p.(Ala2437Val) | 0.917 | PS4_Mod, PM1, PP3_Mod | Likely Pathogenic | 0.900 | 2437 | This work |
| chr19:g.38990650G>C | c.7317G>C | p.(Glu2439Asp) | 0.734 | PM1 | VUS | 0.325 | 2439 | This work |
| chr19:g.38991276C>T | c.7354C>T | p.(Arg2452Trp) | 0.828 | PS3_Mod, PS4_Mod, PM1, PP1_Mod | Likely Pathogenic | 0.975 | 2452 | PMID: 33767344 |
| chr19:g.38991277G>A | c.7355G>A | p.(Arg2452Gln) | 0.928 | PM1, PP3_Mod | VUS | 0.675 | 2452 | This work |
| chr19:g.38991277G>C | c.7355G>C | p.(Arg2452Pro) | 0.953 | PM1_Sup, PM5_Sup, PP3_Mod | VUS | 0.675 | 2452 | This work |
| chr19:g.38991280T>C | c.7358T>C | p.(Ile2453Thr) | 0.891 | PS2_PM6_Mod, PS4_Sup, PM1, PP3_Mod | Likely Pathogenic | 0.949 | 2453 | This work |
| chr19:g.38991282C>T | c.7360C>T | p.(Arg2454Cys) | 0.913 | PS3_Mod, PS4_Mod, PM1_Sup, PM5, PP1, PP3_Mod | Pathogenic | 0.994 | 2454 | PMID: 33767344 |
| chr19:g.38991283G>A | c.7361G>A | p.(Arg2454His) | 0.923 | PS3_Mod, PS4, PM1, PP1_St, PP3_Mod, BS2_Mod | Pathogenic | 0.999 | 2454 | PMID: 33767344 |
| chr19:g.38991294C>T | c.7372C>T | p.(Arg2458Cys) | 0.922 | PS3_Mod, PS4_Mod, PM1_Sup, PM5, PP3_Mod | Likely Pathogenic | 0.988 | 2458 | PMID: 33767344 |
| chr19:g.38991295G>A | c.7373G>A | p.(Arg2458His) | 0.959 | PS3_Mod, PS4, PM1, PP1_St, PP3_Mod | Pathogenic | >0.999 | 2458 | PMID: 33767344 |
| chr19:g.38991295G>T | c.7373G>T | p.(Arg2458Leu) | 0.968 | PS4_Mod, PM1_Sup, PM5, PP3_Mod | Likely Pathogenic | 0.949 | 2458 | This work |
| chr19:g.38991307C>T | c.7385C>T | p.(Pro2462Leu) | 0.883 | PP3_Mod | VUS | 0.325 | 2462 | PMID: 33767344 |
| chr19:g.38991503C>T | c.7487C>T | p.(Pro2496Leu) | 0.96 | PP3_Mod | VUS | 0.325 | 2496 | This work |
| chr19:g.38991538C>T | c.7522C>T | p.(Arg2508Cys) | 0.861 | PS3_Mod, PS4_Mod, PM5_Sup, PP3_Mod | Likely Pathogenic | 0.949 | 2508 | PMID: 33767344 |
| chr19:g.38991539G>A | c.7523G>A | p.(Arg2508His) | 0.898 | PS2/PM6_Sup, PS3_Mod, PS4, PP3_Mod | Likely Pathogenic | 0.988 | 2508 | PMID: 33767344 |
| chr19:g.38991544T>C | c.7528T>C | p.(Tyr2510His) | 0.922 | PP3_Mod | VUS | 0.325 | 2510 | This work |
| chr19:g.38993292A>G | c.7760A>G | p.(Tyr2587Cys) | 0.911 | PP3_Mod | VUS | 0.325 | 2587 | This work |
| chr19:g.38993303C>G | c.7771C>G | p.(Arg2591Gly) | 0.642 | None | VUS | 0.100 | 2591 | This work |
| chr19:g.38993303C>T | c.7771C>T | p.(Arg2591Trp) | 0.697 | None | VUS | 0.100 | 2591 | This work |
| chr19:g.38993309C>G | c.7777C>G | p.(Arg2593Gly) | 0.769 | None | VUS | 0.100 | 2593 | This work |
| chr19:g.38993310G>A | c.7778G>A | p.(Arg2593His) | 0.825 | PS4_Sup | VUS | 0.188 | 2593 | This work |
| chr19:g.38993319C>T | c.7787C>T | p.(Thr2596Ile) | 0.822 | None | VUS | 0.100 | 2596 | This work |
| chr19:g.38993348T>A | c.7816T>A | p.(Cys2606Ser) | 0.818 | BS2_Mod | Likely Benign | 0.025 | 2606 | This work |
| chr19:g.38993563G>A | c.7879G>A | p.(Val2627Met) | 0.808 | PS4_Mod | VUS | 0.325 | 2627 | This work |
| chr19:g.38993563G>C | c.7879G>C | p.(Val2627Leu) | 0.894 | PS4_Mod, PP1_St, PP3_Mod | Likely Pathogenic | 0.975 | 2627 | This work |
| chr19:g.38994938G>A | c.8005G>A | p.(Glu2669Lys) | 0.82 | None | VUS | 0.100 | 2669 | This work |
| chr19:g.38994959C>T | c.8026C>T | p.(Arg2676Trp) | 0.601 | PS4_Mod, PP1_St | Likely Pathogenic | 0.900 | 2676 | This work |
| chr19:g.38994987C>T | c.8054C>T | p.(Ser2685Phe) | 0.777 | PS4_Sup | VUS | 0.188 | 2685 | This work |
| chr19:g.38995508G>C | c.8188G>C | p.(Asp2730His) | 0.881 | PP3_Mod | VUS | 0.325 | 2730 | This work |
| chr19:g.38995509A>G | c.8189A>G | p.(Asp2730Gly) | 0.722 | PS4_Sup, PP1_St | VUS | 0.812 | 2730 | This work |
| chr19:g.38995518G>A | c.8198G>A | p.(Gly2733Asp) | 0.923 | PS4_Mod, PP3_Mod | VUS | 0.675 | 2733 | This work |
| chr19:g.38995701G>A | c.8290G>A | p.(Glu2764Lys) | 0.842 | None | VUS | 0.100 | 2764 | This work |
| chr19:g.38995965C>T | c.8327C>T | p.(Ser2776Phe) | 0.693 | BS1 | Likely Benign | 0.006 | 2776 | This work |
| chr19:g.38995998C>G | c.8360C>G | p.(Thr2787Ser) | 0.466 | BA1 | Benign | <0.001 | 2787 | This work |
| chr19:g.38996563C>T | c.8518C>T | p.(Arg2840Trp) | 0.659 | None | VUS | 0.100 | 2840 | This work |
| chr19:g.38996572T>C | c.8527T>C | p.(Ser2843Pro) | 0.761 | PS4_Sup | VUS | 0.188 | 2843 | This work |

| | | | | | | | | |
|---|---|---|---|---|---|---|---|---|
| chr19:g.38997001T>A | c.8600T>A | p.(Leu2867Gln) | 0.923 | PP3_Mod | VUS | 0.325 | 2867 | This work |
| chr19:g.38997132G>A | c.8638G>A | p.(Glu2880Lys) | 0.901 | PS4_Sup, PP3_Mod | VUS | 0.500 | 2880 | This work |
| chr19:g.38997148C>G | c.8654C>G | p.(Thr2885Arg) | 0.656 | None | VUS | 0.100 | 2885 | This work |
| chr19:g.38997505C>T | c.8729C>T | p.(Thr2910Met) | 0.779 | PS4_Sup | VUS | 0.188 | 2910 | This work |
| chr19:g.38998461C>T | c.8926C>T | p.(His2976Tyr) | 0.201 | PS4_Sup, BP4 | VUS | 0.100 | 2976 | This work |
| chr19:g.39002230G>A | c.9152G>A | p.(Arg3051His) | 0.581 | None | VUS | 0.100 | 3051 | This work |
| chr19:g.39002919G>A | c.9268G>A | p.(Ala3090Thr) | 0.454 | PS4_Sup, BP4 | VUS | 0.100 | 3090 | This work |
| chr19:g.39002961G>A | c.9310G>A | p.(Glu3104Lys) | 0.868 | PS3_Mod, PS4_Mod, PP1_Mod, PP3_Mod | Likely Pathogenic | 0.975 | 3104 | PMID: 33767344 |
| chr19:g.39003007G>A | c.9356G>A | p.(Arg3119His) | 0.711 | None | VUS | 0.100 | 3119 | This work |
| chr19:g.39005692C>T | c.9499C>T | p.(Arg3167Term) | | None | VUS | 0.100 | 3167 | This work |
| chr19:g.39006807A>G | c.9635A>G | p.(Glu3212Gly) | 0.624 | None | VUS | 0.100 | 3212 | This work |
| chr19:g.39006821T>C | c.9649T>C | p.(Ser3217Pro) | 0.918 | PP3_Mod | VUS | 0.325 | 3217 | This work |
| chr19:g.39006824G>A | c.9652G>A | p.(Val3218Met) | 0.608 | PS4_Sup | VUS | 0.188 | 3218 | This work |
| chr19:g.39006848G>C | c.9676G>C | p.(Glu3226Gln) | 0.62 | PS4_Sup | VUS | 0.188 | 3226 | This work |
| chr19:g.39008071T>C | c.9758T>C | p.(Ile3253Thr) | 0.845 | None | VUS | 0.100 | 3253 | PMID: 33767344 |
| chr19:g.39008110T>C | c.9797T>C | p.(Met3266Thr) | 0.742 | PS4_Sup | VUS | 0.188 | 3266 | This work |
| chr19:g.39008161G>A | c.9848G>A | p.(Arg3283Gln) | 0.527 | None | VUS | 0.100 | 3283 | This work |
| chr19:g.39008163T>A | c.9850T>A | p.(Trp3284Arg) | 0.873 | PP3_Mod | VUS | 0.325 | 3284 | This work |
| chr19:g.39008181G>A | c.9868G>A | p.(Glu3290Lys) | 0.848 | None | VUS | 0.100 | 3290 | This work |
| chr19:g.39009877C>T | c.10042C>T | p.(Arg3348Cys) | 0.784 | PS4_Sup | VUS | 0.188 | 3348 | PMID: 33767344 |
| chr19:g.39009878G>A | c.10043G>A | p.(Arg3348His) | 0.705 | PS4_Sup | VUS | 0.188 | 3348 | This work |
| chr19:g.39009932G>A | c.10097G>A | p.(Arg3366His) | 0.68 | BS1 | Likely Benign | 0.006 | 3366 | PMID: 33767344 |
| chr19:g.39009935A>G | c.10100A>G | p.(Lys3367Arg) | 0.632 | BS3_Sup | Likely Benign | 0.051 | 3367 | This work |
| chr19:g.39010064C>A | c.10229C>A | p.(Pro3410Gln) | 0.934 | PP3_Mod | VUS | 0.325 | 3410 | This work |
| chr19:g.39010072A>T | c.10237A>T | p.(Ile3413Phe) | 0.914 | PS4_Sup, PP3_Mod | VUS | 0.500 | 3413 | This work |
| chr19:g.39010087A>G | c.10252A>G | p.(Asn3418Asp) | 0.535 | PS4_Mod | VUS | 0.325 | 3418 | This work |
| chr19:g.39016072C>T | c.10556C>T | p.(Pro3519Leu) | 0.809 | None | VUS | 0.100 | 3519 | PMID: 33767344 |
| chr19:g.39016132G>A | c.10616G>A | p.(Arg3539His) | 0.877 | PP3_Mod, BS1, BS2 | Likely Benign | 0.001 | 3539 | This work |
| chr19:g.39018347G>C | c.10747G>C | p.(Glu3583Gln) | 0.32 | BA1 | Benign | <0.001 | 3583 | PMID: 33767344 |
| chr19:g.39018350G>C | c.10750G>C | p.(Glu3584Gln) | 0.798 | None | VUS | 0.100 | 3584 | This work |
| chr19:g.39019012G>T | c.10891G>T | p.(Ala3631Ser) | 0.85 | PS4_Sup, PP3_Mod | VUS | 0.500 | 3631 | This work |
| chr19:g.39019642G>C | c.11086G>C | p.(Asp3696His) | 0.809 | PS4_Sup | VUS | 0.188 | 3696 | This work |
| chr19:g.39019676G>T | c.11120G>T | p.(Arg3707Leu) | 0.906 | PP3_Mod | VUS | 0.325 | 3707 | This work |
| chr19:g.39019682C>T | c.11126C>T | p.(Ala3709Val) | 0.793 | PS4_Sup | VUS | 0.188 | 3709 | This work |
| chr19:g.39019688C>T | c.11132C>T | p.(Thr3711Met) | 0.637 | PS4_Sup, PP1 | VUS | 0.325 | 3711 | This work |
| chr19:g.39025366C>G | c.11266C>G | p.(Gln3756Glu) | 0.326 | BA1 | Benign | <0.001 | 3756 | PMID: 33767344 |
| chr19:g.39025414C>T | c.11314C>T | p.(Arg3772Trp) | 0.939 | PS4_Sup, PM5_Sup, PP3_Mod | VUS | 0.675 | 3772 | PMID: 33767344 |
| chr19:g.39025415G>A | c.11315G>A | p.(Arg3772Gln) | 0.888 | PS4_Mod, PP1_St, PP3_Mod, BS2_Mod | Likely Pathogenic | 0.900 | 3772 | PMID: 33767344 |
| chr19:g.39025837G>A | c.11416G>A | p.(Gly3806Arg) | 0.933 | PS4_Sup, PP1, PP3_Mod | VUS | 0.675 | 3806 | This work |
| chr19:g.39026638G>A | c.11518G>A | p.(Val3840Ile) | 0.648 | None | VUS | 0.100 | 3840 | This work |
| chr19:g.39034005G>A | c.11708G>A | p.(Arg3903Gln) | 0.97 | PS4, PP3_Mod | Likely Pathogenic | 0.900 | 3903 | This work |
| chr19:g.39034020A>T | c.11723A>T | p.(Asn3908Ile) | 0.949 | PS4_Sup, PP3_Mod | VUS | 0.500 | 3908 | This work |
| chr19:g.39034045T>G | c.11748T>G | p.(Ile3916Met) | 0.765 | None | VUS | 0.100 | 3916 | This work |
| chr19:g.39034191A>G | c.11798A>G | p.(Tyr3933Cys) | 0.983 | PP3_Mod, BS1 | Likely Benign | 0.025 | 3933 | PMID: 33767344 |
| chr19:g.39034206G>A | c.11813G>A | p.(Gly3938Asp) | 0.799 | None | VUS | 0.100 | 3938 | This work |
| chr19:g.39034450C>T | c.11947C>T | p.(Arg3983Cys) | 0.931 | PS2_PM6_Mod, PS4_Sup, PP3_Mod, BS3_Sup | VUS | 0.675 | 3983 | PMID: 33767344 |
| chr19:g.39034456T>C | c.11953T>C | p.(Trp3985Arg) | 0.941 | PS4_Mod, PP3_Mod | VUS | 0.675 | 3985 | This work |
| chr19:g.39034461C>G | c.11958C>G | p.(Asp3986Glu) | 0.763 | PS4, PP1, BS3_Sup | VUS | 0.675 | 3986 | PMID: 33767344 |
| chr19:g.39034472G>T | c.11969G>T | p.(Gly3990Val) | 0.75 | PS3_Mod, PS4, PP1_St | Pathogenic | 0.994 | 3990 | PMID: 33767344 |
| chr19:g.39037100G>A | c.12028G>A | p.(Glu4010Lys) | 0.178 | PS4_Sup, BP4 | VUS | 0.100 | 4010 | This work |
| chr19:g.39037136A>G | c.12064A>G | p.(Met4022Val) | 0.491 | BP4 | VUS | 0.100 | 4022 | This work |
| chr19:g.39038893A>T | c.12115A>T | p.(Ile4039Phe) | 0.621 | PS4_Sup | VUS | 0.188 | 4039 | This work |
| chr19:g.39038899C>T | c.12121C>T | p.(Arg4041Trp) | 0.435 | BP4 | VUS | 0.100 | 4041 | This work |
| chr19:g.39038927C>A | c.12149C>A | p.(Ser4050Tyr) | 0.942 | PS4_Mod, PP1_St, PP3_Mod | Likely Pathogenic | 0.975 | 4050 | This work |
| chr19:g.39039020C>T | c.12242C>T | p.(Thr4081Met) | 0.649 | None | VUS | 0.100 | 4081 | This work |
| chr19:g.39051780G>C | c.12310G>C | p.(Gly4104Arg) | 0.307 | BS2_Mod, BP4 | Likely Benign | 0.012 | 4104 | This work |
| chr19:g.39051825A>T | c.12355A>T | p.(Asn4119Tyr) | 0.813 | PS4_Sup | VUS | 0.188 | 4119 | This work |
| chr19:g.39051853C>T | c.12383C>T | p.(Ala4128Val) | 0.248 | PS4_Sup, BP4 | VUS | 0.100 | 4128 | This work |
| chr19:g.39051868A>G | c.12398A>G | p.(Glu4133Gly) | 0.881 | PS4_Sup, PP3_Mod | VUS | 0.500 | 4133 | This work |
| chr19:g.39051876C>A | c.12406C>A | p.(Arg4136Ser) | 0.783 | BS2_Mod | Likely Benign | 0.025 | 4136 | This work |
| chr19:g.39051883T>C | c.12413T>C | p.(Ile4138Thr) | 0.96 | PS4_Sup, PP3_Mod | VUS | 0.500 | 4138 | This work |
| chr19:g.39052002G>A | c.12532G>A | p.(Gly4178Ser) | 0.979 | PS4_Sup, PP3_Mod | VUS | 0.500 | 4178 | PMID: 33767344 |
| chr19:g.39052003G>T | c.12533G>T | p.(Gly4178Val) | 0.918 | PS4_Mod, PP3_Mod | VUS | 0.675 | 4178 | This work |
| chr19:g.39052023G>A | c.12553G>A | p.(Ala4185Thr) | 0.402 | BP4 | VUS | 0.100 | 4185 | This work |
| chr19:g.39055663T>G | c.12689T>G | p.(Met4230Arg) | 0.928 | PS4_Mod, PP3_Mod | VUS | 0.675 | 4230 | This work |
| chr19:g.39055674G>C | c.12700G>C | p.(Val4234Leu) | 0.89 | PS4_Mod, PP1_St, PP3_Mod | Likely Pathogenic | 0.975 | 4234 | This work |
| chr19:g.39055674G>T | c.12700G>T | p.(Val4234Leu) | 0.89 | PS1_Mod, PS4_Mod, PP1, PP3_Mod | Likely Pathogenic | 0.949 | 4234 | This work |
| chr19:g.39055822A>T | c.12848A>T | p.(Glu4283Val) | 0.561 | None | VUS | 0.100 | 4283 | This work |
| chr19:g.39055855C>T | c.12881C>T | p.(Thr4294Met) | 0.528 | BA1 | Benign | <0.001 | 4294 | PMID: 33767344 |
| chr19:g.39055858C>T | c.12884C>T | p.(Ala4295Val) | 0.152 | BS1, BS2_Mod, BP2, BP4 | Benign | <0.001 | 4295 | PMID: 33767344 |
| chr19:g.39057618A>G | c.13505A>G | p.(Glu4502Gly) | 0.532 | None | VUS | 0.100 | 4502 | PMID: 33767344 |
| chr19:g.39057626G>C | c.13513G>C | p.(Asp4505His) | 0.663 | BA1 | Benign | <0.001 | 4505 | PMID: 33767344 |
| chr19:g.39061259C>T | c.13672C>T | p.(Arg4558Trp) | 0.79 | PS4_Sup | VUS | 0.188 | 4558 | This work |

| | | | | | | | | |
|---|---|---|---|---|---|---|---|---|
| chr19:g.39061260G>A | c.13673G>A | p.(Arg4558Gln) | 0.957 | PP3_Mod | VUS | 0.325 | 4558 | PMID: 33767344 |
| chr19:g.39061289C>G | c.13702C>G | p.(Leu4568Val) | 0.775 | None | VUS | 0.100 | 4568 | PMID: 33767344 |
| chr19:g.39062672C>T | c.13760C>T | p.(Pro4587Leu) | 0.785 | None | VUS | 0.100 | 4587 | This work |
| chr19:g.39062825G>A | c.13913G>A | p.(Gly4638Asp) | 0.917 | PM1_Sup, PP3_Mod | VUS | 0.500 | 4638 | This work |
| chr19:g.39062830A>G | c.13918A>G | p.(Met4640Val) | 0.911 | PS4_Sup, PM1_Sup, PP3_Mod | VUS | 0.675 | 4640 | PMID: 33767344 |
| chr19:g.39062846G>A | c.13934G>A | p.(Arg4645Gln) | 0.567 | PM1_Sup | VUS | 0.188 | 4645 | This work |
| chr19:g.39062902T>C | c.13990T>C | p.(Cys4664Arg) | 0.936 | PS4_Sup, PM1_Sup, PP3_Mod | VUS | 0.675 | 4664 | This work |
| chr19:g.39062906_39062907delinsCT | c.13994_13995delTCinsCT | p.(Leu4665Pro) | | PS2_PM6_Sup, PM1_Sup | VUS | 0.325 | 4665 | This work |
| chr19:g.39063820C>T | c.14002C>T | p.(Pro4668Ser) | 0.888 | PM1_Sup, PP3_Mod | VUS | 0.500 | 4668 | This work |
| chr19:g.39063869T>C | c.14051T>C | p.(Phe4684Ser) | 0.982 | PS4_Sup, PM1_Sup, PP3_Mod | VUS | 0.675 | 4684 | This work |
| chr19:g.39063944C>T | c.14126C>T | p.(Thr4709Met) | 0.901 | PM1_Sup, PP3_Mod, BS3_Sup | VUS | 0.325 | 4709 | PMID: 33767344 |
| chr19:g.39066597G>A | c.14168G>A | p.(Arg4723His) | 0.79 | PM1_Sup, BS2_Mod | Likely Benign | 0.051 | 4723 | This work |
| chr19:g.39068571A>C | c.14186A>C | p.(His4729Pro) | 0.758 | PM1_Sup | VUS | 0.188 | 4729 | This work |
| chr19:g.39068582T>G | c.14197T>G | p.(Tyr4733Asp) | 0.842 | PS4_Sup, PM1_Sup | VUS | 0.325 | 4733 | This work |
| chr19:g.39068586G>A | c.14201G>A | p.(Gly4734Glu) | 0.871 | PS4_Sup, PM1_Sup, PP3_Mod | VUS | 0.675 | 4734 | This work |
| chr19:g.39068594C>T | c.14209C>T | p.(Arg4737Trp) | 0.882 | PS4_Mod, PM1_Sup, PM5_Sup, PP1, PP3_Mod | Likely Pathogenic | 0.949 | 4737 | This work |
| chr19:g.39068595G>A | c.14210G>A | p.(Arg4737Gln) | 0.89 | PS4, PM1_Sup, PP1_St, PP3_Mod, BS2_Mod | Likely Pathogenic | 0.988 | 4737 | This work |
| chr19:g.39068655G>A | c.14270G>A | p.(Arg4757His) | 0.452 | PM1_Sup, BP4 | VUS | 0.100 | 4757 | This work |
| chr19:g.39068845G>T | c.14364+1G>T | | | PS4_Sup, PM1_Sup | VUS | 0.325 | | This work |
| chr19:g.39070679_39070680delinsAA | c.14422_14423delinsAA | p.(Phe4808Asn) | | PM1_Sup | VUS | 0.188 | 4808 | This work |
| chr19:g.39070681C>A | c.14424C>A | p.(Phe4808Leu) | 0.923 | PS4_Sup, PM1_Sup, PP3_Mod | VUS | 0.675 | 4808 | This work |
| chr19:g.39070706A>T | c.14449A>T | p.(Ile4817Phe) | 0.911 | PS4_Sup, PM1_Sup, PP3_Mod | VUS | 0.675 | 4817 | This work |
| chr19:g.39070715G>A | c.14458G>A | p.(Gly4820Arg) | 0.835 | PS4_Sup, PM1_Sup | VUS | 0.325 | 4820 | This work |
| chr19:g.39070715G>T | c.14458G>T | p.(Gly4820Trp) | 0.918 | PS4_Sup, PM1_Sup, PP3_Mod | VUS | 0.675 | 4820 | This work |
| chr19:g.39070728T>C | c.14471T>C | p.(Leu4824Pro) | 0.984 | PS4_Mod, PM1_Sup, PP3_Mod | VUS | 0.812 | 4824 | This work |
| chr19:g.39070734C>T | c.14477C>T | p.(Thr4826Ile) | 0.977 | PS3, PS4, PM1_Sup, PP1_St, PP3_Mod | Pathogenic | >0.999 | 4826 | PMID: 33767344 |
| chr19:g.39070754C>T | c.14497C>T | p.(His4833Tyr) | 0.975 | PS3_Mod, PS4_Mod, PM1_Sup, PP1_Mod, PP3_Mod | Likely Pathogenic | 0.988 | 4833 | PMID: 33767344 |
| chr19:g.39070766C>G | c.14509C>G | p.(Gln4837Glu) | 0.914 | PS4_Sup, PM1_Sup, PP3_Mod | VUS | 0.675 | 4837 | This work |
| chr19:g.39070767del | c.14510delA | p.(Gln4837Argfs*3) | | PS4_Sup, BS2_Mod, BS3_Sup | Likely Benign | 0.025 | 4837 | This work |
| chr19:g.39071010C>G | c.14512C>G | p.(Leu4838Val) | 0.882 | PS3_Sup, PS4_Mod, PM1_Sup, PP3_Mod | Likely Pathogenic | 0.900 | 4838 | PMID: 33767344 |
| chr19:g.39071022G>A | c.14524G>A | p.(Val4842Met) | 0.933 | PM1_Sup, PP3_Mod | VUS | 0.500 | 4842 | This work |
| chr19:g.39071037G>C | c.14539G>C | p.(Val4847Leu) | 0.762 | PS4_Sup, PM1_Sup, PP1_St | Likely Pathogenic | 0.900 | 4847 | This work |
| chr19:g.39071043G>A | c.14545G>A | p.(Val4849Ile) | 0.817 | PS3_Mod, PS4, PM1_Sup, PP1_St | Pathogenic | 0.997 | 4849 | PMID: 33767344 |
| chr19:g.39071056C>T | c.14558C>T | p.(Thr4853Ile) | 0.982 | PM1_Sup, PP3_Mod | VUS | 0.500 | 4853 | This work |
| chr19:g.39071065C>G | c.14567C>G | p.(Ala4856Gly) | 0.937 | PM1_Sup, PP3_Mod | VUS | 0.500 | 4856 | This work |
| chr19:g.39071079C>T | c.14581C>T | p.(Arg4861Cys) | 0.916 | PS4_Sup, PM1_Sup, PM5_Sup, PP3_Mod | VUS | 0.812 | 4861 | This work |
| chr19:g.39071080G>A | c.14582G>A | p.(Arg4861His) | 0.912 | PS3_Mod, PS4_Sup, PM1_Sup, PP3_Mod | Likely Pathogenic | 0.900 | 4861 | PMID: 33767344 |
| chr19:g.39071125A>G | c.14627A>G | p.(Lys4876Arg) | 0.912 | PS4_Mod, PM1_Sup, PP1_Mod, PP3_Mod | Likely Pathogenic | 0.949 | 4876 | This work |
| chr19:g.39071137T>C | c.14639T>C | p.(Met4880Thr) | 0.956 | PS4_Sup, PM1_Sup, PP3_Mod | VUS | 0.675 | 4880 | This work |
| chr19:g.39075614G>A | c.14678G>A | p.(Arg4893Gln) | 0.963 | PM1_Sup, PP3_Mod | VUS | 0.500 | 4893 | This work |
| chr19:g.39075616G>A | c.14680G>A | p.(Ala4894Thr) | 0.922 | PS3_Mod, PM1_Sup, PP3_Mod | VUS | 0.812 | 4894 | This work |
| chr19:g.39075718A>G | c.14782A>G | p.(Ile4928Val) | 0.837 | PM1_Sup | VUS | 0.188 | 4928 | This work |
| chr19:g.39075739G>A | c.14803G>A | p.(Gly4935Ser) | 0.962 | PS4_Mod, PM1_Sup, PP1_St, PP3_Mod | Likely Pathogenic | 0.988 | 4935 | This work |
| chr19:g.39076587T>C | c.14813T>C | p.(Ile4938Thr) | 0.956 | PS4_Sup, PM1_Sup, PP1_St, PP3_Mod, BS2_Mod | Likely Pathogenic | 0.900 | 4938 | This work |
| chr19:g.39076588C>G | c.14814C>G | p.(Ile4938Met) | 0.855 | PM1_Sup, PP3_Mod | VUS | 0.500 | 4938 | This work |
| chr19:g.39076591C>A | c.14817C>A | p.(Asp4939Glu) | 0.808 | PS4_Sup, PM1_Sup | VUS | 0.325 | 4939 | This work |
| chr19:g.39076592G>A | c.14818G>A | p.(Ala4940Thr) | 0.882 | PS4_Mod, PM1_Sup, PP3_Mod | VUS | 0.812 | 4940 | This work |
| chr19:g.39076599G>T | c.14825G>T | p.(Gly4942Val) | 0.922 | PM1_Sup, PP3_Mod | VUS | 0.500 | 4942 | This work |
| chr19:g.39076741T>A | c.14879T>A | p.(Phe4960Tyr) | 0.965 | PM1_Sup, PP3_Mod | VUS | 0.500 | 4960 | This work |
| chr19:g.39076780C>T | c.14918C>T | p.(Pro4973Leu) | 0.898 | PS4_Mod, PM1_Sup, PP3_Mod | VUS | 0.812 | 4973 | PMID: 33767344 |
| chr19:g.39076830A>G | c.14968A>G | p.(Met4990Val) | 0.646 | PS4_Sup, PM1_Sup | VUS | 0.325 | 4990 | This work |
| chr19:g.39078002G>C | c.15059G>C | p.(Trp5020Ser) | 0.918 | PP3_Mod | VUS | 0.325 | 5020 | This work |
| chr19:g.39078003G>C | c.15060G>C | p.(Trp5020Cys) | 0.892 | PS4_Sup, PP3_Mod | VUS | 0.500 | 5020 | This work |

Supplemental Table 3

| | Benign (13) | Likely Benign (17) | VUS (219) | Likely Pathogenic (57) | Pathogenic (29) |
|---|---|---|---|---|---|
| **PS1** | 0 | 0 | 0 | 0 | 2 |
| **PS1_Moderate** | 0 | 0 | 0 | 1 | 0 |
| **PS2_PM6_Mod** | 0 | 0 | 1 | 2 | 1 |
| **PS2_PM6_Supp** | 0 | 0 | 1 | 2 | 0 |
| **PS3** | 0 | 0 | 0 | 0 | 4 |
| **PS3_Supp** | 0 | 0 | 0 | 1 | 0 |
| **PS3_Mod** | 0 | 1 | 3 | 18 | 23 |
| **PS4** | 0 | 0 | 2 | 4 | 19 |
| **PS4_Mod** | 0 | 0 | 19 | 42 | 8 |
| **PS4_Supp** | 0 | 3 | 89 | 9 | 2 |
| **PM1** | 1 | 3 | 70 | 28 | 17 |
| **PM1_Supp** | 0 | 1 | 42 | 18 | 9 |
| **PM5** | 0 | 0 | 3 | 7 | 5 |
| **PM5_Supp** | 0 | 0 | 6 | 4 | 1 |
| **PP1_Strong** | 0 | 0 | 1 | 10 | 21 |
| **PP1_Mod** | 0 | 0 | 2 | 7 | 3 |
| **PP1** | 0 | 1 | 7 | 13 | 5 |
| **PP3** | 0 | 3 | 91 | 46 | 27 |
| **BA1** | 11 | 0 | 0 | 0 | 0 |
| **BS1 Criteria** | 2 | 8 | 0 | 0 | 0 |
| **BS2** | 1 | 2 | 0 | 0 | 3 |
| **BS2_Moderate** | 1 | 9 | 2 | 4 | 6 |
| **BP2** | 1 | 0 | 1 | 0 | 0 |
| **BS3_Sup** | 1 | 3 | 4 | 0 | 0 |
| **BP4** | 1 | 2 | 18 | 0 | 0 |

Supplemental Table 4

| Genomic Position | posterior probability | gnomAD Allele Frequencies | | | | | | Allele Contribution (Allele frequency * Posterior probability) | | | | | |
|---|---|---|---|---|---|---|---|---|---|---|---|---|---|
| | | vep_gnomad_af | ep_gnomad_af_a | ep_gnomad_af_s | ep_gnomad_af_a | ep_gnomad_af_e | ep_gnomad_af_r | vep_gnomad_af | vep_gnomad_af_afr | vep_gnomad_af_sas | vep_gnomad_af_am | vep_gnomad_af_eas | vep_gnomad_af_nfe |
| chr19:g.38924507T>G | 0.94924 | | | | | | | 0 | 0 | 0 | 0 | 0 | 0 |
| chr19:g.38931390_38931392del | 0.32459 | | | | | | | 0 | 0 | 0 | 0 | 0 | 0 |
| chr19:g.38931436A>G | 0.89991 | | | | | | | 0 | 0 | 0 | 0 | 0 | 0 |
| chr19:g.38931442T>C | 0.99409 | 4.21E-06 | 0 | 0 | 0 | 0 | 9.47E-06 | 4.19002E-06 | 0 | 0 | 0 | 0 | 9.40963E-06 |
| chr19:g.38931458G>C | 0.81214 | | | | | | | 0 | 0 | 0 | 0 | 0 | 0 |
| chr19:g.38931469C>T | 0.94793 | 4.13E-06 | 0 | 0 | 2.94E-05 | 0 | 0 | 3.91966E-06 | 0 | 0 | 2.79E-05 | 0 | 0 |
| chr19:g.38931470G>A | 0.81214 | 1.24E-05 | 6.63E-05 | 0 | 0 | 0 | 1.85E-05 | 1.00672E-05 | 5.38127E-05 | 0 | 0 | 0 | 1.49844E-05 |
| chr19:g.38931491C>A | 0.32459 | 5.46E-05 | 0 | 0 | 0 | 0 | 0.000120602 | 1.77123E-05 | 0 | 0 | 0 | 0 | 3.91462E-05 |
| chr19:g.38933001G>A | 0.49988 | 7.09E-06 | 0 | 0 | 0 | 0 | 1.56E-05 | 3.5456E-06 | 0 | 0 | 0 | 0 | 7.77709E-06 |
| chr19:g.38933001G>T | 0.81214 | | | | | | | 0 | 0 | 0 | 0 | 0 | 0 |
| chr19:g.38933013T>C | 0.32459 | | | | | | | 0 | 0 | 0 | 0 | 0 | 0 |
| chr19:g.38933035C>A | 0.67519 | | | | | | | 0 | 0 | 0 | 0 | 0 | 0 |
| chr19:g.38933074C>T | 0.67519 | 3.19E-05 | 0 | 3.27E-05 | 0 | 0.000300782 | 1.55E-05 | 2.15286E-05 | 0 | 2.20535E-05 | 0 | 0.000203085 | 1.04863E-05 |
| chr19:g.38934430G>A | 0.05072 | 0.000134506 | 0.000120356 | 9.80E-05 | 0.000141155 | 0 | 0.000201679 | 6.82213E-06 | 6.10447E-06 | 4.96995E-06 | 7.15939E-06 | 0 | 1.02291E-05 |
| chr19:g.38934819C>A | 0.81214 | | | | | | | 0 | 0 | 0 | 0 | 0 | 0 |
| chr19:g.38934827C>A | 0.89991 | | | | | | | 0 | 0 | 0 | 0 | 0 | 0 |
| chr19:g.38934831G>A | 0.67519 | | | | | | | 0 | 0 | 0 | 0 | 0 | 0 |
| chr19:g.38934843A>G | 0.81214 | | | | | | | 0 | 0 | 0 | 0 | 0 | 0 |
| chr19:g.38934851C>T | 0.99968 | | | | | | | 0 | 0 | 0 | 0 | 0 | 0 |
| chr19:g.38934852G>A | 0.81214 | 7.96E-06 | 0 | 3.27E-05 | 2.89E-05 | 0 | 0 | 6.46059E-06 | 0 | 2.65284E-05 | 2.34927E-05 | 0 | 0 |
| chr19:g.38934852G>T | 0.94793 | | | | | | | 0 | 0 | 0 | 0 | 0 | 0 |
| chr19:g.38934857G>A | 0.81214 | | | | | | | 0 | 0 | 0 | 0 | 0 | 0 |
| chr19:g.38934860G>A | 0.67519 | | | | | | | 0 | 0 | 0 | 0 | 0 | 0 |
| chr19:g.38934861A>G | 0.67519 | | | | | | | 0 | 0 | 0 | 0 | 0 | 0 |
| chr19:g.38934890G>A | 0.67519 | 3.19E-05 | 0 | 6.54E-05 | 2.90E-05 | 0 | 4.40E-05 | 2.15147E-05 | 0 | 4.41705E-05 | 1.95526E-05 | 0 | 2.97357E-05 |
| chr19:g.38934892G>T | 0.67519 | | | | | | | 0 | 0 | 0 | 0 | 0 | 0 |
| chr19:g.38934893C>T | 0.99863 | 3.98E-06 | 0 | 0 | 0 | 0 | 0 | 3.97924E-06 | 0 | 0 | 0 | 0 | 0 |
| chr19:g.38934897A>C | 0.81214 | | | | | | | 0 | 0 | 0 | 0 | 0 | 0 |
| chr19:g.38934897A>G | 0.94793 | | | | | | | 0 | 0 | 0 | 0 | 0 | 0 |
| chr19:g.38935311G>A | 0.18771 | 2.01E-05 | 6.23E-05 | 0 | 2.91E-05 | 5.46E-05 | 1.78E-05 | 3.76931E-06 | 1.1688E-05 | 0 | 5.45764E-06 | 1.02574E-05 | 3.33748E-06 |
| chr19:g.38937121C>T | 0.32459 | 9.55E-05 | 4.01E-05 | 6.53E-05 | 0 | 0.000100231 | 0.000170358 | 3.09888E-05 | 1.30023E-05 | 2.12039E-05 | 0 | 3.25338E-05 | 5.52964E-05 |
| chr19:g.38937132G>A | 0.49988 | 1.06E-05 | 0 | 0 | 0 | 0 | 2.32E-05 | 5.30189E-06 | 0 | 0 | 0 | 0 | 1.16111E-05 |
| chr19:g.38937157T>A | 0.67519 | | | | | | | 0 | 0 | 0 | 0 | 0 | 0 |
| chr19:g.38937160A>T | 0.81214 | | | | | | | 0 | 0 | 0 | 0 | 0 | 0 |
| chr19:g.38937350G>A | 0.98779 | | | | | | | 0 | 0 | 0 | 0 | 0 | 0 |
| chr19:g.38937350G>C | 0.99863 | 1.42E-05 | 0 | 3.27E-05 | 0 | 5.01E-05 | 1.55E-05 | 1.41363E-05 | 0 | 3.26179E-05 | 0 | 5.00516E-05 | 1.54862E-05 |
| chr19:g.38939140C>T | 0.32459 | 1.99E-05 | 0 | 3.27E-05 | 0 | 0 | 3.52E-05 | 6.46094E-06 | 0 | 1.0602E-05 | 0 | 0 | 1.14302E-05 |
| chr19:g.38939141G>T | 0.67519 | 1.06E-05 | 0 | 0 | 0 | 0.000100241 | 7.75E-06 | 7.17041E-06 | 0 | 0 | 0 | 6.76814E-05 | 5.23582E-06 |
| chr19:g.38939313C>T | 0.89991 | 1.99E-05 | 6.16E-05 | 6.53E-05 | 0 | 0 | 8.79E-06 | 1.7896E-05 | 5.53995E-05 | 5.87869E-05 | 0 | 0 | 7.91324E-06 |
| chr19:38939320_38939321insGGA | 0.49988 | | | | | | | 0 | 0 | 0 | 0 | 0 | 0 |
| chr19:g.38939352G>A | 0.99863 | | | | | | | 0 | 0 | 0 | 0 | 0 | 0 |
| chr19:g.38939352G>C | 0.99863 | | | | | | | 0 | 0 | 0 | 0 | 0 | 0 |
| chr19:g.38939355G>A | 0.81214 | | | | | | | 0 | 0 | 0 | 0 | 0 | 0 |
| chr19:g.38939431G>A | 0.67519 | 7.96E-06 | 0 | 0 | 0 | 0 | 8.81E-06 | 5.37756E-06 | 0 | 0 | 0 | 0 | 5.94734E-06 |
| chr19:g.38939431G>T | 0.81214 | | | | | | | 0 | 0 | 0 | 0 | 0 | 0 |
| chr19:g.38942425C>A | 0.49988 | | | | | | | 0 | 0 | 0 | 0 | 0 | 0 |
| chr19:g.38942482C>A | 0.67519 | | | | | | | 0 | 0 | 0 | 0 | 0 | 0 |
| chr19:g.38942482C>G | 0.81200 | 3.98E-06 | 0 | 0 | 0 | 0 | 8.79E-06 | 3.22973E-06 | 0 | 0 | 0 | 0 | 7.14135E-06 |
| chr19:g.38942482C>T | 0.99409 | 7.07E-06 | 0 | 0 | 0 | 5.01E-05 | 0 | 7.03168E-06 | 0 | 0 | 0 | 4.98291E-05 | 0 |
| chr19:g.38942483G>A | 0.99715 | 3.98E-06 | 0 | 0 | 0 | 0 | 0 | 3.96623E-06 | 0 | 0 | 0 | 0 | 0 |
| chr19:g.38942483G>T | 0.94793 | | | | | | | 0 | 0 | 0 | 0 | 0 | 0 |
| chr19:g.38943625C>T | 0.32459 | 4.03E-06 | 0 | 0 | 0 | 0 | 8.96E-06 | 1.30783E-06 | 0 | 0 | 0 | 0 | 2.90757E-06 |
| chr19:g.38943636G>T | 0.32459 | | | | | | | 0 | 0 | 0 | 0 | 0 | 0 |
| chr19:g.38945887A>G | 0.32459 | 0.000335874 | 0.000160295 | 0 | 0.000225759 | 5.01E-05 | 0.000611588 | 0.000109021 | 5.20301E-05 | 0 | 7.32792E-05 | 1.62702E-05 | 0.000198515 |
| chr19:g.38945893C>G | 0.49988 | | | | | | | 0 | 0 | 0 | 0 | 0 | 0 |
| chr19:g.38945894T>C | 0.81214 | | | | | | | 0 | 0 | 0 | 0 | 0 | 0 |
| chr19:g.38945909G>A | 0.32459 | 1.41E-05 | 8.02E-05 | 0 | 0 | 0 | 1.55E-05 | 4.59012E-06 | 2.60192E-05 | 0 | 0 | 0 | 5.02524E-06 |
| chr19:g.38945987T>C | 0.32459 | 3.19E-05 | 0 | 0 | 0 | 0 | 6.49E-05 | 1.03551E-05 | 0 | 0 | 0 | 0 | 2.10745E-05 |
| chr19:g.38945999A>C | 0.99409 | | | | | | | 0 | 0 | 0 | 0 | 0 | 0 |
| chr19:g.38945999A>G | 0.94924 | | | | | | | 0 | 0 | 0 | 0 | 0 | 0 |
| chr19:g.38946103G>A | 0.81200 | 5.57E-05 | 6.15E-05 | 3.27E-05 | 0.00014455 | 0 | 5.27E-05 | 4.52019E-05 | 4.99508E-05 | 2.65221E-05 | 0.000117375 | 0 | 4.28232E-05 |
| chr19:g.38946111C>A | 0.67519 | | | | | | | 0 | 0 | 0 | 0 | 0 | 0 |
| chr19:g.38946111C>T | 0.99715 | | | | | | | 0 | 0 | 0 | 0 | 0 | 0 |
| chr19:g.38946112G>A | 0.67519 | 0.000113131 | 0.000440811 | 0 | 0.000141091 | 0 | 0.00012386 | 7.63849E-05 | 0.000297631 | 0 | 9.52636E-05 | 0 | 8.36291E-05 |
| chr19:g.38946129T>C | 0.94924 | | | | | | | 0 | 0 | 0 | 0 | 0 | 0 |
| chr19:g.38946129T>G | 0.89991 | | | | | | | 0 | 0 | 0 | 0 | 0 | 0 |
| chr19:g.38946144G>T | 0.89991 | | | | | | | 0 | 0 | 0 | 0 | 0 | 0 |
| chr19:g.38946168C>T | 0.94924 | 3.98E-06 | 0 | 0 | 2.89E-05 | 0 | 0 | 3.77482E-06 | 0 | 0 | 2.7441E-05 | 0 | 0 |
| chr19:g.38948179G>C | 0.67519 | | | | | | | 0 | 0 | 0 | 0 | 0 | 0 |
| chr19:g.38948185C>T | 0.99863 | 0.000106059 | 0 | 0 | 0 | 5.01E-05 | 0.000193519 | 0.000105913 | 0 | 0 | 0 | 5.00516E-05 | 0.000193254 |
| chr19:g.38948186G>T | 0.99863 | | | | | | | 0 | 0 | 0 | 0 | 0 | 0 |
| chr19:g.38948815G>C | 0.49988 | | | | | | | 0 | 0 | 0 | 0 | 0 | 0 |
| chr19:g.38948887G>A | 0.10000 | 0.000543568 | 0.000123082 | 0 | 5.65E-05 | 0 | 0.000748432 | 5.43568E-05 | 1.23082E-05 | 0 | 5.6494E-06 | 0 | 7.48432E-05 |
| chr19:g.38951101C>T | 0.02505 | 2.12E-05 | 0 | 0 | 0 | 0 | 4.64E-05 | 5.31317E-07 | 0 | 0 | 0 | 0 | 1.16344E-06 |
| chr19:g.38951191C>T | 0.18771 | | | | | | | 0 | 0 | 0 | 0 | 0 | 0 |
| chr19:g.38954139G>A | 0.10000 | 0.000163795 | 8.06E-05 | 0.00042492 | 0 | 0.000100583 | 0.000225512 | 1.63795E-05 | 8.06127E-06 | 4.2492E-05 | 0 | 1.00583E-05 | 2.25512E-05 |
| chr19:g.38956784G>A | 0.10000 | 1.22E-05 | 0 | 0 | 0 | 0 | 1.80E-05 | 1.2167E-06 | 0 | 0 | 0 | 0 | 1.80261E-06 |
| chr19:g.38956856G>A | 0.10000 | 6.82E-05 | 0 | 0 | 8.70E-05 | 0 | 0.00011516 | 6.81931E-06 | 0 | 0 | 8.69817E-06 | 0 | 1.1516E-05 |
| chr19:g.38956955G>A | 0.10000 | 0.000101349 | 0 | 6.77E-05 | 0 | 0 | 0.000207177 | 1.01349E-05 | 0 | 6.77461E-06 | 0 | 0 | 2.07177E-05 |
| chr19:g.38956987C>T | 0.32459 | 2.95E-05 | 0 | 0 | 1.00E-04 | 0 | 3.38E-05 | 9.56937E-06 | 0 | 0 | 3.24525E-05 | 0 | 1.09567E-05 |
| chr19:g.38957026G>A | 0.18771 | | | | | | | 0 | 0 | 0 | 0 | 0 | 0 |
| chr19:g.38957026G>C | 0.89991 | | | | | | | 0 | 0 | 0 | 0 | 0 | 0 |
| chr19:g.38957032G>A | 0.32459 | | | | | | | 0 | 0 | 0 | 0 | 0 | 0 |
| chr19:g.38958295G>A | 0.05072 | 3.98E-06 | 0 | 0 | 2.89E-05 | 0 | 0 | 2.01686E-07 | 0 | 0 | 1.46623E-06 | 0 | 0 |
| chr19:g.38959642C>T | 0.10000 | 2.39E-05 | 0 | 3.27E-05 | 0.00011566 | 0 | 8.79E-06 | 2.38618E-06 | 0 | 3.26627E-06 | 1.1566E-05 | 0 | 8.79183E-07 |
| chr19:g.38959751C>T | 0.10000 | 7.56E-05 | 0 | 0.000588005 | 0 | 0 | 0 | 7.55533E-06 | 0 | 5.88005E-05 | 0 | 0 | 0 |
| chr19:g.38960044A>C | 0.32459 | | | | | | | 0 | 0 | 0 | 0 | 0 | 0 |
| chr19:g.38960055G>A | 0.10000 | 1.19E-05 | 0 | 0 | 0 | 0 | 2.64E-05 | 1.19334E-06 | 0 | 0 | 0 | 0 | 2.63889E-06 |
| chr19:g.38968456A>G | 0.10000 | | | | | | | 0 | 0 | 0 | 0 | 0 | 0 |
| chr19:g.38973933A>G | 0.00590 | 0.000854787 | 0.000215936 | 0.000667891 | 0.001027241 | 0 | 0.001371346 | 5.04324E-06 | 1.27402E-06 | 3.94056E-06 | 6.06072E-06 | 0 | 8.09094E-06 |
| chr19:g.38973969C>T | 0.10000 | 0.000197171 | 0 | 0 | 0 | 0 | 0.000121563 | 1.97171E-05 | 0 | 0 | 0 | 0 | 1.21563E-05 |
| chr19:g.38973985C>T | 0.10000 | 6.51E-05 | 0.00016276 | 0 | 0 | 0.000141523 | 9.59E-05 | 6.51348E-06 | 1.6276E-05 | 0 | 0 | 1.41523E-05 | 9.5944E-06 |
| chr19:g.38973997C>T | 0.32459 | 5.90E-06 | 0 | 0 | 0 | 0 | 1.45E-05 | 1.91575E-06 | 0 | 0 | 0 | 0 | 4.72007E-06 |
| chr19:g.38976319T>C | 0.32459 | | | | | | | 0 | 0 | 0 | 0 | 0 | 0 |

| | | | | | | | | | | | | | |
|---|---|---|---|---|---|---|---|---|---|---|---|---|---|
| chr19:g.38976328A>G | 0.10000 | 1.80E-05 | 0 | 3.27E-05 | 2.83E-05 | 0 | 2.36E-05 | 1.80261E-06 | 0 | 3.27397E-06 | 2.83142E-06 | 0 | 2.36466E-06 |
| chr19:g.38976331G>A | 0.00590 | 0.001198201 | 4.05E-05 | 0 | 0.000141611 | 0 | 0.00222559 | 7.06939E-06 | 2.39157E-07 | 0 | 8.35505E-07 | 0 | 1.3131E-05 |
| chr19:g.38976427A>G | 0.32459 | 1.22E-05 | 0.000124797 | 0 | 0 | 0 | 9.09E-06 | 3.95013E-06 | 4.05079E-05 | 0 | 0 | 0 | 2.94953E-06 |
| chr19:g.38976478C>T | 0.49988 | 4.06E-06 | 0 | 0 | 2.90E-05 | 0 | 0 | 2.0317E-06 | 0 | 0 | 1.44893E-05 | 0 | 0 |
| chr19:g.38976481T>G | 0.18771 | | | | | | | 0 | 0 | 0 | 0 | 0 | 0 |
| chr19:g.38976636T>C | 0.10000 | 1.23E-05 | 0 | 0 | 0 | 0 | 1.85E-05 | 1.2258E-06 | 0 | 0 | 0 | 0 | 1.84597E-06 |
| chr19:g.38976735A>G | 0.10000 | | | | | | | 0 | 0 | 0 | 0 | 0 | 0 |
| chr19:g.38976736T>A | 0.10000 | | | | | | | 0 | 0 | 0 | 0 | 0 | 0 |
| chr19:g.38980791C>T | 0.10000 | 0 | 0 | 0 | 0 | 0 | 0 | 0 | 0 | 0 | 0 | 0 | 0 |
| chr19:g.38981282A>C | 0.10000 | | | | | | | 0 | 0 | 0 | 0 | 0 | 0 |
| chr19:g.38985019T>A | 0.18771 | 0.000218677 | 0 | 0 | 5.65E-05 | 0 | 8.73E-05 | 4.10479E-05 | 0 | 0 | 1.06051E-05 | 0 | 1.63785E-05 |
| chr19:g.38985021G>C | 0.32459 | 8.07E-06 | 0 | 0 | 0 | 0 | 1.81E-05 | 2.62092E-06 | 0 | 0 | 0 | 0 | 5.86209E-06 |
| chr19:g.38985066G>C | 0.89991 | 4.00E-06 | 0 | 0 | 0 | 0 | 8.88E-06 | 3.60056E-06 | 0 | 0 | 0 | 0 | 7.99295E-06 |
| chr19:g.38985094G>A | 0.49988 | 7.99E-06 | 0 | 0 | 0 | 0 | 1.77E-05 | 3.99297E-06 | 0 | 0 | 0 | 0 | 8.83867E-06 |
| chr19:g.38985104C>G | 0.89991 | | | | | | | 0 | 0 | 0 | 0 | 0 | 0 |
| chr19:g.38985105G>A | 0.67519 | 1.20E-05 | 0 | 0 | 0 | 5.46E-05 | 1.77E-05 | 8.09891E-06 | 0 | 0 | 0 | 3.68352E-05 | 1.19589E-05 |
| chr19:g.38985195G>A | 0.32459 | 8.51E-05 | 0.000641591 | 0.000163345 | 2.82E-05 | 0 | 7.79E-06 | 2.76362E-05 | 0.000208254 | 5.30203E-05 | 9.16041E-06 | 0 | 2.52867E-06 |
| chr19:g.38985204C>T | 0.99715 | 1.20E-05 | 0 | 3.27E-05 | 0 | 0 | 8.86E-06 | 1.19436E-05 | 0 | 3.25738E-05 | 0 | 0 | 8.83481E-06 |
| chr19:g.38985205G>A | 0.99409 | 0 | 0 | 0 | 0 | 0 | 0 | 0 | 0 | 0 | 0 | 0 | 0 |
| chr19:g.38985205G>C | 0.94793 | | | | | | | 0 | 0 | 0 | 0 | 0 | 0 |
| chr19:g.38985205G>T | 0.81214 | | | | | | | 0 | 0 | 0 | 0 | 0 | 0 |
| chr19:g.38985219G>A | 0.99968 | | | | | | | 0 | 0 | 0 | 0 | 0 | 0 |
| chr19:g.38985261A>T | 0.49988 | | | | | | | 0 | 0 | 0 | 0 | 0 | 0 |
| chr19:g.38985265G>A | 0.49988 | | | | | | | 0 | 0 | 0 | 0 | 0 | 0 |
| chr19:g.38986905C>T | 0.81214 | 3.89E-05 | 0 | 3.27E-05 | 5.64E-05 | 0.000150391 | 3.87E-05 | 3.16109E-05 | 0 | 2.65267E-05 | 4.58371E-05 | 0.000122139 | 3.14481E-05 |
| chr19:g.38986918C>G | 0.89991 | | | | | | | 0 | 0 | 0 | 0 | 0 | 0 |
| chr19:g.38986923C>G | 0.99409 | | | | | | | 0 | 0 | 0 | 0 | 0 | 0 |
| chr19:g.38986923C>T | 0.99715 | 2.12E-05 | 0 | 0 | 0 | 5.01E-05 | 3.87E-05 | 2.11699E-05 | 0 | 0 | 0 | 4.99925E-05 | 3.86163E-05 |
| chr19:g.38986934G>T | 0.89991 | | | | | | | 0 | 0 | 0 | 0 | 0 | 0 |
| chr19:g.38986941T>A | 0.67519 | | | | | | | 0 | 0 | 0 | 0 | 0 | 0 |
| chr19:g.38986946G>A | 0.32459 | 1.42E-05 | 0.000160282 | 0 | 0 | 0 | 0 | 4.59583E-06 | 5.2026E-05 | 0 | 0 | 0 | 0 |
| chr19:g.38987055C>T | 0.02505 | 0.00014522 | 4.01E-05 | 0.000914554 | 5.65E-05 | 0 | 7.77E-05 | 3.63776E-06 | 1.00457E-06 | 2.29096E-05 | 1.41453E-06 | 0 | 1.94554E-06 |
| chr19:g.38987056G>A | 0.32459 | 2.13E-05 | 0 | 3.27E-05 | 2.82E-05 | 0 | 2.33E-05 | 6.89991E-06 | 0 | 1.06027E-05 | 9.16351E-06 | 0 | 7.5655E-06 |
| chr19:g.38987095G>A | 0.81214 | | | | | | | 0 | 0 | 0 | 0 | 0 | 0 |
| chr19:g.38987127C>T | 0.32459 | 9.16E-05 | 0 | 0 | 0.000231294 | 0.000108826 | 0.000105768 | 2.9731E-05 | 0 | 0 | 7.50757E-05 | 3.53238E-05 | 3.43312E-05 |
| chr19:g.38987128G>A | 0.32459 | 3.98E-06 | 0 | 0 | 0 | 0 | 8.82E-06 | 1.29276E-06 | 0 | 0 | 0 | 0 | 2.86134E-06 |
| chr19:g.38987142C>T | 0.81214 | | | | | | | 0 | 0 | 0 | 0 | 0 | 0 |
| chr19:g.38987541G>A | 0.67519 | 2.40E-05 | 0 | 0 | 2.90E-05 | 0 | 3.54E-05 | 1.62043E-05 | 0 | 0 | 1.95481E-05 | 0 | 2.39353E-05 |
| chr19:g.38987550A>C | 0.81214 | | | | | | | 0 | 0 | 0 | 0 | 0 | 0 |
| chr19:g.38989817A>G | 0.00000 | 0.000456172 | 0.000440635 | 0 | 0.000254008 | 0 | 0.00081326 | 0 | 0 | 0 | 0 | 0 | 0 |
| chr19:g.38989863G>A | 0.99934 | | | | | | | 0 | 0 | 0 | 0 | 0 | 0 |
| chr19:g.38989874T>C | 0.49988 | | | | | | | 0 | 0 | 0 | 0 | 0 | 0 |
| chr19:g.38989881A>G | 0.01220 | 0.001018582 | 0.000160256 | 0.002057748 | 0.000338677 | 5.01E-05 | 0.001464254 | 1.24267E-05 | 1.95513E-06 | 2.51045E-05 | 4.13186E-06 | 6.11468E-07 | 1.78639E-05 |
| chr19:g.38990279G>C | 0.32459 | | | | | | | 0 | 0 | 0 | 0 | 0 | 0 |
| chr19:g.38990282C>A | 0.89991 | | | | | | | 0 | 0 | 0 | 0 | 0 | 0 |
| chr19:g.38990283G>A | 0.89991 | 8.39E-06 | 0 | 3.27E-05 | 0 | 0 | 8.99E-06 | 7.54736E-06 | 0 | 2.94454E-05 | 0 | 0 | 8.08995E-06 |
| chr19:g.38990289_38990291del | 0.94924 | | | | | | | 0 | 0 | 0 | 0 | 0 | 0 |
| chr19:g.38990290A>G | 0.89991 | 3.19E-05 | 0.000115048 | 0 | 0 | 0 | 0 | 2.87199E-05 | 0.000103533 | 0 | 0 | 0 | 0 |
| chr19:g.38990295G>A | 0.99968 | 0 | 0 | 0 | 0 | 0 | 0 | 0 | 0 | 0 | 0 | 0 | 0 |
| chr19:g.38990307G>A | 0.94793 | | | | | | | 0 | 0 | 0 | 0 | 0 | 0 |
| chr19:g.38990310C>T | 0.99968 | 2.09E-05 | 0 | 0 | 0 | 0 | 3.58E-05 | 2.08792E-05 | 0 | 0 | 0 | 0 | 3.58039E-05 |
| chr19:g.38990320T>A | 0.67519 | | | | | | | 0 | 0 | 0 | 0 | 0 | 0 |
| chr19:g.38990322C>T | 0.67519 | 8.35E-06 | 0 | 0 | 2.90E-05 | 0 | 8.97E-06 | 5.6397E-06 | 0 | 0 | 1.9606E-05 | 0 | 6.05356E-06 |
| chr19:g.38990323G>A | 0.89991 | 8.36E-06 | 0 | 0 | 0 | 0 | 8.97E-06 | 7.52276E-06 | 0 | 0 | 0 | 0 | 8.07587E-06 |
| chr19:g.38990331G>A | 0.89991 | | | | | | | 0 | 0 | 0 | 0 | 0 | 0 |
| chr19:g.38990332A>G | 0.81214 | | | | | | | 0 | 0 | 0 | 0 | 0 | 0 |
| chr19:g.38990336C>G | 0.67519 | | | | | | | 0 | 0 | 0 | 0 | 0 | 0 |
| chr19:g.38990337T>G | 0.89991 | | | | | | | 0 | 0 | 0 | 0 | 0 | 0 |
| chr19:g.38990344C>G | 0.81214 | | | | | | | 0 | 0 | 0 | 0 | 0 | 0 |
| chr19:g.38990346G>A | 0.81214 | 4.82E-05 | 0 | 0 | 0.000142304 | 0 | 5.56E-05 | 3.91562E-05 | 0 | 0 | 0.000115571 | 0 | 4.51225E-05 |
| chr19:g.38990359A>G | 0.81214 | | | | | | | 0 | 0 | 0 | 0 | 0 | 0 |
| chr19:g.38990370G>A | 0.89991 | | | | | | | 0 | 0 | 0 | 0 | 0 | 0 |
| chr19:g.38990371G>C | 0.99409 | | | | | | | 0 | 0 | 0 | 0 | 0 | 0 |
| chr19:g.38990446A>G | 0.32459 | 1.50E-05 | 8.49E-05 | 0 | 0 | 0 | 1.64E-05 | 4.85579E-06 | 2.7552E-05 | 0 | 0 | 0 | 5.31079E-06 |
| chr19:g.38990457G>A | 0.49988 | 3.35E-05 | 0.000127119 | 3.31E-05 | 2.86E-05 | 5.12E-05 | 1.64E-05 | 1.67609E-05 | 6.35441E-05 | 1.65315E-05 | 1.43191E-05 | 2.55798E-05 | 8.18912E-06 |
| chr19:g.38990615G>A | 0.94793 | 1.41E-05 | 0 | 3.27E-05 | 0 | 0 | 1.55E-05 | 1.34107E-05 | 0 | 3.09619E-05 | 0 | 0 | 1.46866E-05 |
| chr19:g.38990624G>A | 0.89991 | 7.96E-06 | 6.16E-05 | 0 | 0 | 5.44E-05 | 0 | 7.16232E-06 | 5.53927E-05 | 0 | 0 | 4.89294E-05 | 0 |
| chr19:g.38990624G>T | 0.98779 | | | | | | | 0 | 0 | 0 | 0 | 0 | 0 |
| chr19:g.38990625A>T | 0.81214 | | | | | | | 0 | 0 | 0 | 0 | 0 | 0 |
| chr19:g.38990633G>A | 0.99863 | 3.89E-05 | 4.01E-05 | 0 | 0 | 0 | 7.75E-05 | 3.88643E-05 | 4.00156E-05 | 0 | 0 | 0 | 7.74012E-05 |
| chr19:g.38990637G>A | 0.99968 | | | | | | | 0 | 0 | 0 | 0 | 0 | 0 |
| chr19:g.38990637G>T | 0.94924 | | | | | | | 0 | 0 | 0 | 0 | 0 | 0 |
| chr19:g.38990640G>A | 0.81214 | | | | | | | 0 | 0 | 0 | 0 | 0 | 0 |
| chr19:g.38990643C>T | 0.89991 | | | | | | | 0 | 0 | 0 | 0 | 0 | 0 |
| chr19:g.38990650G>C | 0.32459 | | | | | | | 0 | 0 | 0 | 0 | 0 | 0 |
| chr19:g.38991276C>T | 0.94793 | | | | | | | 0 | 0 | 0 | 0 | 0 | 0 |
| chr19:g.38991277G>A | 0.67519 | 3.98E-06 | 0 | 0 | 0 | 0 | 8.81E-06 | 2.68793E-06 | 0 | 0 | 0 | 0 | 5.94617E-06 |
| chr19:g.38991277G>C | 0.67519 | | | | | | | 0 | 0 | 0 | 0 | 0 | 0 |
| chr19:g.38991280T>C | 0.94924 | | | | | | | 0 | 0 | 0 | 0 | 0 | 0 |
| chr19:g.38991282C>T | 0.99409 | 1.59E-05 | 0 | 6.53E-05 | 0 | 0 | 8.81E-06 | 1.58295E-05 | 0 | 6.49435E-05 | 0 | 0 | 8.75434E-06 |
| chr19:g.38991283G>A | 0.99863 | 7.96E-06 | 6.16E-05 | 0 | 0 | 0 | 8.81E-06 | 7.95056E-06 | 6.15375E-05 | 0 | 0 | 0 | 8.79308E-06 |
| chr19:g.38991294C>T | 0.98779 | 7.08E-06 | 0 | 0 | 0 | 0 | 7.75E-06 | 6.99004E-06 | 0 | 0 | 0 | 0 | 7.65646E-06 |
| chr19:g.38991295G>A | 0.99968 | 7.96E-06 | 0 | 0 | 0 | 5.44E-05 | 8.80E-06 | 7.95803E-06 | 0 | 0 | 0 | 5.43836E-05 | 8.80108E-06 |
| chr19:g.38991295G>T | 0.94924 | | | | | | | 0 | 0 | 0 | 0 | 0 | 0 |
| chr19:g.38991307C>T | 0.32459 | 7.08E-06 | 0 | 3.27E-05 | 2.82E-05 | 0 | 0 | 2.2972E-06 | 0 | 1.06027E-05 | 9.16351E-06 | 0 | 0 |
| chr19:g.38991503C>T | 0.32459 | 8.84E-05 | 0 | 0 | 0 | 0.000651694 | 5.42E-05 | 2.87028E-05 | 0 | 0 | 0 | 0.000211533 | 1.76019E-05 |
| chr19:g.38991538C>T | 0.94924 | | | | | | | 0 | 0 | 0 | 0 | 0 | 0 |
| chr19:g.38991539G>A | 0.98779 | 3.98E-06 | 0 | 0 | 0 | 0 | 0 | 3.93038E-06 | 0 | 0 | 0 | 0 | 0 |
| chr19:g.38991544T>C | 0.32459 | | | | | | | 0 | 0 | 0 | 0 | 0 | 0 |
| chr19:g.38993292A>G | 0.32459 | 2.00E-05 | 0 | 0 | 0.000115788 | 0 | 0 | 6.48775E-06 | 0 | 0 | 3.75835E-05 | 0 | 0 |
| chr19:g.38993303C>G | 0.10000 | 1.07E-05 | 4.02E-05 | 0 | 2.82E-05 | 0 | 7.77E-06 | 1.06719E-06 | 4.02026E-06 | 0 | 2.82438E-06 | 0 | 7.77303E-07 |
| chr19:g.38993303C>T | 0.10000 | | | | | | | 0 | 0 | 0 | 0 | 0 | 0 |
| chr19:g.38993309C>G | 0.10000 | 3.56E-06 | 0 | 0 | 0 | 0 | 7.77E-06 | 3.56039E-07 | 0 | 0 | 0 | 0 | 7.77158E-07 |
| chr19:g.38993310G>A | 0.18771 | 2.00E-05 | 0 | 0 | 0 | 0 | 3.53E-05 | 3.76218E-06 | 0 | 0 | 0 | 0 | 6.62724E-06 |
| chr19:g.38993319C>T | 0.10000 | 2.41E-05 | 0 | 0 | 0.000144684 | 0 | 0 | 2.40668E-06 | 0 | 0 | 1.44684E-05 | 0 | 0 |
| chr19:g.38993348T>A | 0.02505 | 4.06E-06 | 0 | 0 | 0 | 0 | 8.90E-06 | 1.01788E-07 | 0 | 0 | 0 | 0 | 2.22928E-07 |
| chr19:g.38993563G>A | 0.32459 | | | | | | | 0 | 0 | 0 | 0 | 0 | 0 |

| | | | | | | | | | | | | | |
|---|---|---|---|---|---|---|---|---|---|---|---|---|---|
| chr19:g.38993563G>C | 0.94793 | 3.19E-05 | 0 | 0 | 0 | 0 | 6.50E-05 | 3.02466E-05 | 0 | 0 | 0 | 0 | 6.15699E-05 |
| chr19:g.38994938G>A | 0.10000 | | | | | | | 0 | 0 | 0 | 0 | 0 | 0 |
| chr19:g.38994959C>T | 0.89991 | 1.19E-05 | 0 | 0 | 0 | 0 | 2.64E-05 | 1.07402E-05 | 0 | 0 | 0 | 0 | 2.3751E-05 |
| chr19:g.38994987C>T | 0.18771 | 3.98E-06 | 0 | 3.27E-05 | 0 | 0 | 0 | 7.47426E-07 | 0 | 6.13191E-06 | 0 | 0 | 0 |
| chr19:g.38995508G>C | 0.32459 | | | | | | | 0 | 0 | 0 | 0 | 0 | 0 |
| chr19:g.38995509A>G | 0.81214 | | | | | | | 0 | 0 | 0 | 0 | 0 | 0 |
| chr19:g.38995518G>A | 0.67519 | | | | | | | 0 | 0 | 0 | 0 | 0 | 0 |
| chr19:g.38995701G>A | 0.10000 | 1.19E-05 | 0 | 0 | 0 | 0.000108731 | 8.79E-06 | 1.19294E-06 | 0 | 0 | 0 | 1.08731E-05 | 8.79028E-07 |
| chr19:g.38995965C>T | 0.00590 | 0.00072705 | 0.000204666 | 6.62E-05 | 0.00011352 | 0 | 0.001449569 | 4.28959E-06 | 1.20753E-06 | 3.90392E-07 | 6.6977E-07 | 0 | 8.55246E-06 |
| chr19:g.38996563C>T | 0.10000 | 1.78E-05 | 0 | 0.000130779 | 0 | 0 | 0 | 1.77699E-06 | 0 | 1.30779E-05 | 0 | 0 | 0 |
| chr19:g.38996572T>C | 0.18771 | 8.02E-06 | 0 | 3.27E-05 | 0 | 0 | 8.88E-06 | 1.50547E-06 | 0 | 6.14074E-06 | 0 | 0 | 1.66753E-06 |
| chr19:g.38997001T>A | 0.32459 | | | | | | | 0 | 0 | 0 | 0 | 0 | 0 |
| chr19:g.38997132G>A | 0.49988 | 3.98E-06 | 0 | 0 | 0 | 0 | 8.81E-06 | 1.9904E-06 | 0 | 0 | 0 | 0 | 4.40633E-06 |
| chr19:g.38997148C>G | 0.10000 | | | | | | | 0 | 0 | 0 | 0 | 0 | 0 |
| chr19:g.38997505C>T | 0.18771 | | | | | | | 0 | 0 | 0 | 0 | 0 | 0 |
| chr19:g.38998461C>T | 0.10000 | | | | | | | 0 | 0 | 0 | 0 | 0 | 0 |
| chr19:g.39002230G>A | 0.10000 | 0.000226348 | 0 | 0 | 0 | 0 | 9.29E-05 | 2.26348E-05 | 0 | 0 | 0 | 0 | 9.28908E-06 |
| chr19:g.39002919G>A | 0.10000 | | | | | | | 0 | 0 | 0 | 0 | 0 | 0 |
| chr19:g.39002961G>A | 0.94793 | 3.98E-06 | 0 | 3.27E-05 | 0 | 0 | 0 | 3.77105E-06 | 0 | 3.09619E-05 | 0 | 0 | 0 |
| chr19:g.39003007G>A | 0.10000 | 1.59E-05 | 0 | 0 | 0 | 0.000217533 | 0 | 1.59355E-06 | 0 | 0 | 0 | 2.17533E-05 | 0 |
| chr19:g.39005692C>T | 0.10000 | 7.96E-06 | 0 | 0 | 0 | 0 | 1.76E-05 | 7.95919E-07 | 0 | 0 | 0 | 0 | 1.75858E-06 |
| chr19:g.39006807A>G | 0.10000 | 0.000284364 | 0.000108425 | 0 | 0.000110019 | 0 | 0.000560617 | 2.84364E-05 | 1.08425E-05 | 0 | 1.10019E-05 | 0 | 5.60617E-05 |
| chr19:g.39006821T>C | 0.32459 | | | | | | | 0 | 0 | 0 | 0 | 0 | 0 |
| chr19:g.39006824G>A | 0.18771 | 1.07E-05 | 0 | 4.05E-05 | 0 | 0 | 1.27E-05 | 2.0022E-06 | 0 | 7.60083E-06 | 0 | 0 | 2.37572E-06 |
| chr19:g.39006848G>C | 0.18771 | | | | | | | 0 | 0 | 0 | 0 | 0 | 0 |
| chr19:g.39008071T>C | 0.10000 | 3.56E-05 | 0.000200965 | 0 | 5.65E-05 | 0.000150527 | 0 | 3.55905E-06 | 2.00965E-05 | 0 | 5.64525E-06 | 1.50527E-05 | 0 |
| chr19:g.39008110T>C | 0.18771 | | | | | | | 0 | 0 | 0 | 0 | 0 | 0 |
| chr19:g.39008161G>A | 0.10000 | 2.00E-05 | 0 | 0 | 8.68E-05 | 0 | 1.78E-05 | 2.00335E-06 | 0 | 0 | 8.68106E-06 | 0 | 1.7838E-06 |
| chr19:g.39008163T>A | 0.32459 | | | | | | | 0 | 0 | 0 | 0 | 0 | 0 |
| chr19:g.39008181G>A | 0.10000 | | | | | | | 0 | 0 | 0 | 0 | 0 | 0 |
| chr19:g.39009877C>T | 0.18771 | 6.73E-05 | 0 | 0 | 0 | 0 | 0.000132024 | 1.26315E-05 | 0 | 0 | 0 | 0 | 2.47823E-05 |
| chr19:g.39009878G>A | 0.18771 | 1.77E-05 | 0 | 0 | 8.47E-05 | 0 | 1.55E-05 | 3.32374E-06 | 0 | 0 | 1.58906E-05 | 0 | 2.91502E-06 |
| chr19:g.39009932G>A | 0.00590 | 0.000860488 | 0.000200819 | 0.000620591 | 0.000903751 | 0 | 0.001349926 | 5.07688E-06 | 1.18483E-06 | 3.66148E-06 | 5.33213E-06 | 0 | 7.96456E-06 |
| chr19:g.39009935A>G | 0.05072 | | | | | | | 0 | 0 | 0 | 0 | 0 | 0 |
| chr19:g.39010064C>A | 0.32459 | | | | | | | 0 | 0 | 0 | 0 | 0 | 0 |
| chr19:g.39010072A>T | 0.49988 | | | | | | | 0 | 0 | 0 | 0 | 0 | 0 |
| chr19:g.39010087A>G | 0.32459 | | | | | | | 0 | 0 | 0 | 0 | 0 | 0 |
| Chr19:g.39016072C>T | 0.10000 | | | | | | | 0 | 0 | 0 | 0 | 0 | 0 |
| chr19:g.39016132G>A | 0.00137 | 0.001639693 | 0.000604108 | 0.000653381 | 8.49E-05 | 0 | 0.003051059 | 2.2484E-06 | 8.28371E-07 | 8.95936E-07 | 1.16351E-07 | 0 | 4.18371E-06 |
| chr19:g.39018350G>C | 0.10000 | | | | | | | 0 | 0 | 0 | 0 | 0 | 0 |
| chr19:g.39019012G>T | 0.49988 | | | | | | | 0 | 0 | 0 | 0 | 0 | 0 |
| chr19:g.39019642G>C | 0.18771 | | | | | | | 0 | 0 | 0 | 0 | 0 | 0 |
| chr19:g.39019676G>T | 0.32459 | | | | | | | 0 | 0 | 0 | 0 | 0 | 0 |
| chr19:g.39019682C>T | 0.18771 | | | | | | | 0 | 0 | 0 | 0 | 0 | 0 |
| chr19:g.39019688C>T | 0.32459 | 4.02E-06 | 6.17E-05 | 0 | 0 | 0 | 0 | 1.30541E-06 | 2.00167E-05 | 0 | 0 | 0 | 0 |
| chr19:g.39025414C>T | 0.67519 | 1.20E-05 | 0 | 0 | 0 | 0 | 2.65E-05 | 8.10098E-06 | 0 | 0 | 0 | 0 | 1.79156E-05 |
| chr19:g.39025415G>A | 0.89991 | 1.78E-05 | 0 | 3.28E-05 | 0 | 0 | 1.56E-05 | 1.59948E-05 | 0 | 2.95556E-05 | 0 | 0 | 1.40158E-05 |
| chr19:g.39025837G>A | 0.67519 | | | | | | | 0 | 0 | 0 | 0 | 0 | 0 |
| chr19:g.39026638G>A | 0.10000 | 8.16E-05 | 0.000241313 | 3.29E-05 | 5.66E-05 | 0.000401526 | 3.11E-05 | 8.16489E-06 | 2.41313E-05 | 3.29164E-06 | 5.66027E-06 | 4.01526E-05 | 3.10709E-06 |
| chr19:g.39034005G>A | 0.89991 | 1.19E-05 | 0 | 0 | 0 | 0 | 2.64E-05 | 1.07349E-05 | 0 | 0 | 0 | 0 | 2.37306E-05 |
| chr19:g.39034020A>T | 0.49988 | | | | | | | 0 | 0 | 0 | 0 | 0 | 0 |
| chr19:g.39034045T>G | 0.10000 | | | | | | | 0 | 0 | 0 | 0 | 0 | 0 |
| chr19:g.39034191A>G | 0.02505 | 0.000848692 | 0.000160244 | 0.000620591 | 0.000903189 | 0 | 0.001332094 | 2.12597E-05 | 4.0141E-06 | 1.55458E-05 | 2.26249E-05 | 0 | 3.3369E-05 |
| chr19:g.39034206G>A | 0.10000 | | | | | | | 0 | 0 | 0 | 0 | 0 | 0 |
| chr19:g.39034450C>T | 0.67519 | | | | | | | 0 | 0 | 0 | 0 | 0 | 0 |
| chr19:g.39034456T>C | 0.67519 | | | | | | | 0 | 0 | 0 | 0 | 0 | 0 |
| chr19:g.39034461C>G | 0.67519 | | | | | | | 0 | 0 | 0 | 0 | 0 | 0 |
| chr19:g.39034472G>T | 0.99409 | | | | | | | 0 | 0 | 0 | 0 | 0 | 0 |
| chr19:g.39037100G>A | 0.10000 | 3.18E-05 | 6.15E-05 | 0 | 0 | 5.44E-05 | 4.40E-05 | 3.18155E-06 | 6.15385E-06 | 0 | 0 | 5.43656E-06 | 4.39584E-06 |
| chr19:g.39037136A>G | 0.10000 | 1.99E-05 | 0 | 0 | 0 | 0.000271828 | 0 | 1.98825E-06 | 0 | 0 | 0 | 2.71828E-05 | 0 |
| chr19:g.39038893A>T | 0.18771 | | | | | | | 0 | 0 | 0 | 0 | 0 | 0 |
| chr19:g.39038899C>T | 0.10000 | 1.19E-05 | 0 | 0 | 0 | 0 | 2.64E-05 | 1.19301E-06 | 0 | 0 | 0 | 0 | 2.63746E-06 |
| chr19:g.39038927C>A | 0.94793 | | | | | | | 0 | 0 | 0 | 0 | 0 | 0 |
| chr19:g.39039020C>T | 0.10000 | 4.77E-05 | 6.15E-05 | 6.53E-05 | 2.89E-05 | 0.000326193 | 1.76E-05 | 4.77164E-06 | 6.15157E-06 | 6.53253E-06 | 2.89084E-06 | 3.26193E-05 | 1.75799E-06 |
| chr19:g.39051780G>C | 0.01220 | | | | | | | 0 | 0 | 0 | 0 | 0 | 0 |
| chr19:g.39051825A>T | 0.18771 | 7.11E-06 | 0 | 0 | 0 | 0 | 1.57E-05 | 1.33443E-06 | 0 | 0 | 0 | 0 | 2.93967E-06 |
| chr19:g.39051853C>T | 0.10000 | | | | | | | 0 | 0 | 0 | 0 | 0 | 0 |
| chr19:g.39051868A>G | 0.49988 | 3.54E-05 | 0 | 6.53E-05 | 0 | 0 | 6.21E-05 | 1.76936E-05 | 0 | 3.26548E-05 | 0 | 0 | 3.10181E-05 |
| chr19:g.39051876C>A | 0.02505 | 2.39E-05 | 0 | 0 | 2.89E-05 | 0 | 3.52E-05 | 5.98528E-07 | 0 | 0 | 7.24365E-07 | 0 | 8.82508E-07 |
| chr19:g.39051883T>C | 0.49988 | | | | | | | 0 | 0 | 0 | 0 | 0 | 0 |
| chr19:g.39052002G>A | 0.49988 | | | | | | | 0 | 0 | 0 | 0 | 0 | 0 |
| chr19:g.39052003G>T | 0.67519 | | | | | | | 0 | 0 | 0 | 0 | 0 | 0 |
| chr19:g.39052023G>A | 0.10000 | 0.000327006 | 0.000200529 | 0 | 0.000141195 | 0 | 0.00063093 | 3.27006E-05 | 2.00529E-05 | 0 | 1.41195E-05 | 0 | 6.3093E-05 |
| chr19:g.39055663T>G | 0.67519 | | | | | | | 0 | 0 | 0 | 0 | 0 | 0 |
| chr19:g.39055674G>C | 0.94793 | | | | | | | 0 | 0 | 0 | 0 | 0 | 0 |
| chr19:g.39055674G>T | 0.94924 | | | | | | | 0 | 0 | 0 | 0 | 0 | 0 |
| chr19:g.39055822A>T | 0.10000 | | | | | | | 0 | 0 | 0 | 0 | 0 | 0 |
| chr19:g.39055858C>T | 0.00000 | 0.001615639 | 0.001628257 | 0 | 0 | 0 | 0.001853499 | 0 | 0 | 0 | 0 | 0 | 0 |
| chr19:g.39057618A>G | 0.10000 | 7.72E-05 | 0 | 7.44E-05 | 0 | 0 | 0.00014602 | 7.72414E-06 | 0 | 7.43716E-06 | 0 | 0 | 1.4602E-05 |
| chr19:g.39061259C>T | 0.18771 | 3.98E-06 | 0 | 0 | 0 | 0 | 8.79E-06 | 7.46403E-07 | 0 | 0 | 0 | 0 | 1.64999E-06 |
| chr19:g.39061260G>A | 0.32459 | 3.54E-05 | 4.01E-05 | 0 | 2.82E-05 | 0.000100241 | 4.64E-05 | 1.14746E-05 | 1.30013E-05 | 0 | 9.15938E-06 | 3.25371E-05 | 1.5075E-05 |
| chr19:g.39061289C>G | 0.10000 | | | | | | | 0 | 0 | 0 | 0 | 0 | 0 |
| chr19:g.39062672C>T | 0.10000 | 3.18E-05 | 0 | 0 | 0 | 0 | 6.97E-05 | 3.18377E-06 | 0 | 0 | 0 | 0 | 6.96778E-06 |
| chr19:g.39062825G>A | 0.49988 | | | | | | | 0 | 0 | 0 | 0 | 0 | 0 |
| chr19:g.39062830A>G | 0.67519 | 3.98E-06 | 0 | 0 | 0 | 0 | 8.79E-06 | 2.68487E-06 | 0 | 0 | 0 | 0 | 5.93501E-06 |
| chr19:g.39062846G>A | 0.18771 | 5.97E-05 | 0.000307579 | 0 | 2.89E-05 | 0.000380559 | 8.79E-06 | 1.11969E-05 | 5.77356E-05 | 0 | 5.42765E-06 | 7.14347E-05 | 1.65014E-06 |
| chr19:g.39062902T>C | 0.67519 | 3.99E-06 | 0 | 0 | 0 | 0 | 8.83E-06 | 2.69223E-06 | 0 | 0 | 0 | 0 | 5.96142E-06 |
| chr19:g.39062906_39062907delinsCT | 0.32459 | | | | | | | | | | | | |
| chr19:g.39063820C>T | 0.49988 | 1.77E-05 | 0 | 0 | 0 | 0 | 7.74E-06 | 8.83786E-06 | 0 | 0 | 0 | 0 | 3.87078E-06 |
| chr19:g.39063869T>C | 0.67519 | | | | | | | 0 | 0 | 0 | 0 | 0 | 0 |
| chr19:g.39063944C>T | 0.32459 | 4.27E-05 | 0 | 3.27E-05 | 2.83E-05 | 0 | 7.83E-06 | 1.38578E-05 | 0 | 1.0611E-05 | 9.1801E-06 | 0 | 2.54106E-06 |
| chr19:g.39066597G>A | 0.05072 | 0.00016978 | 8.02E-05 | 0 | 0 | 0 | 0.00032541 | 8.61127E-06 | 4.06736E-06 | 0 | 0 | 0 | 1.65048E-05 |
| chr19:g.39068571A>C | 0.18771 | | | | | | | 0 | 0 | 0 | 0 | 0 | 0 |
| chr19:g.39068582T>G | 0.32459 | | | | | | | 0 | 0 | 0 | 0 | 0 | 0 |
| chr19:g.39068586G>A | 0.67519 | | | | | | | 0 | 0 | 0 | 0 | 0 | 0 |
| chr19:g.39068594C>T | 0.94924 | | | | | | | 0 | 0 | 0 | 0 | 0 | 0 |
| chr19:g.39068595G>A | 0.98779 | 3.98E-06 | 0 | 0 | 0 | 0 | 8.79E-06 | 3.92913E-06 | 0 | 0 | 0 | 0 | 8.68738E-06 |

| | | | | | | | | | | | | | |
|---|---|---|---|---|---|---|---|---|---|---|---|---|---|
| chr19:g.39068655G>A | 0.10000 | 9.96E-05 | 0.000123411 | 3.27E-05 | 5.79E-05 | 0 | 0.000158546 | 9.95588E-06 | 1.23411E-05 | 3.26669E-06 | 5.78838E-06 | 0 | 1.58546E-05 |
| chr19:g.39068845G>T | 0.18771 | | | | | | | 0 | 0 | 0 | 0 | 0 | 0 |
| chr19:g.39070679_39070680delinsAA | 0.18771 | | | | | | | | | | | | |
| chr19:g.39070681C>A | 0.67519 | | | | | | | 0 | 0 | 0 | 0 | 0 | 0 |
| chr19:g.39070706A>T | 0.67519 | | | | | | | 0 | 0 | 0 | 0 | 0 | 0 |
| chr19:g.39070715G>A | 0.32459 | | | | | | | 0 | 0 | 0 | 0 | 0 | 0 |
| chr19:g.39070715G>T | 0.67519 | | | | | | | 0 | 0 | 0 | 0 | 0 | 0 |
| chr19:g.39070728T>C | 0.81214 | | | | | | | 0 | 0 | 0 | 0 | 0 | 0 |
| chr19:g.39070734C>T | 0.99985 | 3.98E-06 | 0 | 0 | 0 | 0 | 8.80E-06 | 3.97751E-06 | 0 | 0 | 0 | 0 | 8.7967E-06 |
| chr19:g.39070754C>T | 0.98779 | | | | | | | 0 | 0 | 0 | 0 | 0 | 0 |
| chr19:g.39070766C>G | 0.67519 | | | | | | | 0 | 0 | 0 | 0 | 0 | 0 |
| Chr19:g.39070767del | 0.02505 | | | | | | | | | | | | |
| chr19:g.39071010C>G | 0.89991 | | | | | | | 0 | 0 | 0 | 0 | 0 | 0 |
| chr19:g.39071022G>A | 0.49988 | 7.43E-05 | 0.000200385 | 9.80E-05 | 2.82E-05 | 5.01E-05 | 2.32E-05 | 3.71416E-05 | 0.000100168 | 4.89854E-05 | 1.41145E-05 | 2.50541E-05 | 1.16163E-05 |
| chr19:g.39071037G>C | 0.89991 | | | | | | | 0 | 0 | 0 | 0 | 0 | 0 |
| chr19:g.39071043G>A | 0.99715 | 1.77E-05 | 4.01E-05 | 0 | 2.82E-05 | 0 | 2.32E-05 | 1.76309E-05 | 3.99627E-05 | 0 | 2.81379E-05 | 0 | 2.3164E-05 |
| chr19:g.39071056C>T | 0.49988 | | | | | | | 0 | 0 | 0 | 0 | 0 | 0 |
| chr19:g.39071065C>G | 0.49988 | | | | | | | 0 | 0 | 0 | 0 | 0 | 0 |
| chr19:g.39071079C>T | 0.81200 | | | | | | | 0 | 0 | 0 | 0 | 0 | 0 |
| chr19:g.39071080G>A | 0.89991 | | | | | | | 0 | 0 | 0 | 0 | 0 | 0 |
| chr19:g.39071125A>G | 0.94924 | | | | | | | 0 | 0 | 0 | 0 | 0 | 0 |
| chr19:g.39071137T>C | 0.67519 | | | | | | | 0 | 0 | 0 | 0 | 0 | 0 |
| chr19:g.39075614G>A | 0.49988 | | | | | | | 0 | 0 | 0 | 0 | 0 | 0 |
| chr19:g.39075616G>A | 0.81214 | | | | | | | 0 | 0 | 0 | 0 | 0 | 0 |
| chr19:g.39075718A>G | 0.18771 | 3.19E-05 | 0 | 0 | 0 | 0 | 6.49E-05 | 5.98412E-06 | 0 | 0 | 0 | 0 | 1.21763E-05 |
| chr19:g.39075739G>A | 0.98779 | | | | | | | 0 | 0 | 0 | 0 | 0 | 0 |
| chr19:g.39076587T>C | 0.89991 | | | | | | | 0 | 0 | 0 | 0 | 0 | 0 |
| chr19:g.39076588C>G | 0.49988 | | | | | | | 0 | 0 | 0 | 0 | 0 | 0 |
| chr19:g.39076591C>A | 0.32459 | | | | | | | 0 | 0 | 0 | 0 | 0 | 0 |
| chr19:g.39076592G>A | 0.81214 | | | | | | | 0 | 0 | 0 | 0 | 0 | 0 |
| chr19:g.39076599G>T | 0.49988 | | | | | | | 0 | 0 | 0 | 0 | 0 | 0 |
| chr19:g.39076741T>A | 0.49988 | | | | | | | 0 | 0 | 0 | 0 | 0 | 0 |
| chr19:g.39076780C>T | 0.81214 | 2.78E-05 | 0 | 0 | 8.67E-05 | 0 | 3.52E-05 | 2.2614E-05 | 0 | 0 | 7.04453E-05 | 0 | 2.85708E-05 |
| chr19:g.39076830A>G | 0.32459 | | | | | | | 0 | 0 | 0 | 0 | 0 | 0 |
| chr19:g.39078002G>C | 0.32459 | | | | | | | 0 | 0 | 0 | 0 | 0 | 0 |
| chr19:g.39078003G>C | 0.49988 | | | | | | | 0 | 0 | 0 | 0 | 0 | 0 |

Supplemental Table 5

| | | gnomAD_All | gnomAD_AFR | gnomAD_SAS | gnomAD_AMR | gnomAD_EAS | gnomAD_NFE |
|---|---|---|---|---|---|---|---|
| **Sum of Frequencies * Posterior Probabilities** | Sum All | 0.00167 | 0.00161 | 0.00092 | 0.00111 | 0.00137 | 0.00209 |
| | Sum VUS/LP/P | 0.00159 | 0.00158 | 0.00084 | 0.00105 | 0.00137 | 0.00196 |
| | Sum P/LP | 0.00047 | 0.00036 | 0.00031 | 0.00008 | 0.00030 | 0.00063 |
| | Sum VUS | 0.00113 | 0.00122 | 0.00053 | 0.00097 | 0.00107 | 0.00134 |
| | No Criteria Met | 0.00021 | 0.00010 | 0.00008 | 0.00008 | 0.00014 | 0.00025 |
| | VUS/LP/P (PS4) | 0.00078 | 0.00054 | 0.00051 | 0.00053 | 0.00046 | 0.00105 |
| **Predicted Pathogenic Allele Frequency** | All | 1/599 | 1/623 | 1/1,092 | 1/905 | 1/729 | 1/480 |
| | VUS/LP/P | 1/628 | 1/632 | 1/1,193 | 1/948 | 1/730 | 1/510 |
| | P/LP | 1/2,149 | 1/2,810 | 1/3,227 | 1/11,979 | 1/3,298 | 1/1,604 |
| | VUS | 1/887 | 1/815 | 1/1,891 | 1/1,030 | 1/937 | 1/748 |
| | No Criteria Met | 1/4,827 | 1/9,815 | 1/12,066 | 1/12,972 | 1/6,913 | 1/3,958 |
| | VUS/LP/P (PS4) | 1/1,282 | 1/1,854 | 1/1,971 | 1/1,897 | 1/2,158 | 1/953 |
| **Predicted Pathogenic Allele Prevalence** | All | 1/300 | 1/312 | 1/546 | 1/453 | 1/365 | 1/240 |
| | VUS/LP/P | 1/314 | 1/316 | 1/597 | 1/474 | 1/365 | 1/255 |
| | P/LP | 1/1,075 | 1/1,405 | 1/1,614 | 1/5,990 | 1/1,649 | 1/802 |
| | VUS | 1/444 | 1/408 | 1/946 | 1/515 | 1/469 | 1/374 |
| | No Criteria Met | 1/2,414 | 1/4,908 | 1/6,033 | 1/6,486 | 1/3,457 | 1/1,979 |
| | VUS/LP/P (PS4) | 1/641 | 1/927 | 1/986 | 1/949 | 1/1,079 | 1/477 |
